# Discovery of 2-amide-3-methylester thiophenes that target SARS-CoV-2 Mac1 and repress coronavirus replication, validating Mac1 as an anti-viral target

**DOI:** 10.1101/2023.08.28.555062

**Authors:** Sarah Wazir, Tomi A. O. Parviainen, Jessica J. Pfannenstiel, Men Thi Hoai Duong, Daniel Cluff, Sven T. Sowa, Albert Galera-Prat, Dana Ferraris, Mirko M. Maksimainen, Anthony R. Fehr, Juha P. Heiskanen, Lari Lehtiö

## Abstract

The COVID-19 pandemic caused by severe acute respiratory syndrome coronavirus 2 (SARS-CoV-2) virus has made it clear that further development of antiviral therapies will be needed to combat additional SARS-CoV-2 variants or novel CoVs. Here, we describe small molecule inhibitors for SARS-CoV-2 Mac1, which counters ADP-ribosylation mediated innate immune responses. The compounds inhibiting Mac1 were discovered through high-throughput screening (HTS) using a protein FRET-based competition assay and the best hit compound had an IC_50_ of 14 µM. Three validated HTS hits have the same 2-amide-3-methylester thiophene scaffold and the scaffold was selected for structure-activity relationship (SAR) studies through commercial and synthesized analogs. We studied the compound binding mode in detail using X-ray crystallography and this allowed us to focus on specific features of the compound and design analogs. Compound **27** (MDOLL-0229) had an IC_50_ of 2.1 µM and was generally selective for CoV Mac1 proteins after profiling for activity against a panel of viral and human ADP-ribose binding proteins. The improved potency allowed testing of its effect on virus replication and indeed, **27** inhibited replication of both MHVa prototype CoV, and SARS-CoV-2. Furthermore, sequencing of a drug-resistant MHV identified mutations in Mac1, further demonstrating the specificity of **27.** Compound **27** is the first Mac1 targeted small molecule demonstrated to inhibit coronavirus replication in a cell model. This, together with its well-defined binding mode, makes **27** a good candidate for further hit/lead-optimization efforts.

## Introduction

SARS-CoV-2 is a highly pathogenic coronavirus and is the causative agent of COVID-19, which has caused major health and economic impacts worldwide ^1^. The virus primarily infects lung epithelial cells and uses cellular machinery for the translation of viral proteins and replication of viral RNA. Upon infection, the host innate immune response is activated by interferons (IFNs) that initiate the cellular antiviral defense systems to inhibit viral replication at most steps of the viral lifecycle ^2–4^. One such host cell defense mechanism that is induced by virus infection through the interferon (IFN) response is the post-translational ADP-ribosylation of human and or viral proteins which can inhibit virus replication in a variety of ways ^5^.

In human cells protein ADP-ribosylation is mainly executed by Diphtheria toxin-like PARP enzymes of the human ARTD family ^6^. IFN responsive PARPs include PARP7, PARP9, PARP10, PARP11, PARP12, PARP13 and PARP14 ^7–9^ and these proteins have been shown to inhibit the replication of a diverse panel of viruses ^10^. Several positive-sense RNA viruses, including coronaviruses (CoVs), alphaviruses, and Hepatitis E virus counter this immune response by encoding for macrodomains, which have ADP-ribose binding and ADP-ribosylhydrolase activity. SARS-CoV-2 non-structural protein 3 (nsp3) contains three macrodomains in tandem, but only the first one (Mac1) possesses hydrolase activity ^11^. Both the SARS-CoV and SARS-CoV-2 Mac1 proteins have been shown to enzymatically reverse mono-ADP-ribosylation, suppress host IFN production, and promote viral pathogenesis ^12–14^. Therefore, SARS-CoV-2 Mac1 is an intriguing target for drug discovery and small molecule inhibitors might offer novel therapeutics to combat COVID-19.

Initial compounds targeting virus macrodomains have been described for alphaviruses ^15,16^ and coronaviruses ^17–20^. These efforts have used both screening of compound libraries ^18,21,22^, *in crystallo* fragment screening to discover early small inhibitors ^19^ and further development to produce more potent compounds ^17,20^. The majority of the compounds have reached low-mid µM potency, though a few have reach sub-µM IC_50_’s, and in general they have lacked the properties allowing cell permeability and subsequent validation of Mac1 as a pharmacological target.

Using a recently developed robust protein FRET based assay ^21,23^ we performed high-throughput screening of compounds to identify novel inhibitors of SARS-CoV-2 Mac1. We identified three hit compounds that all contained a 2-amide-3-methylester thiophene scaffold. The initial hit compounds showed 14 –110 µM potencies and stabilized Mac1 in a direct binding assay. This encouraged us to start a hit optimization campaign with an aim to reach low µM potency allowing pharmacological testing of Mac1 inhibition. The synthesis was guided by a co-crystal structure with Mac1 and allowed us to define a SAR. The structure-based optimizations improved IC_50_ values of the analogues to 2.1 µM after modifications of the 2-amide-3-methylester thiophene core. Both the hit compound and the optimized compound show clear selectivity towards SARS-CoV-2 over both other virus macrodomains and against all human macrodomains. The optimized compound also inhibited coronavirus replication in cell culture without any effects on metabolic activity of the cells. This knowledge provides a basis for the lead optimization efforts and strengthens the view that Mac1 could be a promising target to treat coronavirus infections.

## Results

### Screening and hit validation

Earlier we established a FRET-based ADP-ribosylation binding assay which was validated for the screening of compound libraries for SARS-CoV-2 Mac1 ^21^ (Table S1). Here, the assay was used to screen a larger compound library for SARS-CoV-2 Mac1 to obtain initial hit molecules. The screening of 30 000 compounds resulted in 74 hit compounds that inhibited >24% at 30 µM concentration, which is defined by a robust 5σ from the mean of all the compounds not filtered out due to excessive fluorescence (Fig. 1A). The hit compounds were then evaluated with a counter screening method to filter out false positives that is based on the same FRET readout testing for the inhibition of tankyrase 2 ARC4 peptide binding ^24^. In total, 66 compounds were identified which also inhibited TNKS2 ARC4 indicating that these were unselective protein inhibitors at the used concentration (Fig. 1B). One of the hits that did not inhibit TNKS2 ARC4 was additionally excluded at this stage as the difference in the inhibition with the two excitation wavelengths was large (>40%). The difference indicated that the compound would interfere with the FRET signal rather than with protein-protein interaction. In contrast to the fluorescent proteins, small molecules have sharper excitation and emission peaks while the inhibition of the FRET signal resulting from proximity of YFP and CFP is expected to be similar. Further, dose-response and thermal shifts were measured three times for each of the 8 hits. These results revealed 6-{[3-(methoxycarbonyl)-5,6,7,8-tetrahydro-4H-cyclohepta[b]thiophen-2-yl]carbamoyl}cyclohex-3-ene-1-carboxylic acid (**1**) as the best hit compound based on a positive shift in the melting temperature (1.3 ± 0.25 °C ΔTm at 100 μM concentration) and IC_50_ value of 14 μM (Fig. 1C, Table S2). We did not find previous publications or activity data for **1** and intriguingly two other hit compounds **2** and **3** have lower potency but share the same core structure with **1** (Fig. 1C, Fig. S1). All these three compounds show direct binding to Mac1 in the thermal shift assay (Fig. S2) and therefore the validation results support that the 2-amide-3-methylester thiophene scaffold would be a good starting point for hit optimization. An overall step-by-step view of the screening and validation is shown in Fig. 1D.

**Fig. 1.**
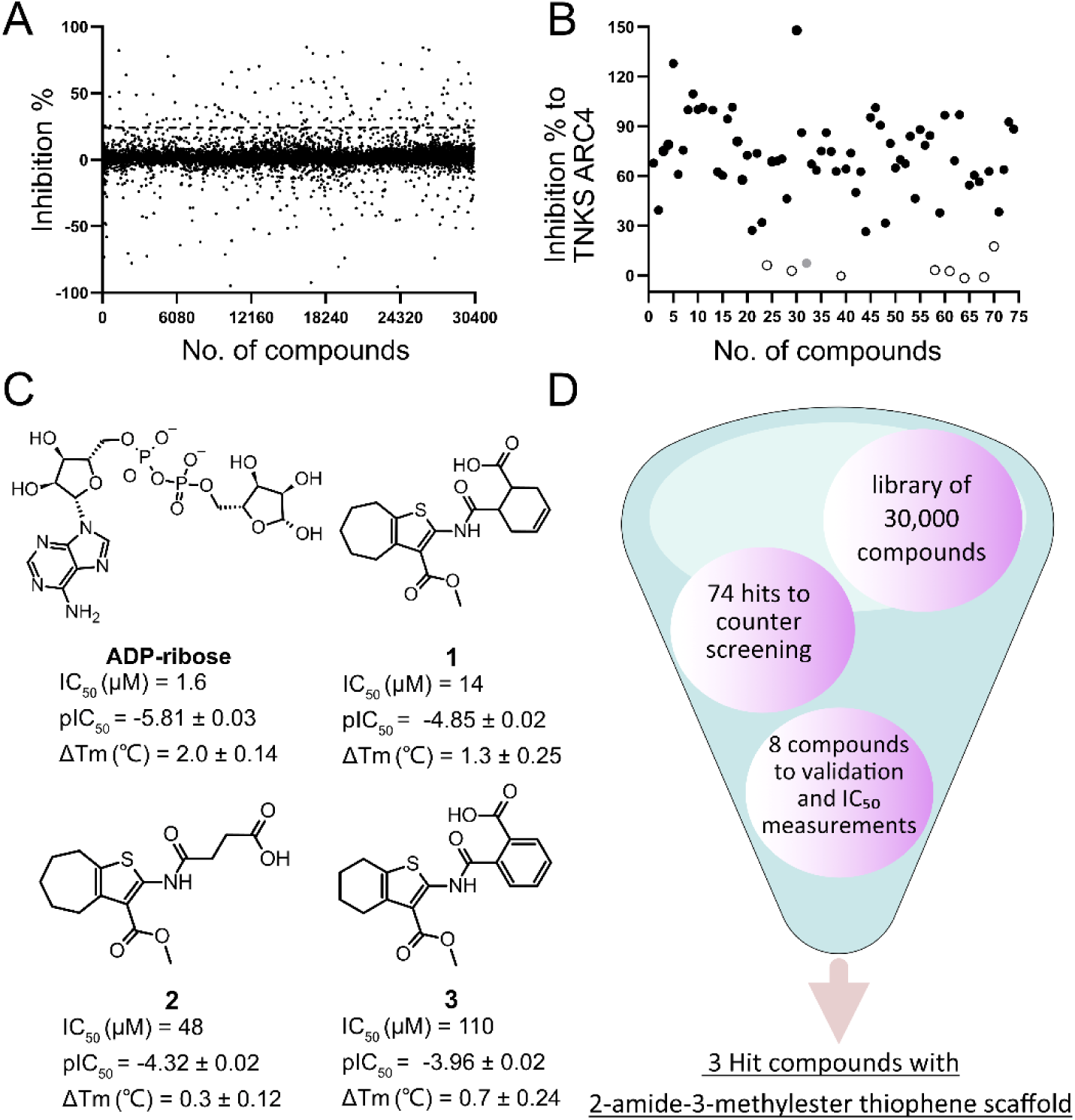
High-throughput compound screening to find SARS-CoV-2 Mac1 inhibitors. A) Normalized screening data for the compounds at 30 µM concentration. The hit limit is indicated with dashed lines was set to 24% 5σ from the mean of all data points. B) Counter screening of the initial hit compounds with TNKS ARC4. Selected compounds are highlighted in open circles with black outline. The compound highlighted in grey was identified as a false positive. C) Three screening hits with 2-amide-3-methylester thiophene scaffold and ADP-ribose control along with the measured IC_50_ values and thermal shifts along with standard deviations of three replicates. D) Overview of the screening process.

### Profiling for selectivity

In order to assess whether **1** would be a general macrodomain inhibitor or whether it would have the required selectivity to virus macrodomains, we profiled it against a wide panel consisting of all human as well as selected virus macrodomains. Macrodomains have a conserved structure and they bind and/or hydrolyse ADP-ribosyl groups and therefore it is plausible that discovered inhibitors would also have off-target effects. We additionally included ARH3, which is a human ADP-ribosylhydrolase with a different fold ^25^. Compound **1** was found to be very selective towards SARS-CoV-2 Mac1 among viral macrodomains as it did not show effective inhibition towards any other tested viral macrodomain, even at 100 μM concentration (Fig. 2). It was also selective towards Mac1 over all human macrodomains. The only affected human macrodomains were PARP9 MD1 and ALC1, and even then it only showed inhibition at 100 μM (≥ 70% PARP9, ≥ 50 % ALC1). Potency measurements confirmed that **1** shows 5-fold selectivity for SARS-CoV-2 Mac1 (IC_50_ = 14 μM) over PARP9 MD1 which has a modest IC_50_ of 64 μM (pIC_50_ ± SEM = -4.19 ± 0.07). It inhibits ALC1 with an IC_50_ of 6.8 μM (pIC_50_ ± SEM = -5.12 ± 0.04) but the maximum inhibition reached only 58% even with 1 mM concentration. ALC1 binds poly-ADP-ribose (PAR) and not mono-ADP-ribose (MAR) ^26,27^ and therefore **1** may bind only to one subsite in the protein and it would still be able to interact with PAR. The profiling further convinced us of the potential of optimizing 2-amide-3-methylester thiophene especially as SARS-CoV-2 Mac1 inhibitor.

**Fig. 2.**
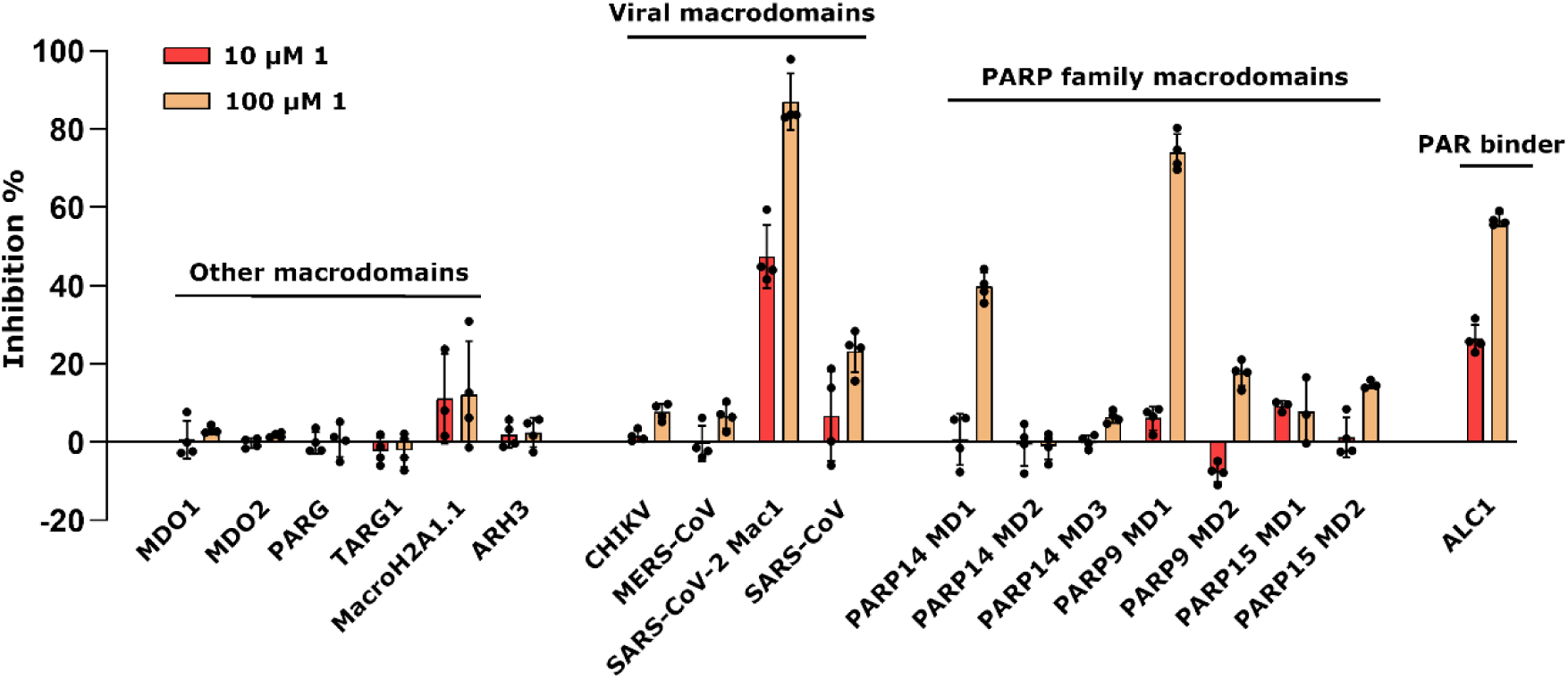
Inhibition profile of **1** against viral and human ADP-ribosyl readers and erasers. Inhibition % was calculated from rFRET signals of CFP-fused binders with YFP-GAP(MAR) and for ALC1 with YFP-GAP(PAR). Measurements were taken in quadruplicates (n = 4). Individual data points are shown.

### Crystal structure comparison of SARS-CoV-2 Mac1 in complex with 1

To allow efficient optimization of the compound we solved a co-crystal structure of **1** in complex with SARS-CoV-2 Mac1 (PDB: 8TV6). The structure was solved in space group P1 with two copies of Mac1 in the asymmetric unit. The structure was refined at 1.74 Å resolution and data collection and refinement statistics are shown in Table S2. The model built to the electron density included residues V3-F168 for chain A and V3-L169 for chain B. Compound **1** was only observed in subunit B and it binds to the adenosine site of the binding pocket of SARS-CoV-2 Mac1 (Fig. 3A). Comparison with the ADPr-complex structure revealed that the aliphatic seven-membered ring attached to a five-membered heteroaromatic thiophene unit forms hydrophobic interactions with Phe156 (Fig. 3B & 3C). The carbonyl oxygen of the methyl-ester group also forms a hydrogen bond with the backbone of Ile23, which is also one of the conserved interactions for ADPr. In addition to these, the negatively charged carboxylate group of the compound forms hydrogen bonds with the backbone amides of Phe156 and Asp157. Two water molecules were identified that mediate interactions between ligand and the protein main chain (W1-2, Fig. 3A). Recently, Correy and co-workers reported a detailed study on the SARS-CoV-2 Mac1 and ADPr bound structures as well as small compound fragments analyzing water networks aiding compound binding (Correy et al., 2022). We observed that the binding of **1** replaces a conserved water molecule (W1’), which bridges oxygen of the ribose and nitrogen of the adenine of ADPr (Fig. 3B).

Compound **1** is more potent towards SARS-CoV-2 Mac1 in comparison to **2** and **3**. Since all the three compounds share the same core scaffold it is likely that they share the same binding mode as the key features of **1** like carboxyl and methyl ester groups, and amide linkage are present in all (Fig. 1). The compounds also contain either a seven-membered (**1** and **2**) or a six-membered (**3**) aliphatic ring fused with a thiophene ring. The larger structural differences between the hit compounds occur at the linker between the carboxyl group and the central amide linkage, but it also explains the better affinity of **1**. Compound **1** contains a relatively rigid and sterically demanding cyclohexenyl group and when the linker is changed to a more flexible alkyl in **2** this is tolerated, but the potency drops to 3.5-fold (IC_50_ = 48 μM). On the other hand, the phenyl ring in the same position makes **3** more rigid but could also prevent efficient hydrogen bonding of the carboxyl group with the backbone amides, which could be a reason for its 7-fold drop in the potency (IC_50_ = 110 μM).

**Fig. 3.**
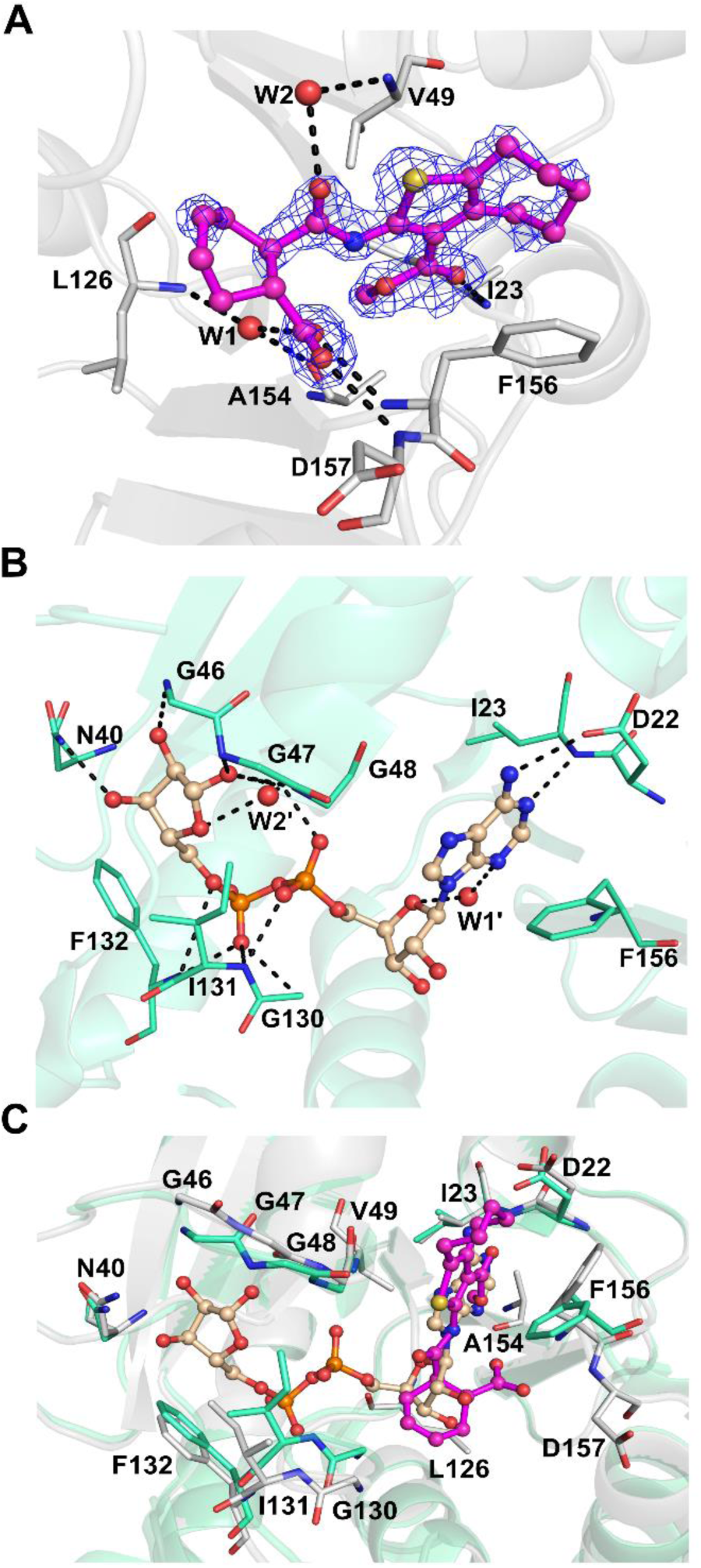
Comparison of SARS-CoV-2 Mac1 crystal structures: A) Co-crystal structure of **1** with SARS-CoV-2 Mac1 (PDB: 8TV6). The sigma-A weighted 2Fo-Fc electron density maps are colored in blue and contoured at 1.0 σ. B) SARS-CoV-2 Mac1 in complex with ADPr (PDB: 6WOJ^13^). C) Comparison of ADPr and **1** binding modes. ADPr and **1** are shown in ball and stick model. Hydrogen bonds are drawn as dashed lines. Water molecules are labelled with W.

### Resynthesis of 1, 3, and cyclic analogs

In order to strengthen the observations based on the hit compounds we decided to establish a synthesis method to be used later to produce analogs solely based on the original hit molecules. The central scaffold of **1** consists of 2-amide-3-methylester thiophene and in the first phase of synthesis we re-synthesized the best hit compounds **1** and **3**, annotated as **1r** and **3r** where r indicates a resynthesized compound. In addition, two novel compounds **10** and **11** with structures that differed from **1r** and **3r** in the size of the fused aliphatic ring systems, were synthesized in order to determine which combination of ring structures would have the most beneficial effect.

We recognized that the central scaffold containing methyl 2-amino-3-carboxythiophene with a fused ring system can be efficiently constructed via Gewald multicomponent reaction (Scheme 1). For the 2-amino-3-methyl ester thiophene fused with six-membered carbon ring (**9b**), a method described by Dang et al. was successfully utilized ^28^. For the cycloheptanyl counterpart (**9a**), a microwave-assisted synthesis method was used. In order to form the target compounds containing an amide linkage, cyclic anhydrides (phthalic anhydride and *cis*-1,2,3,6-tetrahydrophthalic anhydride) were selected as reagents for the next step. The most successful method for preparing the target compounds was to mix the solid starting materials in a reaction tube under argon and to heat the mixture overnight, ca. 16 h. Appropriate reaction temperatures for reactions were determined at the point in which melting occurred in the reaction mixture. Used temperatures varied between 90–130 °C, depending on the melting points of starting materials.

**Scheme 1.**
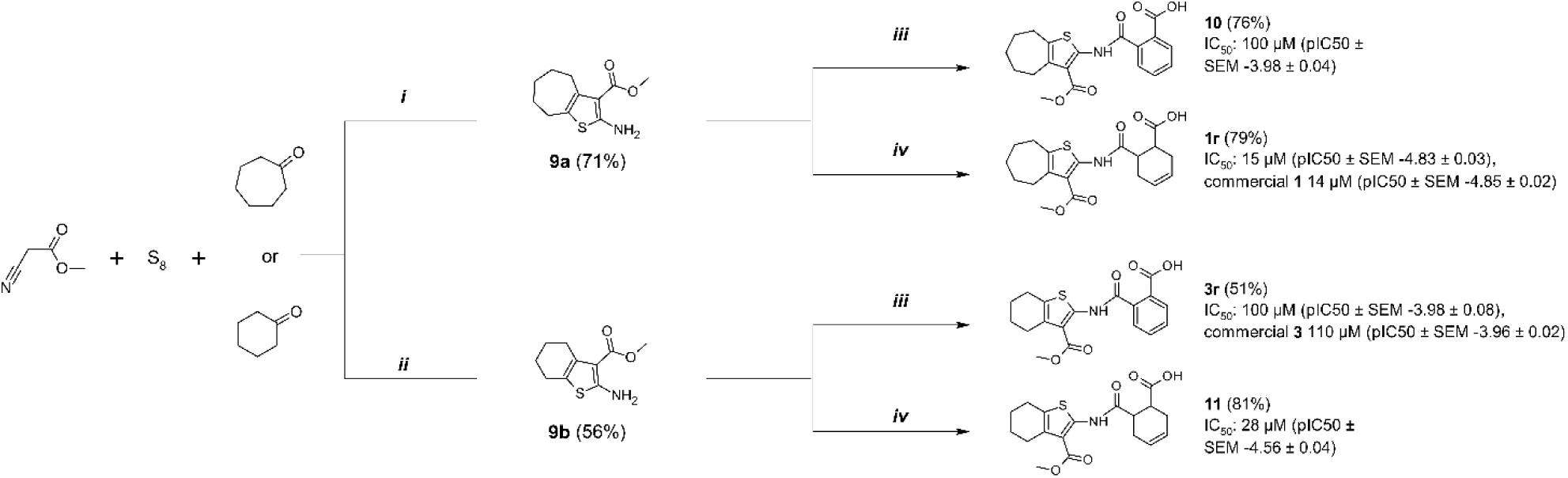
Resynthesis of screening hits and ring analogs with determined IC_50_ (pIC_50_ ± SEM) values. Reagents and conditions: (i) pyrrolidine, DMF, MW 60 °C, 30 min (ii) diethylamine, MeOH, rt, 25 h (iii) phthalic anhydride, Ar, heating, 16 h (iv) *cis*-1,2,3,6-tetrahydrophthalic anhydride, Ar, heating, 16 h.

We confirmed that the results for the re-synthesized compounds **1r** and **3r** correlated well with the initial IC_50_ values of the original hits (Scheme 1). The first two analogs were synthesized by keeping compounds **1** and **3** in mind. It is clear from these analogs that cyclohexenyl unit (**11** and **1r**) is preferred over phenyl structure (**10** and **3r**). Moreover, the combination of the bigger aliphatic fused cyclic structure with the cyclohexenyl moiety in **1r** results in lower IC_50_ thus confirming that the original hit compound had the best combination of these features.

### Exploration of the chemical space using commercial compounds

Based on early results, we evaluated a collection of commercially available analogs of **1** with single or in some cases double substitution to the basic scaffold. We focused on the substitutions on two main sites as this would make it possible to pinpoint compound features leading to improved potency and selectivity (Table 1). The biochemical assay revealed that the replacement of connecting cyclic system between amide and carboxyl groups with bicyclo[2.2.2]octane (**12** and **13**) completely abolished the inhibitory effect of the compounds, while norbornane (**14**) significantly reduces the potency compared to the parent compound **1**. Substitution of cyclohexenyl unit of compound **1** with its saturated counterpart cyclohexanyl unit improved the inhibition for **15** (IC_50_ = 12 µM). With cyclohexanyl moiety connecting the amide and carboxyl functionalities, the fused cyclopentanyl (**16**, IC_50_ = 22 µM) or cyclohexanyl (**17**, IC_50_ **=** 19 µM) rings with thiophene are tolerated, but their potency is less than that of **15**, a compound with fused cycloheptanyl ring. Use of ethyl cyclohexanyl as the fused aliphatic part with thiophene (**18**) almost completely abolished the inhibitory effect, but instead, the introduction of *tert*-butyl substituted cyclohexanyl ring (**19**) fused with the thiophene depicted an IC_50_ of 17 µM. Increasing the size of the ring system attached to the thiophene had a positive effect on the inhibition activity of the compounds. Indeed, compound with an eight–membered fused aliphatic ring (**20**) resulted in an IC_50_ of 7.8 µM, even lower than the base compound **1**.

**Table 1.**
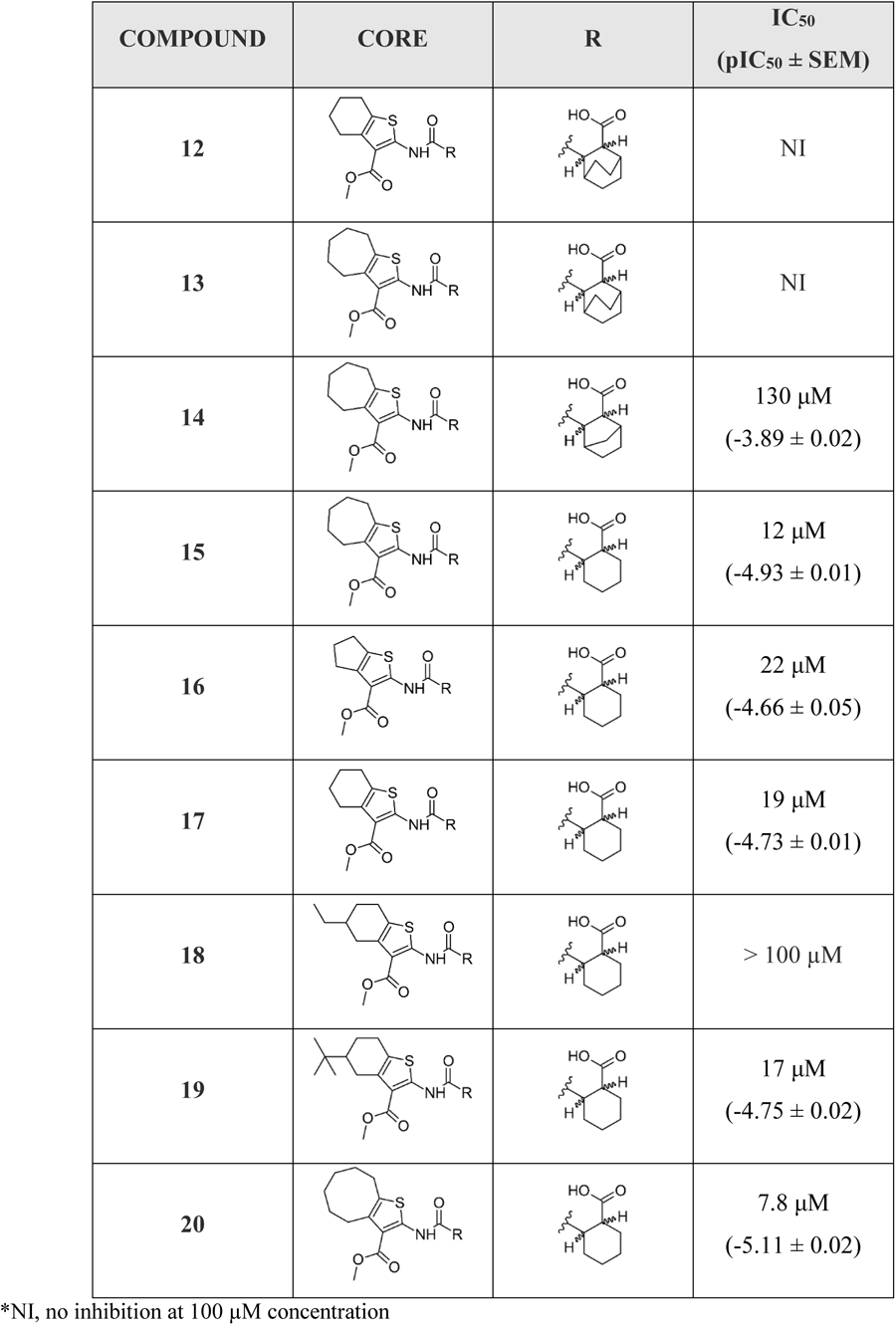
Commercial structural analogs of **1**. IC_50_ (pIC_50_ ± SEM, n=3) values are reported.

| COMPOUND | CORE | R | IC<sub>50</sub><br>(pIC<sub>50</sub> ± SEM) |
| --- | --- | --- | --- |
| 12 |  |  | NI |
| 13 |  |  | NI |
| 14 |  |  | 130 μM<br>(-3.89 ± 0.02) |
| 15 |  |  | 12 μM<br>(-4.93 ± 0.01) |
| 16 |  |  | 22 μM<br>(-4.66 ± 0.05) |
| 17 |  |  | 19 μM<br>(-4.73 ± 0.01) |
| 18 |  |  | > 100 μM |
| 19 |  |  | 17 μM<br>(-4.75 ± 0.02) |
| 20 |  |  | 7.8 μM<br>(-5.11 ± 0.02) |

\*NI, no inhibition at 100 μM concentration

### Synthesis of inhibitor analogs

In addition to the commercial analogs, a collection of compounds were prepared synthetically to allow more systematic and designed SAR studies. As a microwave assisted Gewald reaction turned out to be a good starting point, intermediate 2-aminothiophene compounds with different sizes of fused aliphatic rings and substituents at position 3 could be prepared with varying starting materials for the Gewald reaction. For example, when malononitrile was utilized as an activated nitrile starting material instead of methyl cyanoacetate, a nitrile group was formed in the 3-position of the thiophene unit (**9c**) (Scheme 2). For compound **9d** with the methyl ketone group in the 3-position, a different method originally reported by Abdelwahab et al. was noted to be the most successful ^29^. These 2-aminothiophenes with different sized fused aliphatic ring structures and different substituents at the 3-position of the thiophene were then used as starting materials with various cyclic anhydride compounds in the syntheses of structural analogs of compound **1**. The final products **21**, **22**, **15r**, and **25** –**27** were prepared in a similar manner as described for compound **1r** in Scheme 1. Compounds **1r** and **27** were further modified with an esterification reaction to afford compounds **23** and **28**, respectively. Compound **24** was prepared through hydrolysis of methyl ester functionality in the 3-position of thiophene unit of compound **15r**.

**Scheme 2.**
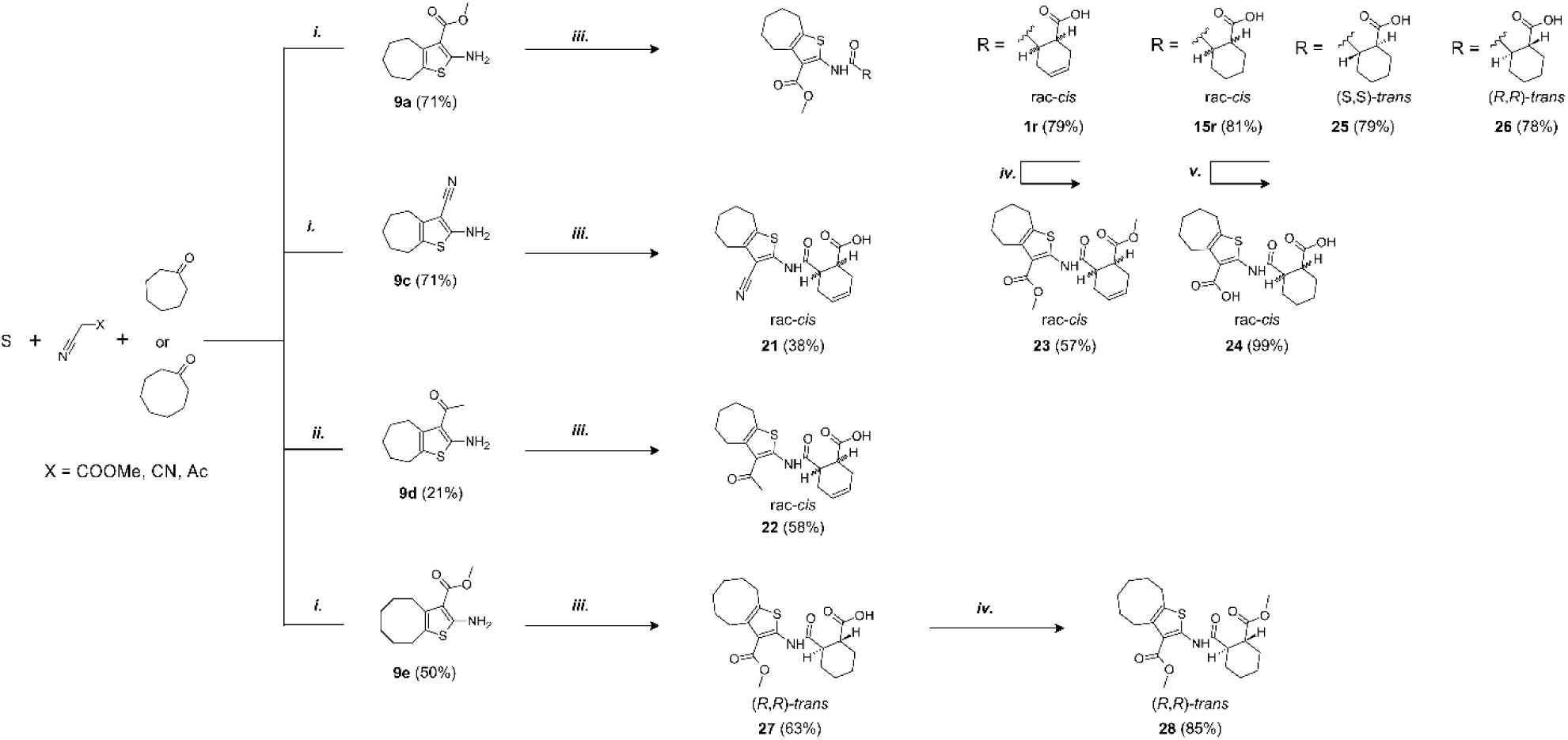
Synthesis of compounds **9**(**a**, **c** – **e**), **15r** and **21** –**28**. Reagents and conditions: (i) DMF, pyrrolidine, MW 60 °C, 30 min (General method A) (ii) MeOH, piperidine, 55 °C, 21 h (iii) corresponding anhydride, Ar, heating, 16 h (General method B) (iv) MeOH, conc H_2_SO_4_, reflux, 2 – 16 h (v) NaOH (aq), 90 °C, 2 h.

### Structure-activity relationship analysis

As seen from the SARS-CoV-2 Mac1 and compound **1** complex structure, the methyl ester group could be changed to improve suboptimal interactions made between its carbonyl oxygen and Ile23. Therefore, we considered replacing the methyl ester group with nitrile (**21**), methyl ketone (**22**), and amide groups. An amide modification for 3-position of the thiophene unit was done but it was disqualified from further studies due to instability issues. In this moiety, maintaining the hydrogen bond acceptor is likely important as there is a hydrogen bond formed with Ile23 in **1** co-crystal structure (Fig. 3A). However, both prepared analogs were inactive and therefore we decided to continue at this stage to explore other parts of the molecule. We synthesized a dimethyl ester derivative (**23**) to remove the ability of **1** to form a negative charge and to maintain the potential to form the hydrogen bonds to the backbone amides as seen in the co-crystal structure of **1**. This modification improved the compound potency slightly (IC_50_ = 11 µM).

The hit compound **1** is a racemic mixture of two enantiomers, namely (S,R) and (R,S), with *cis* configuration in the cyclic structure originated from the final reaction step with an anhydride. While it is possible that all isomers could inhibit Mac1 it is important to confirm the specific stereoisomers of the compound that are the most potent. For this we took guidance from the commercial analogs (Table 1) and decided to continue with cycloalkyl analogs that showed improvement in IC_50_ values. Therefore compound **15** was re-synthesized along with new compounds also with defined stereochemistry (Table 2). Surprisingly, our IC_50_ results were better for the resynthesized compound **15r** (rac-*cis*) in comparison to the commercial analog **15** (6.5 µM vs. 12 µM). From compound **15r**, an analog **24** with a carboxylic acid functionality was prepared with hydrolysis of the methyl ester at the 3-position of thiophene unit, but this was determined to be without inhibitory activity similarly to other ester functionality modifications (**21** & **22**). Compound **25**, the (S,S)-*trans* enantiomer was inactive, but instead, compounds **26** (an (R,R)-*trans* enantiomer) and **27** (an (R,R)-*trans* enantiomer) displayed excellent improvement in IC_50_ of 2.7 µM and 2.1 µM, respectively. Based on these results, we concluded that larger hydrophobic substituents fused with thiophene unit as well as (R,R)-*trans* absolute configuration in the cyclohexyl moiety may improve the potency against SARS-CoV-2 Mac1. As we had seen an improvement of a dimethyl ester **23** over carboxylic acid **1** we also prepared a similar analog of the best compound. Surprisingly, compound **28** was completely inactive. It could be possible that **23**, which is a racemic mixture, allows ester to bind, whereas for the defined R,R-*trans* form of **27** the carboxylate may be more strictly required allowing hydrogen bonding with main chain amides. Compound **23**, however, indicates that it could be possible to remove the negative charge by suitable modification in the future, which could be preferential for cell permeability. The most promising compound in the series would be **27**, which possessed the single digit micromolar potency and demonstrated a thermal shift of 3.2 ± 0.064 °C (Fig. S2). Despite the carboxylic acid, **27** is reasonably hydrophobic with a cLogD at pH 7.4 of 2.55.

**Table 2.**
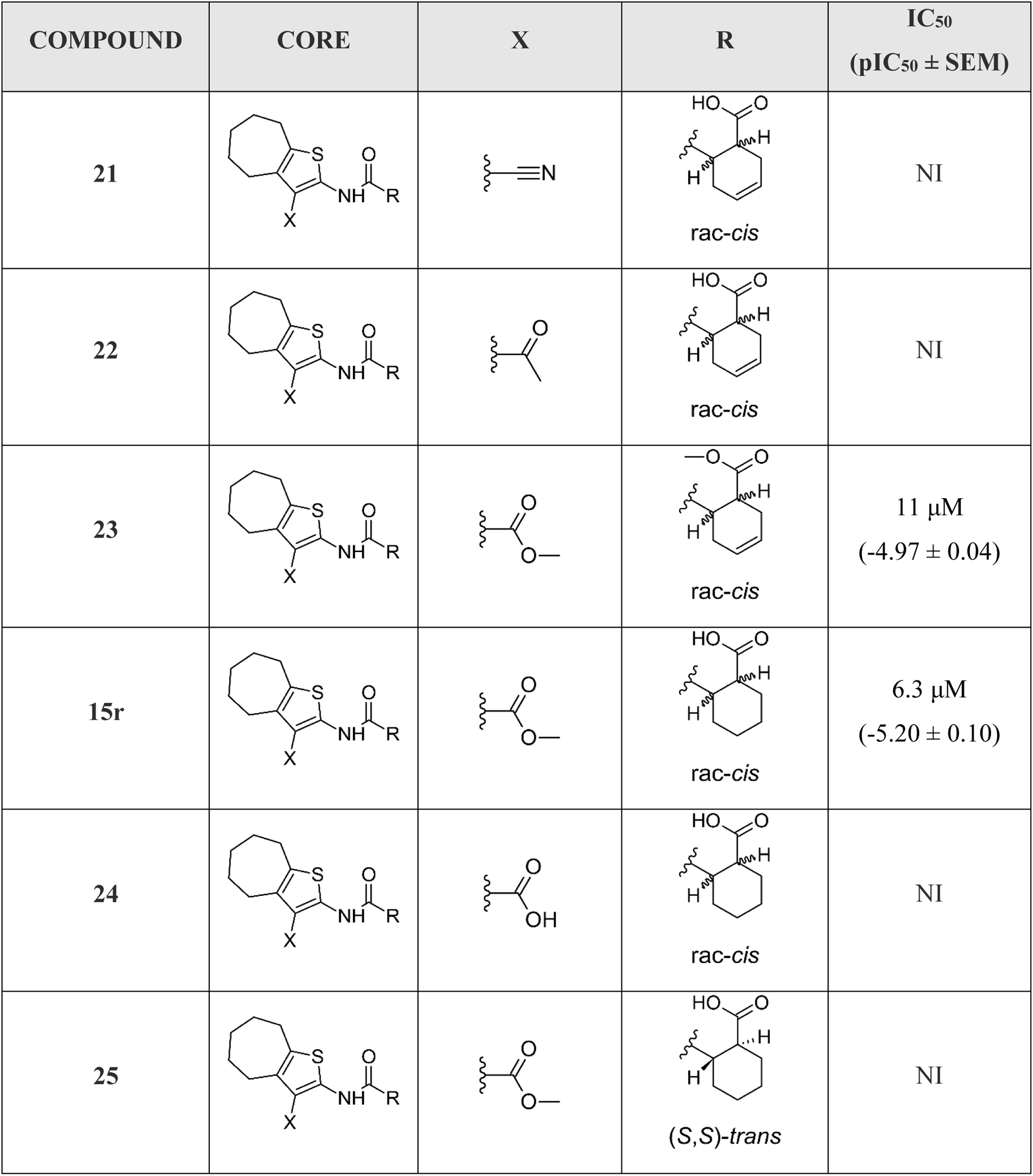
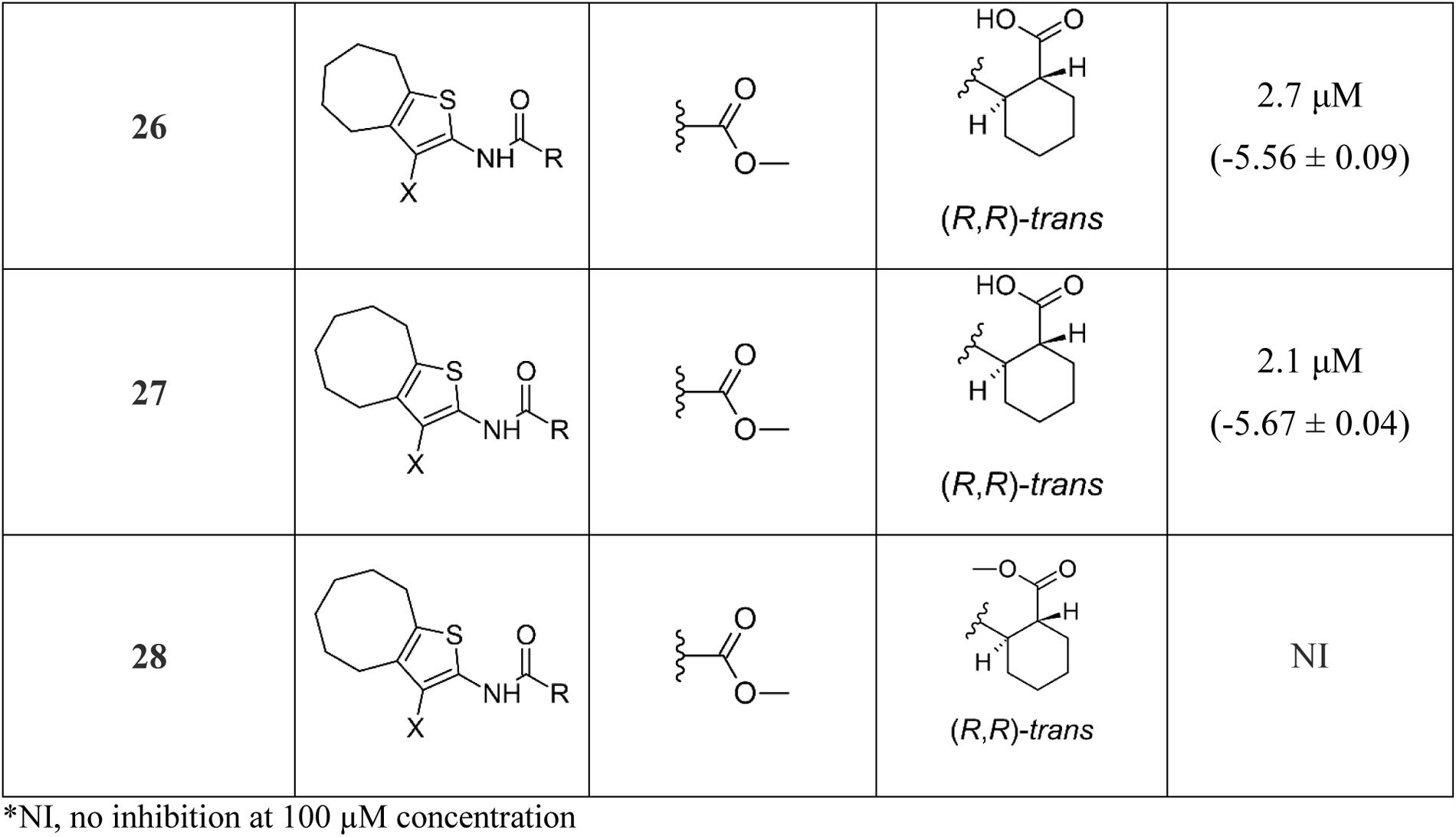
Synthesized structural analogs of **1** with IC_50_ (pIC_50_ ± SEM, n=3) values.

### Crystal structure of SARS-CoV-2 Mac1 in complex with 27

We solved a co-crystal structure of **27** with Mac1 (PDB: 8TV7) to study the binding mode in detail. Co-crystallization resulted in another crystal form in higher P2_1_2_1_2_1_ space group with one protein molecule in the active site. The structure was refined to 1.50 Å resolution and the electron density was good for the entire ligand (Fig. 4A). The improved electron density over the co-crystal structure of **1**, is likely due to the better potency and defined stereochemistry. The direct interactions with the protein are similar for **27** as they were for **1**, including hydrogen bonds with the backbone amides of Asp154 and F156 by the carboxylate, hydrogen bond with Ile23 by the methyl ester carbonyl and hydrophobic interaction with Phe156. The latter interaction is enhanced by the larger 8-membered fused aliphatic ring of **27** and explains also the improvement in potency. Phe156 orientation appears to be subsequently slightly changed (Fig. 4B). The methyl group of the methyl ester is well defined in the electron density and oriented towards a hydrophobic pocket formed by Ile23, Pro15 and Ala155. The conserved water molecule (W1 ̎) binding to carboxylate is present, but the second water molecule (W2 ̎) interacting with the amide carbonyl is moved in comparison to co-crystal structure of **1** and it is now bridging the interaction of **27** and backbone amide of Ile132. The overall co-crystal structure defines the binding mode of **27** in detail and will allow further structure-based design of analogs with improved properties.

**Fig. 4.**
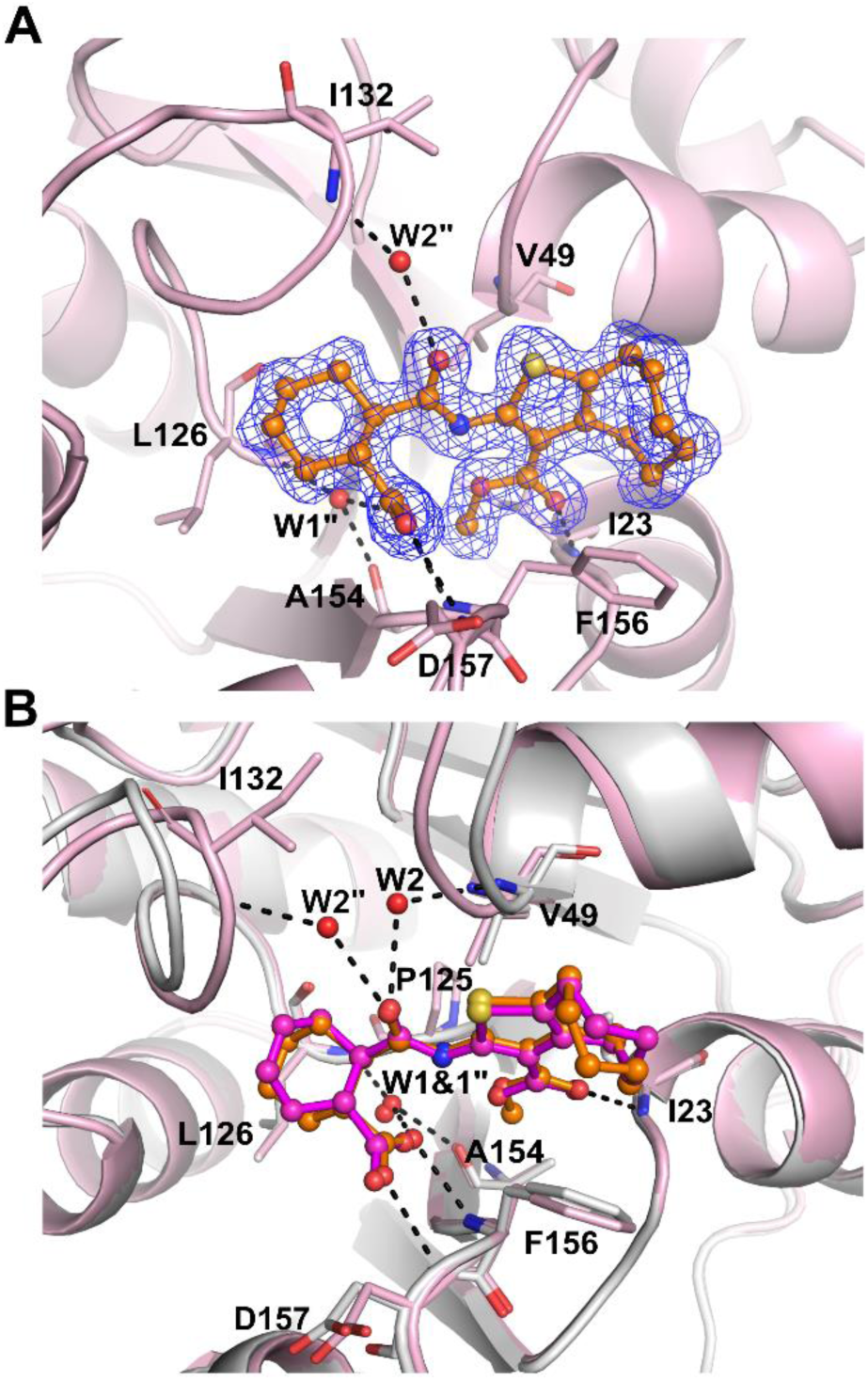
Co-crystal structure of SARS-CoV-2 Mac1 and **27** (PDB: 8TV7). A) The sigma-A weighted 2Fo-Fc electron density map is contoured at 1.0 σ. B) Comparison of **1** and **27** binding modes. **27** (Orange) and **1** (Magenta) are shown in ball and stick model. Hydrogen bonds are drawn as dashed lines. Water molecules are labelled with W.

### Selectivity towards SARS-CoV-2 Mac1

To determine if these modifications improved selectivity, **27** was also profiled against viral and human macrodomains as described above for **1** (Fig. 5A). It did not show effective inhibition against members of human macrodomains except for ALC1 (≥ 50 %) at both 10 μM and 100 μM. In addition to inhibiting of SARS-CoV-2 Mac1 at both tested concentrations it also inhibited macrodomains from SARS-CoV (64%, IC_50_ 20 μM, (pIC_50_ ± SEM = -4.71 ± 0.07)) and MERS-CoV (43%, IC_50_ >100 µM). The selectivity towards ALC1 was improved although the potency towards it was higher in comparison to **1** with an IC_50_ of 3.0 µM (pIC_50_ ± SEM = -5.52 ± 0.01) but again only 63% inhibition was reached at 1 mM concentration. To confirm that the **27** effectively inhibited also the ADP-ribosylhydrolysis activity we tested this with dot-blot using a kinase, SRPK2, which was mono-ADP-ribosylated by PARP10 (Fig. 5B). The inhibitor inhibited the enzymatic activity with similar efficiency as the free ADP-ribose, which in agreement with the similar IC_50_ values measured in FRET assay (2.1 µM and 1.6 µM, respectively; Fig. 1). The efficiently and selectivity profile indicates that the compound scaffold is suitable against viral macrodomains and can lead to new biological understanding of coronavirus infections and the possible future outbreaks of its variants.

**Fig. 5.**
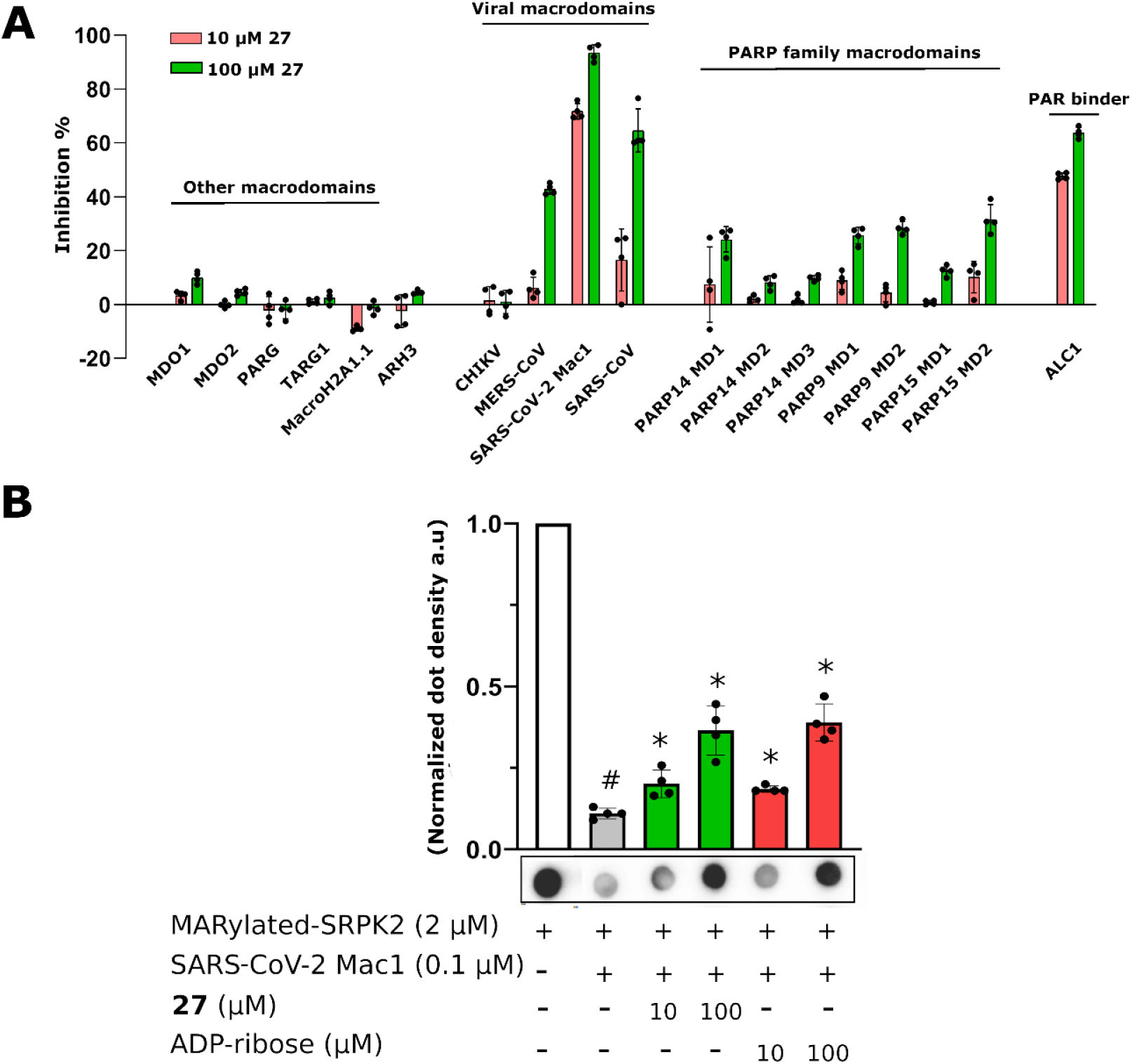
Selectivity panel and inhibition of MAR hydrolysis. A) Inhibition profile of **27** against viral and human ADP-ribosyl readers and erasers. Inhibition % was calculated from rFRET signals of CFP-fused binders with YFP-GAP(MAR) or YFP-GAP(PAR). B) Dot blot assay for MAR hydrolysis and inhibition of Mac1 activity. Measurements were taken in quadruplicates (n = 4). Individual data points are shown (#p<0.05 vs. MARylated SRPK2 control; *p<0.05 vs. SARS CoV-2 Mac1 control).

### Inhibition of virus replication

We recently found that a complete deletion of Mac1 from SARS-CoV-2 resulted in only modest replication defects, making it difficult to screen this virus for antiviral activity of **27** ^14^. However, Mac1 is critical for the replication of murine hepatitis virus strain JHM (MHV-JHM), a model β-CoV ^14,30^. While there may be differences in the potency of **27** against the MHV Mac1 protein compared to SARS-CoV-2, we hypothesized that at high concentrations it would likely inhibit its function, as it inhibited ADP-ribose binding by other β-CoV Mac1 proteins (Fig. 5A). We utilized two different MHV susceptible cell lines, DBT (delayed brain tumor – astrocytoma cells) and L929s (transformed fibroblasts). **27** did not have any cellular cytotoxicity in these cells as measured by an MTT assay (Fig. S3). These cells were infected with MHV-JHM at an MOI of 0.1 PFU/cell, added compound or DMSO after 1 hr of infection, and then collected both cells and supernatants at 20 –24 hpi. We found that **27** inhibited MHV-JHM replication by ∼1-2 logs at 100 µM and nearly 3 logs at 200 µM in both cell types, though it was slightly more effective in L929s (Fig. 6). These results indicate that **27** is capable of inhibiting MHV-JHM replication.

**Fig. 6.**
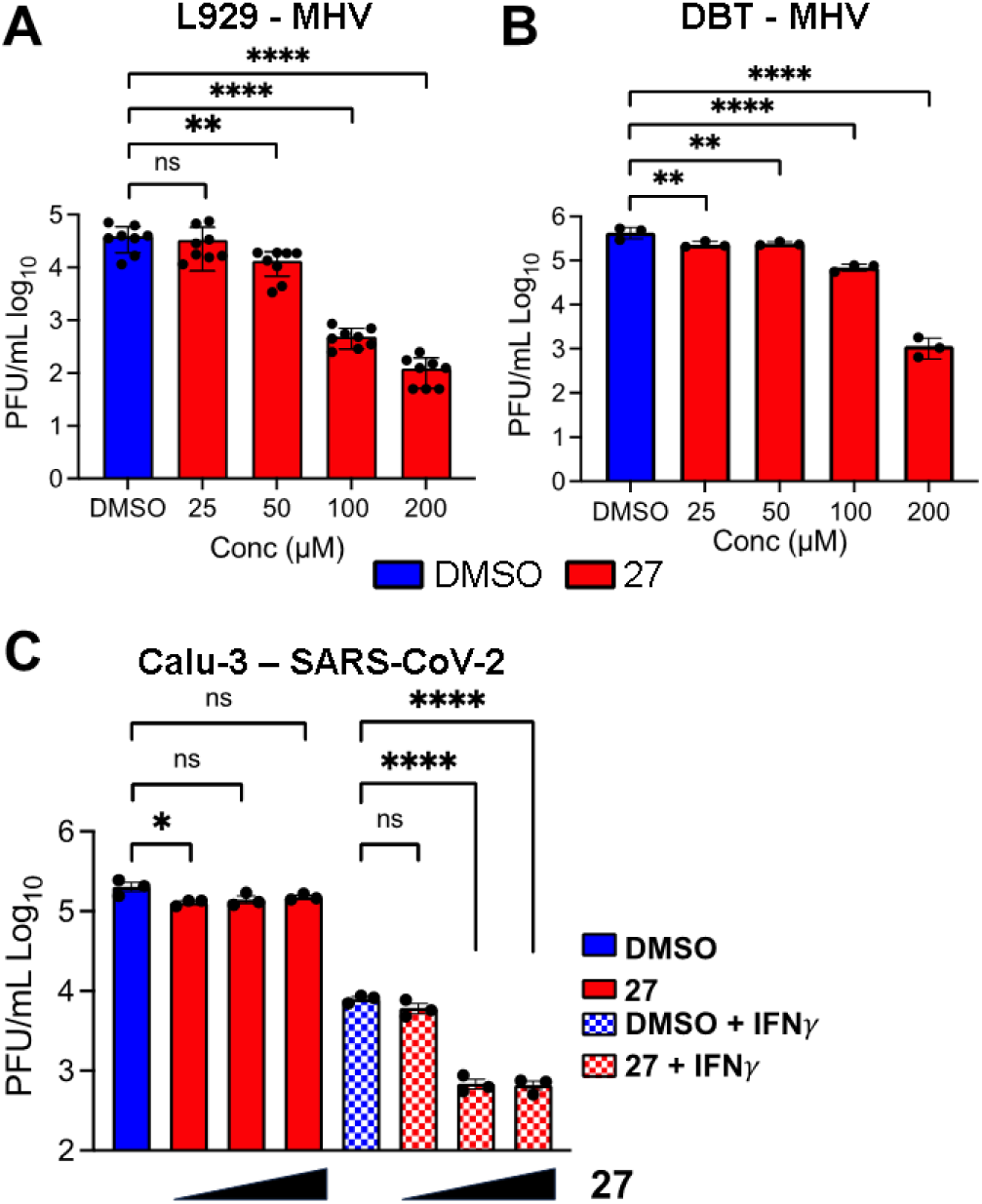
27 inhibits CoV replication. (A-B) L929 (A) or DBT (B) cells were infected with MHV-JHM at an MOI of 0.1 PFU/cell. Compound **27** was added to infected cells at 1 hpi at the indicated concentrations. Cells and supernatants were collected at 24 (A) and 20 (B) hpi and viral titers were then determined by plaque assay. (C) Calu-3 cells were either mock or IFNγ treated (100 U/ml) for 18 hrs before being infected with SARS-CoV-2 at an MOI of 0.1 PFU/cells. DMSO or Compound **27** was then added to infected cells at 12.5, 25, and 50 μM. Cells and supernatants were collected at 48 hpi and viral titers were determined by plaque assay. The results in A are the combined results of 3 independent experiments. The results in B-C are representative of 3 independent experiments. n=8 (A) and n=3 (B-C).

While SARS-CoV-2 ΔMac1 replicated normally in cell culture, it was highly sensitive to IFNγ, as pre-treatment of Calu-3 cells with IFNγ resulted in an ∼20-fold reduction in ΔMac1 replication as compared to WT virus in Calu-3 cells ^14^. Thus, we tested the ability of **27** to inhibit SARS-CoV-2 in the presence and absence of IFNγ. While **27** had minimal, if any, antiviral effects on SARS-CoV-2 in the absence of IFNγ, there was an ∼10-15-fold reduction in virus replication in the presence of IFNγ when cells were treated with as little as 25 μM 27 (Fig. 6C). Furthermore, 27 did not lead to any cytotoxicity in these cells as measured by an MTT assay (Fig. S3C). These results provide strong evidence that **27** specifically targets Mac1 during SARS-CoV-2 infection.

### Resistance mutations

To further confirm the specificity of **27** for Mac1 and to define any potential mechanisms of resistance, we performed 3 independent passages of MHV in DBTs in the presence of 100 μM **27** to identify drug-resistant mutants. After 3 passages, each sample independently passaged sample replicated at near WT levels in the presence of **27** (Fig. 7A). We further passaged each virus 3 more times, then we plaque-picked two independent isolates from separately passaged viruses. Then we isolated RNA sequenced the macrodomain by Sanger sequencing. We identified the same two mutations near the GIF motif in loop 2 of Mac1 from both isolates. These mutations of a G4526A and G4530T resulted in amino acid substitutions of the highly conserved glycine to valine and the preceding alanine to a threonine (A125T/G126V, typically used MHV Mac1 numbering) (Fig. 7B). We overlaid **27** into a model of the WT and A125T/G126V protein, demonstrating that these mutations could limit the ability of **27** to access the ADP-ribose binding site of Mac1 (Fig. 7C). This result demonstrates that **27** targets Mac1 during MHV infection.

**Fig. 7.**
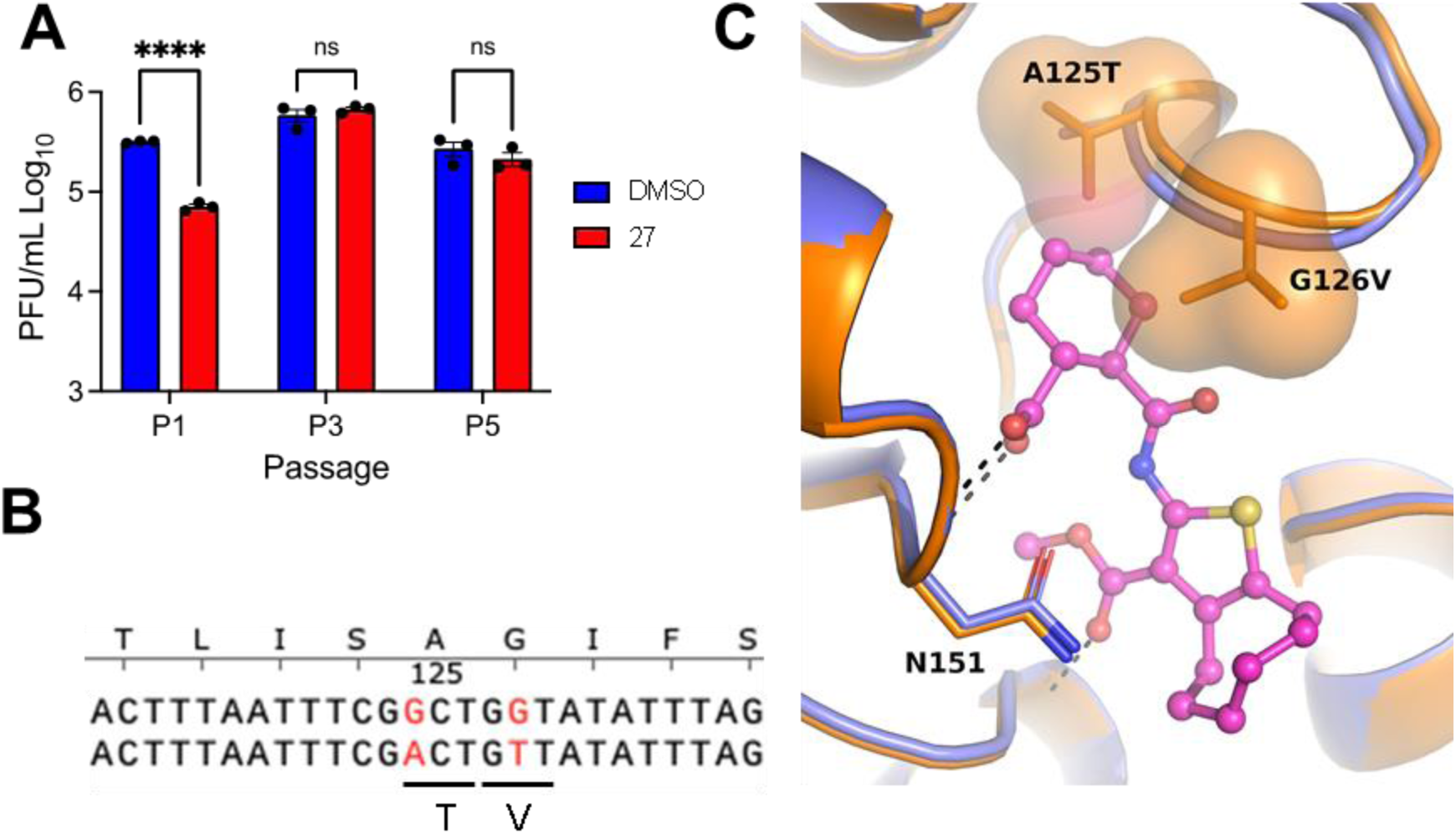
Passaging of MHV in the presence of **27** leads to drug-resistant mutants with mutations in Mac1. (A) MHV was passaged in the presence of **27** as described in Methods. Three independently passaged samples were collected and the viral titers at P1, P3, and P5 were determined by plaque assay. n=3. (B) Genetic changes identified in Mac1 following passaging in the presence of **27**. Two independent passaged plaque purified clones both evolved the same mutations; A125T and G126V. (C) Superimposition of **27** bound to SARS-CoV-2 Mac1 in a co-crystal structure onto MHV Mac1. Models of the WT (blue) and A125T/G126V mutant (orange) were generated with AlphaFold2^31^. Asn151 (Phe in Mac1) and the sidechains of the mutated residues are shown in sticks and as a molecular surface.

## Discussion and Conclusions

Viral macrodomains such as the SARS-CoV-2 Mac1 have emerged as a potential new target for antiviral agents as Mac1 is critical for replication and pathogenesis in several mouse models of CoV infection ^11^. The extreme attenuation of these viruses when Mac1 is deleted, is likely due to a combination of reduced replication and increased IFN responses. Prior work has demonstrated that the CoV Mac1 blocks IFN-I and IFN-III production and separately can promote viral replication ^12,14,32^. The attenuation of the Mac1 mutant viruses in mice make them attractive targets for the development of specific inhibitors. Furthermore, Mac1 inhibitors could function as important research tools to better understand its molecular functions, as the targets of Mac1 are still unknown. Drug repurposing efforts sparked by the COVID-19 pandemic have yielded Mac1 inhibitors with modest potency ^7,22^ and promiscuous profiles ^21^. Newly designed inhibitors have on the other hand been very polar and will require significant modification to improve their cell permeability ^20^. We reported here a new chemical scaffold inhibiting SARS-CoV-2 Mac1 and showed that it has selectivity to this protein over all human macrodomains as well as a collection of viral macrodomains. Furthermore, we showed that despite an ionizable carboxylic acid group, the best compound has a reasonable cLogD of 2.55. Our data demonstrates that this compound can enter cells as we found it could inhibit CoV replication in a dose-dependent manner.

The initial hit compounds we discovered through high-throughput screening and the discovery of multiple hit compounds with the same scaffold helped to prioritize the efforts to improve 2-amide-3-methylester thiophenes through SAR studies. These inhibitors have a unique chemotype and we were able to show using X-ray crystallography that they bind to the ADP-ribose binding cleft of Mac1 and compete with the ADP-ribose for binding (Fig. 3A, 4A). Consistent with this observation, the inhibitors also prevent hydrolysis of ADP-ribosyl group from a protein (Fig. 5B). The crystal structures revealed molecular details of the interaction that facilitated SAR and allowed us to focus the initial efforts on key areas of the compound to improve the potency, while keeping synthetic feasibility in mind. By systematic synthetic modification of ring structures and functional groups of **1**, the initial IC_50_ was improved from 14 µM to 2.1 µM allowing testing of the inhibitor for inhibition of virus infection.

Despite the extreme attenuation of a SARS-CoV-2 Mac1 deletion in mice, it had only a minimal impact in cell culture, as opposed to other CoVs, such as MHV. As MHV can be used at BSL-2 levels and requires Mac1 for replication, we first utilized this virus as a model system to test our compounds for virus inhibition. We found that compound **27** inhibited MHV replication in a dose-dependent manner in multiple cell types without substantial impacts on cell viability. However, high concentrations of compound **27** was required for inhibition (∼50-100 µM), likely due to difference in the ADP-ribose binding pocket of the MHV Mac1 compared to SARS-CoV-2 Mac1. Next, we tested the ability of **27** to inhibit SARS-CoV-2 in the presence of IFNγ as ΔMac1 was shown to be highly sensitive to IFNγ. Again, we found that **27** could inhibit virus replication, and at lower concentrations than were seen for MHV (Fig. 6C), as was expected based on the selectivity of **27** for the SARS-CoV-2 macrodomain (Fig. 5). Finally, we were able to identify a drug-resistant mutant MHV virus with mutations of conserved alanine and glycine to threonine and valine, respectively. Interestingly, these same mutations emerged in two separate passaging lineages, indicating that Mac1 may have very limited options for developing drug resistance. This result provided further evidence that **27** directly targets Mac1 during infection, but also has implications in the biochemistry of Mac1. We hypothesize that these mutations alter the ADP-ribose binding site which results in an alternative ADP-ribose binding confirmation. We previously characterized the impact of these mutations on virus replication and pathogenesis ^30^. While these mutations did not dramatically impact the replication of MHV in cell culture, they did impact its ability to cause disease in mice, indicating a loss of fitness is associated with resistance. It will be of interest to identify additional drug-resistant mutations both in Mac1 and in other regions of the viral genome to determine if resistance to Mac1 drugs generally comes with a fitness cost. Regardless, these results indicate that Mac1 inhibitors have the potential to repress CoV replication and could be further developed into chemical probes for identifying Mac1 functions or potentially even anti-viral therapeutics.

There are 16 macrodomains in humans which in certain cases could also be promising drug targets. ^33^. To evaluate likely off-targets we evaluated inhibition of **1** and **27** against a panel of all human macrodomain. This showed that the chemotype is selective towards CoV Mac1 and this minimal cross-reactivity may be an important feature in the context of antiviral therapeutics. Surprisingly, there was also clear selectivity towards SARS-CoV-2 over the other tested viral macrodomains despite the high similarity. The selectivity for Mac1 over the other macrodomains can be assessed by comparing the available crystal structures and the selectivity can be rationalized to result from differences in the active site residues near the binding pocket (Fig. S4) ^34,35^. These features may be used to utilize the 2-amide-3-methylester thiophene-core also for inhibition of other macrodomains with suitable modifications.

While the results represent an important step in developing Mac1 inhibitors, the potency should be improved further to generate a more effective chemical probe. From the pharmacological point of view, compound properties such as potential hydrolysis of a methyl ester, removal of a negative charged carboxylate and potential metabolic liabilities need to be considered in future optimization efforts. The well-defined experimental binding mode, initial activity in virus inhibition and established synthetic methods described here will enable these studies towards validating Mac1 as a therapeutic pharmacological antiviral target.

### Eperimental Section

#### Chemistry

Commercial reagents and solvents were used as received, except for malononitrile which was recrystallized from diethyl ether prior to use. Moisture-sensitive reactions were carried out under argon in oven-dried glasswares. Microwave-assisted reactions were performed using Biotage Initiator microwave reactor. ^1^H (400 MHz) and ^13^C (100 MHz) characterizations were performed with Bruker Avance 400 MHz NMR spectrometer, and samples were prepared using CDCl_3_ or *d*_6_-DMSO as a solvent. NMR and HPLC data were processed with Spectrus Processor 2019.1.1 program (ACDLabs). ESI+ TOF MS characterizations were performed using Thermo Scientific QExactive Plus Hybrid Quadrupole-Orbitrap high resolution mass spectrometer with ACQUITY UPLC® BEH C18 column, and the compounds were detected in positive and negative mode. Purities of the compounds were assessed with Shimadzu HPLC instrument using water+0.1% TFA (A) and acetonitrile+0.1% TFA (B) as a mobile phase (gradient B 5% to 90% over 10 minutes) with Waters Atlantis T3 column and detection at 220 nm and 260 nm wavelengths with Shimadzu SDP-10AVP UV-vis detector. Purities of compounds were determined to be >95%, except for commercial compound **13** which had a purity of ≥88%. All reactions were monitored with thin-layer chromatography using silica gel 60-coated aluminum sheets and spots were visualized with UV-lamp or iodine staining.

Compound structures were drawn using ChemSketch (ACDLabs) and cLogD values were calculated with MarwinSketch 22.6 (Chemaxon).

#### Synthesis

##### General method A

To a microwave reaction tube containing a solution of cycloheptanone and elemental sulfur in DMF, an activated nitrile compound was added along with a dropwise addition of pyrrolidine. The reaction tube was sealed and subjected to microwave irradiation (absorption level very high) and 60 °C temperature for 30 minutes. Next, the reaction solution was filtered through a thin pad of silica gel rinsed with ethyl acetate and evaporated under reduced pressure. The crude products were purified with flash chromatography (SiO_2_) with hexane:ethyl acetate gradient elution.

##### General method B

A 2-aminothiophene starting material was added with a small excess of an appropriate anhydride to a reaction tube and ground with glass rod. A stirring magnet was added, and the mixture was kept at an argon flow for a few minutes. After that, the tube was sealed and transferred to an oil bath. The reaction mixture was stirred at a slow speed and heated overnight, typically ca. 16 h. Temperatures of the used oil baths are given in parentheses.

###### Methyl 2-amino-5,6,7,8-tetrahydro-4H-cyclohepta[b]thiophene-3-carboxylate (**9a**)

Title compound was prepared using General method A. Specific amounts of used chemicals: cycloheptanone 120 µL (1.0 mmol), methyl cyanoacetate (activated nitrile) 100 µL (1.1 mmol), elemental sulfur 35.4 mg (1.1 mmol), pyrrolidine 80 µL (1.0 mmol), and DMF 3.0 mL. Flash chromatography (SiO_2_) with hexane:ethyl acetate 10:1 to 3:1 gradient elution. Yield: 160.9 mg, 71%, yellowish solid. ^1^H NMR (400 MHz, CDCl_3_) δ ppm 1.57–1.72 (m, 4 H), 1.82 (m, 2H), 2.53–2.68 (m, 2H), 2.92–3.03 (m, 2H), 3.81 (s, 3H), 5.78 (br s, 2H).

###### Methyl 2-amino-4,5,6,7-tetrahydro-1-benzothiophene-3-carboxylate (**9b**)

Cyclohexanone (0.5 mL, 4.8 mmol), methyl cyanoacetate (0.465 mL, 5.27 mmol) and elemental sulfur (185.8 mg, 5.77 mmol) were dissolved in methanol (5 mL), followed by a dropwise addition of diethylamine (0.25 mL). The mixture was allowed to react with constant stirring at room temperature. After 25 h, the reaction mixture was cooled with an ice-water bath, and the formed precipitate was filtered. The precipitate was washed with ice-cold methanol and dried in oven (55°C), yielding the title compound **9b** as a white solid (566.7 mg, 56%). ^1^H NMR (400 MHz, CDCl_3_) δ ppm 1.67–1.85 (m, 4 H) 2.44–2.55 (m, 2 H) 2.66–2.73 (m, 2 H) 3.79 (s, 3 H) 5.93 (br s, 2 H).

###### 2-amino-5,6,7,8-tetrahydro-4H-cyclohepta[b]thiophene-3-carbonitrile (9c)

Title compound was prepared using General method A. Specific amounts of used chemicals: cycloheptanone 370 µL (3.1 mmol), malononitrile (activated nitrile) 209.5 mg (3.1 mmol), elemental sulfur 101.5 mg (3.1 mmol), pyrrolidine 250 µL (3.1 mmol), and DMF 3.5 mL. Yield: 428.8 mg, 71%, yellow solid. ^1^H NMR (400 MHz, CDCl_3_) *δ* ppm 1.60–1.70 (m, 4H), 1.78–1.87 (m, 2H), 2.54–2.68 (m, 4H), 4.48 (br s, 2H).

###### 1-(2-amino-5,6,7,8-tetrahydro-4H-cyclohepta[b]thiophen-3-yl)ethan-1-one (9d)

In a 100 mL round bottom flask, 928.2 mg (11 mmol) of cyanoacetone was dissolved in 16 mL of methanol. Next, 1190 µL (10 mmol) of cycloheptanone and 360.5 mg (11 mmol) of elemental sulfur were added into the flask. 1.1 mL (11 mmol) of piperidine was added in a dropwise fashion with simultaneous stirring. The flask was equipped with a reflux condenser and heated to 55 °C for 21 hours. After that, the reaction mixture was vacuum filtered through a thin pad of celite into a flask containing ice-cold water. The formed precipitate was filtered into a sintered funnel, washed with copious amounts of deionized water, and dried in an oven (55 °C), yielding the title compound **9d** as a tan solid (261.8 mg, 21%). ^1^H NMR (400 MHz, CDCl_3_) δ ppm 1.65–1.72 (m, 4 H), 1.80–1.89 (m, 2 H), 2.42 (s, 3 H), 2.57–2.62 (m, 2 H), 2.80–2.8 (m, 2 H), 6.59 (br s, 2 H).

###### Methyl 2-amino-4,5,6,7,8,9-hexahydrocycloocta[b]thiophene-3-carboxylate (**9e**)

Title compound was prepared using General method A. Specific amounts of used chemicals: cyclooctanone 415.4 mg (3.3 mmol), methyl cyanoacetate (activated nitrile) 320 µL (3.63 mmol), elemental sulfur 116.4 mg (3.63 mmol), pyrrolidine 270 µL (3.3 mmol), and DMF 10 mL. Yield: 393.8 mg, 50%, orange amorphous solid. ^1^H NMR (400 MHz, DMSO-*d*_6_) δ ppm 1.20–1.26 (m, 2 H), 1.36–1.43 (m, 2 H), 1.44–1.57 (m, 4 H), 2.52–2.57 (m, 2 H), 2.70–2.78 (m, 2 H), 3.68 (s, 3 H), 7.16 (s, 2 H).

###### 6-{[3-(methoxycarbonyl)-5,6,7,8-tetrahydro-4H-cyclohepta[b]thiophen-2-yl]carbamoyl}cyclohex-3-ene-1-carboxylic acid (1r)

Title compound was prepared using General method B (120 °C). Specific amounts of used chemicals: 99.4 mg (0.44 mmol) of **9a** and 66.2 mg (0.45 mmol) *cis*-1,2,3,6-tetrahydrophthalic anhydride. The title compound was purified with flash chromatography using DCM:MeOH 98:2 to 92:8 gradient elution. Yield: 131.5 mg, 79%, tan powder. ^1^H NMR (400 MHz, DMSO-*d*_6_) δ ppm 1.48–1.61 (m, 4 H), 1.74–1.84 (m, 2 H), 2.28–2.45 (m, 4 H), 2.64–2.71 (m, 2 H), 2.92–3.00 (m, 2 H), 3.01–3.08 (m, 1 H), 3.09–3.14 (m, 1 H), 3.84 (s, 3 H), 5.69 (s, 2 H), 11.05 (br s, 1 H), 12.31 (br s, 1 H). ^13^C NMR (101 MHz, CDCl_3_) δ ppm 26.0, 26.3, 27.0, 27.8, 28.2, 28.6, 32.2, 39.6, 40.8, 51.6, 112.8, 124.3, 126.0, 131.1, 136.3, 145.5, 167.2, 170.4, 178.1. HRMS (ESI+, TOF) m/z: [M+H]^+^ calcd for C_19_H_23_NO_5_S 378.1369; found 378.1369.

###### 2-{[3-(methoxycarbonyl)-4,5,6,7-tetrahydro-1-benzothiophen-2-yl]carbamoyl}benzoic acid (3r)

Title compound was prepared using General method B (130 °C). Specific amounts of used chemicals: 32.2 mg (0.15 mmol) of **9b** and 30.7 mg (0.2 mmol) of phthalic anhydride. The title compound was purified with recrystallization from isopropanol. Yield: 28.0 mg, 51%, off-white solid. 1H NMR (400 MHz, DMSO-*d*_6_) δ ppm 1.67–1.81 (m, 4 H), 2.61–2.68 (m, 2 H), 2.68–2.75 (m, 2 H), 3.78 (s, 3 H), 7.64–7.74 (m, 3 H), 7.89 (m, 1 H), 11.32 (s, 1 H), 13.26 (br s, 1 H). ^13^C NMR (101 MHz, CDCl_3_) δ ppm 22.3, 23.0, 24.4, 26.3 51.6, 112.5, 127.75, 127.79, 129.5, 131.0 131.1, 131.9, 132.8, 135.7, 147.2, 165.7, 167.2, 168.9. HRMS (ESI+, TOF) m/z: [M+H]^+^ calcd for C_18_H_17_NO_5_S 360.0900; found: 360.0897.

###### 2-{[3-(methoxycarbonyl)-5,6,7,8-tetrahydro-4H-cyclohepta[b]thiophen-2-yl]carbamoyl}benzoic acid (10)

Title compound was prepared using General method B (120 °C). Specific amounts of used chemicals: 93.2 mg (0.41 mmol) of **9a** and 62.1 mg (0.42 mmol) of phthalic anhydride. The title compound was purified from the crude reaction mixture by precipitating with cold DCM., The formed precipitate was filtered, washed with cold DCM, and dried in an oven (55 °C). Yield: 118.2 mg, 76%, tan solid. ^1^H NMR (400 MHz, DMSO-*d*_6_) δ ppm 1.51–1.65 (m, 4 H), 1.77–1.87 (, 2 H), 2.69–2.78 (m, 2 H), 2.91–3.01 (m, 2 H), 3.78 (s, 3 H), 7.60–7.74 (m, 3 H), 7.85–7.90 (m, 1 H), 11.16 (s, 1 H), 13.22 (br s, 1 H). 13C NMR (101 MHz, DMSO-d6) δ ppm 26.8, 27.4, 27.5, 28.0, 31.7, 51.8, 114.1, 127.6, 129.7, 130.6, 130.77, 130.84, 131.9, 135.9, 136.2, 143.0, 165.2, 165.5, 167.4. HRMS (ESI+, TOF) m/z: [M+H]^+^ calcd for C_19_H_19_NO_5_S 374.1056; found 374.1055.

###### 6-{[3-(methoxycarbonyl)-4,5,6,7-tetrahydro-1-benzothiophen-2-yl]carbamoyl}cyclohex-3-ene-1-carboxylic acid (11)

Title compound was prepared using General method B (120 °C). Specific amounts of used chemicals: 31.8mg (0.15 mmol) of **9b** and 23.2 mg (0.15 mmol) *cis*-1,2,3,6-tetrahydrophthalic anhydride. The title compound was purified with flash chromatography using hexane:ethyl acetate 1:1 to EtOAc 100% gradient elution. Yield: 44.6 mg, 81%, tan solid. ^1^H NMR (400 MHz, CDCl_3_) δ ppm 1.72–1.85 (m, 4 H), 2.36–2.47 (m, 1 H), 2.50–2.59 (m, 1 H), 2.60–2.72 (m, 4 H), 2.73–2.79 (m, 2 H), 3.11–3.18 (m, 1 H), 3.22–3.20 (m, 1 H), 3.87 (s, 3 H), 5.76 (s, 2 H), 11.62 (br s, 1 H). ^13^C NMR (101 MHz, CDCl3) δ ppm 22.8, 23.0, 24.3, 26.1, 26.3, 26.5, 39.6, 41.1, 51.5, 111.7, 124.3, 126.0, 127.1, 130.8, 147.4, 167.2, 170.7, 176.6. HRMS (ESI+, TOF) m/z: [M+H]^+^ calcd for C_18_H_21_NO_5_S 364.1213; found 364.1210.

###### (2-{[3-(methoxycarbonyl)-4H,5H,6H,7H,8H-cyclohepta[b]thiophen-2-yl]carbamoyl}cyclohexane-1-carboxylic acid) (**15r**)

Title compound was prepared using General method B (90 °C). Specific amounts of used chemicals: 101.8 mg of **9a** and 89.1 mg of *cis*-1,2-cyclohexanedicarboxylic anhydride. The title compound was purified with recrystallization from chloroform:hexane solution. After that, the filtered solid was dissolved in chloroform, washed with deionized water, organic phase dried with Na_2_SO_4_, filtered and evaporated to dryness. The pure product was dried in a vacuum oven (60 °C). Yield: 138.7 mg, 81%, tan powder. ^1^H NMR (400 MHz, DMSO-*d*_6_) δ ppm 1.31–1.47 (m, 3 H), 1.48–1.63 (m, 5 H), 1.64–2.06 (m, 6 H), 2.62–2.72 (m, 2 H), 2.82–3.03 (m, 4 H), 3.83 (s, 3 H), 11.01 (s, 1 H), 12.17 (br s, 1 H). ^13^C NMR (101 MHz, DMSO-*d*_6_) δ ppm 22.9, 23.3, 25.6, 26.4, 26.7, 27.41, 27.46, 27.9, 31.7, 41.7, 42.7, 51.8, 112.4, 129.9, 135.9, 144.2, 165.7, 171.0, 174.6. HRMS (ESI+, TOF) m/z: [M+H]^+^ calcd for C_19_H_25_NO_5_S; 380.1526 found 380.1522.

###### 6-[(3-cyano-5,6,7,8-tetrahydro-4H-cyclohepta[b]thiophen-2-yl)carbamoyl]cyclohex-3-ene-1-carboxylic acid (21)

Title compound was prepared using General method B (90 °C). Specific amounts of used chemicals: 85.2 mg (0.44 mmol) of **9c** and 81.1 mg (0.53 mmol) of *cis*-1,2,3,6-tetrahydrophthalic anhydride. The title compound was purified from the crude reaction mixture by precipitating with cold chloroform. The formed precipitate was filtered, washed with cold chloroform, and dried in an oven (55 °C). Yield: 58,1 mg, 38%, off-white powder. ^1^H NMR (400 MHz, DMSO-*d*_6_) δ ppm 1.46–1.67 (m, 4 H), 1.71–1.87 (m, 2 H), 2.20–2.39 (m, 3 H), 2.57–2.60 (m, 1 H), 2.61–2.70 (m, 4 H), 2.87–2.96 (m, 1 H), 3.21–3.27 (m, 1 H), 5.56–5.71 (m, 2 H), 11.37 (br s, 1 H), 12.21 (br s, 1 H). ^13^C NMR (101 MHz, DMSO-*d*_6_) δ ppm 25.6, 27.0, 27.3, 28.0, 29.0, 29.1, 32.0, 40.3, 40.7, 96.0, 114.9, 124.7, 125.6, 132.0, 135.6, 144.5, 169.8, 177.8. HRMS (ESI+, TOF) m/z: [M+H]^+^ calcd for C_18_H_20_N_2_O_3_S; 345.1263 found 345.1267.

###### 6-[(3-acetyl-5,6,7,8-tetrahydro-4H-cyclohepta[b]thiophen-2-yl)carbamoyl]cyclohex-3-ene-1-carboxylic acid (22)

Title compound was prepared using General method B (90 °C). Specific amounts of used chemicals: 57.4 mg (0.27 mmol) of **9d** and 52.0 mg (0.34 mmol) of *cis*-1,2,3,6-tetrahydrophthalic anhydride. The title compound was purified with flash chromatography using DCM:MeOH 95:5 elution. Yield: 57.7 mg, 58%, tan powder. ^1^H NMR (400 MHz, DMSO-*d*_6_) δ ppm 1.52–1.67 (m, 4 H), 1.75–1.87 (m, 2 H), 2.29–2.37 (m, 1 H), 2.41–2.47 (m, 3 H), 2.48 (s, 3 H), 2.65–2.76 (m, 2 H), 2.79–2.90 (m, 2 H), 3.00–3.12 (m, 2 H), 5.68 (m, 2 H), 11.7 (br s, 1 H), 12.28 (br s, 1 H). ^13^C NMR (101 MHz, CDCl_3_) δ ppm 26.2, 26.4, 26.7, 27.6, 28.4, 29.3, 31.5, 31.9, 39.7, 41.1, 123.4, 124.3, 126.0, 131.8, 135.2, 145.6, 171.4, 176.5, 197.7. HRMS (ESI+, TOF) m/z: [M+H]^+^ calcd for C_19_H_23_NO_4_S; 362.1420 found 362.1416.

###### Methyl 2-[6-(methoxycarbonyl)cyclohex-3-ene-1-amido]-4H,5H,6H,7H,8H-cyclohepta[b]thiophene-3-carboxylate (**23**)

221.3 mg (0.586 mmol) of compound **1r** was dissolved in 23 mL of methanol, and 200 µL of conc H_2_SO_4_ was added. The reaction was kept at reflux for 16 hours and after the completion of the reaction according to TLC, the reaction mixture was concentrated with reduced pressure. Cold water was added to the concentrated mixture, and the formed precipitate was filtered. The precipitate was subjected to flash chromatography with DCM:hexane 1:1 to DCM 100% to DCM:MeOH 95:5 gradient elution. Yield: 131.8 mg, 57%, white amorphous solid. ^1^H NMR (400 MHz, DMSO-*d*_6_) δ ppm 1.46–1.62 (m, 4 H), 1.75–1.84 (m, 2 H), 2.35–2.42 (m, 2 H), 2.44–2.48 (m, 2 H), 2.65–2.71 (m, 2 H), 2.90–3.03 (m, 2 H), 3.07–3.15 (m, 1 H), 3.17–3.26 (m, 1 H), 3.57 (s, 3 H), 3.84 (s, 3 H), 5.69 (m, 2 H), 10.99 (s, 1 H). ^13^C NMR (101 MHz, CDCl_3_) δ ppm 25.7, 26.2, 27.0, 27.8, 28.1, 28.6, 32.2, 39.8, 40.9, 51.5, 51.9, 112.6, 124.3, 125.8, 130.8, 136.2, 145.7, 167.1, 170.2, 173.5. HRMS (ESI+, TOF) m/z: [M+H]^+^ calcd for C_20_H_25_NO_5_S 392.1526; found 392.1524.

###### 2-[(2-carboxycyclohexane-1-carbonyl)amino]-5,6,7,8-tetrahydro-4H-cyclohepta[b]thiophene-3-carboxylic acid (24)

In a round-bottom flask, 31.5 mg (0.78 mmol) of sodium hydroxide was dissolved in deionized water and 56.9 mg (0.15 mmol) of compound **15r** was added. The solution was kept at reflux for 2 hours. When no more starting material was present according to TLC, the reaction mixture was allowed to cool to room temperature and then transferred to an ice-water bath. Into the mixture, 10 mL of ethyl acetate was added, after which the aqueous phase was acidified with 2 M hydrochloric acid to pH 1–2. The aqueous layer was further extracted two times with 10 mL ethyl acetate. The combined organic phases were washed with brine, dried with MgSO_4_, filtered and evaporated to dryness. Yield: 54.2 mg, 99%, yellowish powder. ^1^H NMR (400 MHz, DMSO-*d*_6_) δ ppm 1.32–1.46 (m, 3 H), 1.47–1.62 (m, 5 H), 1.68 (m, 1 H), 1.75–1.95 (m, 4 H), 1.96–2.08 (m, 1 H), 2.67 (m, 2 H), 2.77–2.92 (m, 2 H), 2.92–3.11 (m, 2 H), 11.36 (br s, 1 H), 12.65 (br s, 2 H). ^13^C NMR (101 MHz, DMSO-*d*_6_) δ ppm 22.9, 23.4, 25.4, 26.4, 26.8, 27.3, 27.6, 27.9, 31.9, 41.7, 42.8, 112.8, 129.5, 136.4, 144.4, 167.2, 170.8, 174.6. HRMS (ESI+, TOF) m/z: [M+H]^+^ calcd for C_18_H_23_NO_5_S 366.1369; found 366.1369.

###### (1S,2S)-2-{[3-(methoxycarbonyl)-4H,5H,6H,7H,8H-cyclohepta[b]thiophen-2-yl]carbamoyl}cyclohexane-1-carboxylic acid)) (**25**)

Title compound was prepared using General method B (80 °C). Specific amounts of used chemicals: 100.9 mg (0.45 mmol) of **9a** and 76.6 mg (0.5 mmol) of (-)*-trans*-1,2-cyclohexanedicarboxylic anhydride. The title compound was purified with flash chromatography (SiO_2_) and eluted with DCM:MeOH 98:2 to 93:7 gradient. Yield: 134.3 mg, 79%, tan powder. ^1^H NMR (400 MHz, DMSO-*d*_6_) δ ppm 1.22–1.39 (m, 4 H), 1.45–1.61 (m, 4 H), 1.67–1.84 (m, 4 H), 1.85–1.93 (m, 1 H), 1.96–2.04 (m, 1 H), 2.43–2.48 (m, 1 H), 2.62–2.75 (m, 3 H), 2.88–2.98 (m, 2 H), 3.83 (s, 3 H), 10.79 (s, 1 H), 12.18 (br s, 1 H). ^13^C NMR (101 MHz, CDCl_3_) δ ppm 25.07, 25.11, 27.1, 27.8, 28.2, 28.6, 29.1, 29.4, 32.3, 44.3, 46.3, 51.5, 112.6, 130.9, 136.2, 145.7, 167.2, 171.7, 179.6. HRMS (ESI+, TOF) m/z: [M+H]^+^ calcd for C_19_H_25_NO_5_S; 380.1526 found 380.1523.

###### (1R,2R)-2-{[3-(methoxycarbonyl)-4H,5H,6H,7H,8H-cyclohepta[b]thiophen-2-yl]carbamoyl}cyclohexane-1-carboxylic acid (26)

Title compound was prepared using General method B (80 °C). Specific amounts of used chemicals: 99.4 mg (0.44 mmol) of **9a** and 79.7 mg (0.52 mmol) of (+)*-trans*-1,2-cyclohexanedicarboxylic anhydride. The crude reaction product was dissolved in chloroform and washed with water. The chloroform layer was dried with Na_2_SO_4_, filtered and evaporated to dryness. After that, the evaporated crude product was subjected to flash chromatography (SiO_2_) and eluted with DCM:MeOH 98:2 to 95:5 gradient. Yield: 129.9 mg, 78%, tan powder. ^1^H NMR (400 MHz, DMSO-*d*_6_) δ ppm 1.21–1.42 (m, 4 H), 1.50–1.62 (m, 4 H), 1.66–1.85 (m, 4 H), 1.86–1.94 (m, 1 H), 1.97–2.03 (m, 1 H), 2.43–2.49 (m, 1 H), 2.62–2.73 (m, 3 H), 2.87–3.01 (m, 2 H), 3.84 (s, 3 H), 10.80 (s, 1 H), 12.20 (br s, 1 H). ^13^C NMR (101 MHz, DMSO-*d*_6_) δ ppm 25.07, 25.11, 27.1, 27.8, 28.2, 28.6, 29.1, 29.4, 32.3, 44.3, 46.3, 51.5, 112.6, 130.9, 136.2, 145.7, 167.1, 171.7, 179.8. HRMS (ESI+, TOF) m/z: [M+H]^+^ calcd for C_19_H_25_NO_5_S; 380.1526 found 380.1522.

###### ((1R,2R)-2-{[3-(methoxycarbonyl)-4H,5H,6H,7H,8H,9H-cycloocta[b]thiophen-2-yl]carbamoyl}cyclohexane-1-carboxylic acid) (**27**)

Title compound was prepared using General method B (85 °C). Specific amounts of used chemicals: 108.3 mg (0.45 mmol) **9e** and 80.2 mg (0.5 mmol) of (+)*-trans*-1,2-cyclohexanedicarboxylic anhydride. The title compound was purified with flash chromatography using DCM 100% to DCM:MeOH 92:8 gradient elution. Yield: 111.5 mg, 63%, tan powder. ^1^H NMR (400 MHz, DMSO-*d*_6_) δ ppm 1.14–1.23 (m, 2 H), 1.25–1.47 (m, 6 H), 1.49–1.63 (m, 4 H), 1.68–1.81 (m, 2 H), 1.86–1.95 (m, 1 H), 1.97–2.05 (m, 1 H), 2.46 (m, 1 H), 2.62–2.75 (m, 3 H), 2.84 (m, 2 H), 3.84 (s, 3 H), 10.99 (s, 1 H), 12.21 (br s, 1 H). ^13^C NMR (101 MHz, DMSO-*d*_6_) δ ppm 25.06, 25.11, 25.3, 25.5, 26.5, 26.8, 29.1, 29.5, 29.9, 32.2, 44.3, 46.2, 51.4, 111.6, 129.8, 133.1, 147.3, 167.0, 171.7, 180.3. HRMS (ESI+, TOF) m/z: [M+H]^+^ calcd for C_20_H_27_NO_5_S 394.1682; found 394.1681.

###### Methyl 2-[(1R,2R)-2-(methoxycarbonyl)cyclohexaneamido]-4H,5H,6H,7H,8H,9H-cycloocta[b]thiophene-3-carboxylate (**28**)

To a flask containing 173.5 mg (0.44 mmol) of compound **27** was added 20 mL of methanol along with 10 drops of conc H_2_SO_4_. The reaction mixture was kept at reflux for two hours, after which the solution was concentrated to a smaller volume with a rotary evaporator and co-evaporated with three portions of toluene. The crude product dissolved in ca. 5 mL of toluene was washed two times with 10 mL of deionized water, two times with 10 mL 0.1 M NaHCO_3_ and with 10 mL of brine. The organic phase was dried with MgSO_4_, filtered and evaporated to dryness. The crude product was subjected to flash chromatography (SiO_2_) with DCM:hexane 2:1 to DCM 100% gradient elution yielding the title compound **28** as a colorless amorphous solid (153.5 mg, 85%). ^1^H NMR (400 MHz, CDCl_3_) δ ppm 1.22–1.31 (m, 2 H), 1.31–1.41 (m, 3 H), 1.41–1.52 (m, 3 H), 1.58–1.68 (m, 4 H), 1.79–1.90 (m, 2 H), 2.01–2.09 (m, 1 H), 2.11–2.18 (m, 1 H), 2.64–2.83 (m, 4 H), 2.85–2.91 (m, 2 H), 3.65 (s, 3 H), 3.90 (s, 3 H), 11.38 (s, 1 H). ^13^C NMR (101 MHz, CDCl_3_) δ ppm 25.10, 25.12, 25.3, 25.5, 26.5, 26.8, 29.1, 29.5, 29.9, 32.3, 44.7, 46.8, 51.4, 51.9, 111.5, 129.7, 133.1, 147.3, 167.0, 171.9, 175.3. HRMS (ESI+, TOF) m/z: [M+H]^+^ calcd for C_21_H_29_NO_5_S 408.1839; found 408.1839.

### Expression constructs, Protein Expression and Purification

Recombinant proteins were produced as previously described ^23^. All the expression constructs are available at Addgene. For the expression of CFP (e.g. Addgene #173083) and YFP tagged recombinant proteins (Addgene #173080), genes were inserted between N-terminal 6x His-tag followed by CFP/YFP tag and a TEV protease cleavage site of pNIC28-Bsa4. Proteins were expressed in *E. coli* strains Rosetta2(DE3) or BL21(DE3). SARS-CoV-2 Mac1 without CFP-tag was cloned and expressed from pNH-TrxT (Addgene #173084).

YFP-GAP (where GAP-tag is a C-terminal peptide K345-F354 of Gαi) and CFP-SARS-CoV-2 Mac1 (residues Q1084-E1192) as well as other CFP-tagged MAR/PAR binders used in profiling were purified using nickel affinity and size exclusion chromatography as described in ^21^. CFP-TNKS2-ARC4 and YFP-peptide for counter screen was produced analogously ^24^. N-terminal His- and TrxT-tags were removed with TEV-protease from SARS-CoV-2 Mac1 to be used for thermal shift assay and for crystallization.

PARP10 and SRPK2 used in the hydrolysis assay were produced as previously described ^36^.

### FRET assay

The ADP-ribosylation binding assay based on FRET technology was used to screen large compound libraries, dose-response measurements and profiling of the hit compounds ^21^. The assay was performed in 384-well black polypropylene flat-bottom plates (Greiner, Bio-one). Reactions were prepared in the assay buffer (10 mM Bis-Tris propane pH 7.0, 3% (w/v) PEG 20,000, 0.01% (v/v) Triton X-100 and 0.5 mM TCEP) with a reaction volume of 10 μL per well. The reactions consisted of 1 μM CFP-SARS-CoV-2 Mac1 and 5 μM YFP-GAP MARylated with a fragment of Pertussis toxin ^21,37^. Dispensing of the solutions for setting up the reaction was carried out by using Microfluidic Liquid Handler (MANTIS®, Formulatrix, Bedford, MA, USA). For screening, profiling and IC_50_ experiments, compounds were dispensed by the Echo 650 acoustic liquid dispenser (Labcyte, Sunnyvate, CA). All the measurements were taken with Tecan Infinite M1000 pro plate reader. The FRET emission signal was measured at 477 nm (10 nm bandwidth) and 527 nm (10 nm bandwidth) upon excitation at 410 nm (20 nm bandwidth). The ratiometric FRET value (rFRET) was calculated by dividing the raw fluorescence intensities at 527 nm by 477 nm after blank deduction.

### Assay validation for SARS-CoV-2 Mac1

The assay quality and performance for the FRET-based assay was also tested for the above-described conditions. Validation was performed on 384-well plates containing 176 maximal and minimal signal points and 32 blank wells. Repeatability of the maximal and minimal signal was measured between different wells, plates and days. In total, five control plates were prepared. The blank wells contained assay buffer only. Maximal signal wells contained assay buffer, 1 μM CFP-SARS-CoV-2 Mac1 and 5 μM MARylated YFP-GAP. Minimal signal wells contained assay buffer, 1 μM CFP-SARS-CoV-2 Mac1 and 5 μM MARylated YFP-GAP and 200 μM ADPr. The data points were used to calculate the mean of maximal and minimal signals, coefficients of variations and standard deviations (Table S1). Three plates were measured on the first day, one plate on the second day, and one final plate on the third day. The overall quality of the assay was determined with: signal-to-noise ratio (S/N), signal-to-background ratio (S/B) and screening window coefficient (Zʹ) (Table S1) ^38,39^.

### Screening of inhibitors

A compound library of 30,000 compounds from SPECS consortium collection was screened against SARS-CoV-2 Mac1 macrodomain. The 384-well plates containing compounds (0.03 μL) in singlets were supplied by the Institute for Molecular Medicine, Finland (FIMM) and the screening was carried out at 30 μM compound concentration. The 4 μL of the FRET mixture consisting of 1 μM CFP-SARS-CoV-2 Mac1 and 5 μM MARylated YFP-GAP was added to the plates followed by the addition of 6 µL of assay buffer to make up the final volume to 10 µL and the plate was measured. Each plate contained blank wells (assay buffer only), positive control wells (assay buffer and FRET mixture), and negative control wells (assay buffer, FRET mixture and 200 μM ADPr). The positive and negative control signals were used to calculate each compound’s inhibition. In order to filter compounds that display intrinsic fluorescence or non-specifically inhibit the FRET interaction at the measured wavelengths, additional filters were applied. If the intensity of the emission at 477 nm was ≤ 1.2 times the average of minimum control wells or if the emission at 527 nm was ≤ 1.2 times the average of maximum control wells the compounds were excluded. These efficiently filter out inherently fluorescent compounds, but do not exclude protein aggregators that result in negative inhibition visible in Fig. 1A. The plates were read additionally with an excitation wavelength of 430 nm (20 nm bandwidth) that allowed monitoring whether the inhibition % was similar with both excitation wavelengths as expected for true SARS-CoV-2 Mac1 inhibitors or whether there was a large difference resulting from compound interference with FRET signal. Based on this final criteria, 70 compounds were finally excluded from the screening. Analysis of the data was done in GraphPad Prism version 8.02 for windows (GraphPad Software, La Jolla, CA, USA).

### Counter screening

Validation of the hit compounds obtained from initial screening was performed against CFP-TNKS2 ARC4 and YFP-TBM utilizing a previously developed FRET-based assay ^24^. The hit compounds were screened in duplicates at a concentration of 30 μM. The pre-plated compounds (0.03 μL) were supplied by the Institute for Molecular Medicine, Finland (FIMM). 4 μL of the FRET mixture consisting of (100 nM CFP-TNKS2 ARC4 and 200 nM YFP-TBM were added to the plates and 10 µL ^2424^(Sowa *et al*, 2020)(Sowa *et al*, 2020) final volume was made by adding 6 µL assay buffer. The plate also contained blank wells (assay buffer only), positive control wells (assay buffer and FRET mixture), negative control wells (assay buffer, FRET mixture and 1 M GdnHCl). The plate was measured as for SARS-CoV-2 Mac1 and the positive and negative control signals were used to calculate inhibition-%.

### Potency measurements

Concentration-response curves for the hit compounds were measured using 100 μM (1 mM for ALC1) to 0.003 μM with half-log dilutions. Measurements were carried out in quadruplicates and repeated three times for potent compounds. Reaction was prepared by addition of 100 nL of the compound to the assay plates followed by the addition of 4 μL of the FRET mixture consisting of 1 μM CFP-SARS-CoV-2 Mac1 and 5 μM MARylated YFP-GAP and addition of 6 µL of assay buffer. IC_50_ curves were fitted using sigmoidal dose response curve (four variables) in GraphPad Prism version 8.02.

### Differential scanning fluorimetry

To study the protein-ligand binding interactions we used nano differential scanning fluorimetry (NanoDSF). All the samples were prepared in triplicates in the assay buffer (25 mM HEPES pH 7.5, 150 mM NaCl) with a final reaction volume of 15 µL. Samples contained 0.3 mg/ml SARS-CoV-2 Mac1, 10 and 100 µM compounds and ADPr as a control. Melting curves were measured using Prometheus NT.48 (nanoTemper, Germany) with the temperature range from 20 °C to 90 °C with 1°C increment per minute.

### ADP-ribosyl hydrolysis inhibition

For MARylated SRPK2 preparation, 10 μM SRPK2 was incubated with 5 μM PARP10 and 20 μM β-NAD^+^ in reaction buffer (50 mM Tris pH 7.2) at room temperature. After 1 h, 6.7 μM β-NAD^+^ and 1 μM PARP10 were added to the reaction mixture and let incubate at room temperature for 1.5 h. The MARylated SRPK2 was purified using a 1 mL HiTrap IMAC HP column charged with Ni2^+^, followed by buffer exchange to 30 mM HEPES pH 7.5, 500 mM NaCl, 0.5 mM TCEP. To test the hydrolysis activity of SARS-CoV-2 Mac1, the reaction mixtures of 100 nM SARS-CoV-2 Mac1 and 2 µM MARylated SRPK2 in the presence or absence of compounds were incubated at room temperature for 1.5 h. Then, 0.5 μL per spot of the reaction solution was transferred to the nitrocellulose membrane using Echo 650 acoustic liquid dispenser (Labcyte, Sunnyvate, CA). After drying of the spots, the membrane was blocked with 5%(w/v) skimmed milk in TBS-T on a shaker for 15 min. The blocking solution was discarded, and the membrane was incubated with 1 μg/mL Nluc-eAf1521 ^21^ in 1%(w/v) skimmed milk solution for 20 min. Then, the membrane was shortly rinsed and incubated with TBS-T on a shaker for 15 min. After a final rinsing, the membrane was visualized using 500 μL of 1:1000 NanoGlo substrate (Promega, catalogue number: N1110).

### Profiling of macrodomain inhibition

The discovered Mac1 inhibitor **1** and the improved analog **27** were profiled against a panel of human and viral macrodomains and human ARH3 using the FRET assay described above. Assay conditions varied for each enzyme based on the optimizations carried out previously ^21^. In order to efficiently evaluate the selectivity of the compound, it was profiled at 10 µM and 100 µM concentrations. DMSO, 200 µM ADPr or 10 µM untagged-ALC1 controls were included in all the reactions to exclude the effects of DMSO and to calculate the inhibition-%.

### Crystallization, data collection, processing and refinement

The apo crystals for SARS-CoV-2 Mac1 were produced by a sitting drop vapor diffusion method as previously reported ^40^. Protein crystallization was set up in a Swissci 3D 96-well plate. 100 nL of 20 mg/mL SARS-CoV-2 Mac1 was mixed with 80 nL of the crystallization solution using Mosquito pipetting robot (TTP Labtech). The well solution contained the reported crystallization condition: 0.1 M CHES pH 9.5, 32% PEG (v/v) 3000, which was further optimized with varying PEG 3000 concentrations between 28-32% (v/v) to check the reproducibility and to produce co-crystals by micro seeding method. SARS-CoV-2 Mac1 and inhibitors complexes were prepared by mixing 1 mM ligand with 950 µM protein (17.3 mg/ml). Crystallization drops were set up by mixing 100 nL of SARS-CoV-2 Mac1 and ligand solution, 20 nL of seed stock and 80 nL of the reservoir solution. After dispensing the solutions plates were sealed and left for imaging at RT to be equilibrated through vapor diffusion with 40 µL crystallization solution. Crystallization trials were monitored with IceBear ^41^ and co-crystals appeared within 24 h. For cryo-cooling, the crystals were soaked in the crystallization solution supplemented with 10% (v/v) glycerol, 10% MPD and 10% PEG 200 with 0.5 mM of the compounds. X-ray diffraction data were collected on beamline ID30A-1 (Massif-1) at ESRF, Grenoble, France. The dataset was processed by the autoPROC pipeline for **1** ^42^ and using XDS for **27** ^43^(Table S2).

The structures were solved with MOLREP ^44^ by the method of molecular replacement by using SARS-CoV-2 Mac1 (PDB: 6vxs; ^40^) as a search model. Model building and refinement were performed with Coot and REFMAC5, respectively ^45–48^ (Table S2). The structures were visualized in PyMOL version 1.7.2.1 (Schrödinger).

#### Cell culture

Delayed brain tumor (DBT), L929, VeroE6, and HeLa cells expressing the MHV receptor carcinoembryonic antigen-related cell adhesion molecule 1 (CEACAM1) (HeLa-MHVR) were grown in Dulbecco’s Modified Eagle Medium (DMEM) supplemented with 10% fetal bovine serum (FBS), 100 U/ml penicillin and 100 mg/ml streptomycin, HEPES, sodium pyruvate, non-essential amino acids, and L-glutamine. Calu-3 cells (ATCC) were grown in MEM supplemented with 20% FBS. Human IFN-γ was purchased from R&D Systems.

#### Cell viability assay

DBT, L929, and Calu-3 cells were treated with **27** for 24 hours. Cellular metabolic activity was assessed using a CyQUANT MTT Cell Proliferation Assay (Thermo Fisher Scientific) following manufacturer’s instructions.

#### Virus infection

Cells were infected with recombinant MHV-JHM or SARS-CoV-2 at a multiplicity of infection (MOI) of 0.1 PFU/cell with a 45-60 min adsorption phase, unless otherwise stated. Progeny cell-associated and cell-free virus was collected at indicated timepoints and viral titers were determined by plaque assay on Hela-MHVR (MHV) or Vero E6 (SARS-CoV-2) cells. All statistical analysis for viral infections was performed using GraphPad Prism software with an ordinary one-way ANOVA with a Dunnet’s test to correct for multiple comparisons to assess differences between treated and untreated samples. Graphs are expressed as means ± standard errors of the means. The n value represents the number of biological replicates for each group. Significant p values were denoted with asterisks. *, p≤0.05; **, p≤0.01, ***, p≤0.001, ****, p≤0.0001.

#### Identification of drug-resistant mutant viruses

DBT cells were infected in triplicate as described above. After each infection, progeny virus from each individual well was titered and then passaged to a new well of DBT cells. After 6 passages, 2 consecutive plaque picks were performed from 2 of the 3 individually passaged viral samples to collect individual isolates of MHV. RNA was isolated from these isolates using Trizol per manufacturer’s instructions. cDNA was prepared using MMLV-reverse transcriptase per the manufacturer’s instructions (Thermo Fisher Scientific) and PCR was performed using the following primers: Forward - 5’-ggctgttgtggatggcaagca-3’ and Reverse – 5’-gctttggtaccagcaacggag-3’. PCR products were sequenced by Sanger Sequencing (Azenta).

## Data availability

Atomic coordinates and structure factors will be available at the Protein Data Bank with the ids 8TV6 and 8TV7. Raw diffraction data will be available at fairdata.fi (<u>10.23729/3abcfe89-5b16-45cf-9eca-</u> <u>0f4bc26a5beb</u>). All other study data are included in the article and/or supporting information.

## Acknowledgements

The use of the facilities and expertise of the Biocenter Oulu Structural Biology core facility, a member of Biocenter Finland, Instruct-ERIC Centre Finland and FINStruct, are gratefully acknowledged. We are grateful to local contacts at the ESRF for assistance in using beamline ID30A-1. We thank also Heli Alanen for helping with protein production. This work was funded by Sigrid Jusélius foundation (for LL), Emil Aaltonen foundation (for TP), the University of Oulu proof-of-concept project (for LL), National Institutes of Health (NIH) grants P20 GM113117, R35GM138029, and P30GM110761 (a CTSA grant from NCATS awarded to the University of Kansas for Frontiers: University of Kansas Clinical and Translational Science Institute (#UL1TR002366)) (for ARF).

