## Supplementary figures and tables for "Discovery of 2-amide-3-methylester thiophenes that target SARS-CoV-2 Mac1 and repress coronavirus replication, validating Mac1 as an anti-viral target"

**CONTENT**

**Figure S1:** $IC_{50}$ measurements of hit compounds.
**Figure S2:** Thermal shift assay.
**Figure S3:** Cell viability assays.
**Figure S4:** Comparison of Mac1 and MacroD2 crystal structures.
**Table S1.** Assay validation statistics.
**Table S2.** Hit validation and counter screening.
**Table S3.** Crystallography data and refinement statistics.
**Figure S4-35.** NMR spectra
**Figure S36.** HPLC chromatogram of **27** at 220nm detection wavelength.
**Figure S37.** HPLC chromatogram of **27** at 260nm detection wavelength.

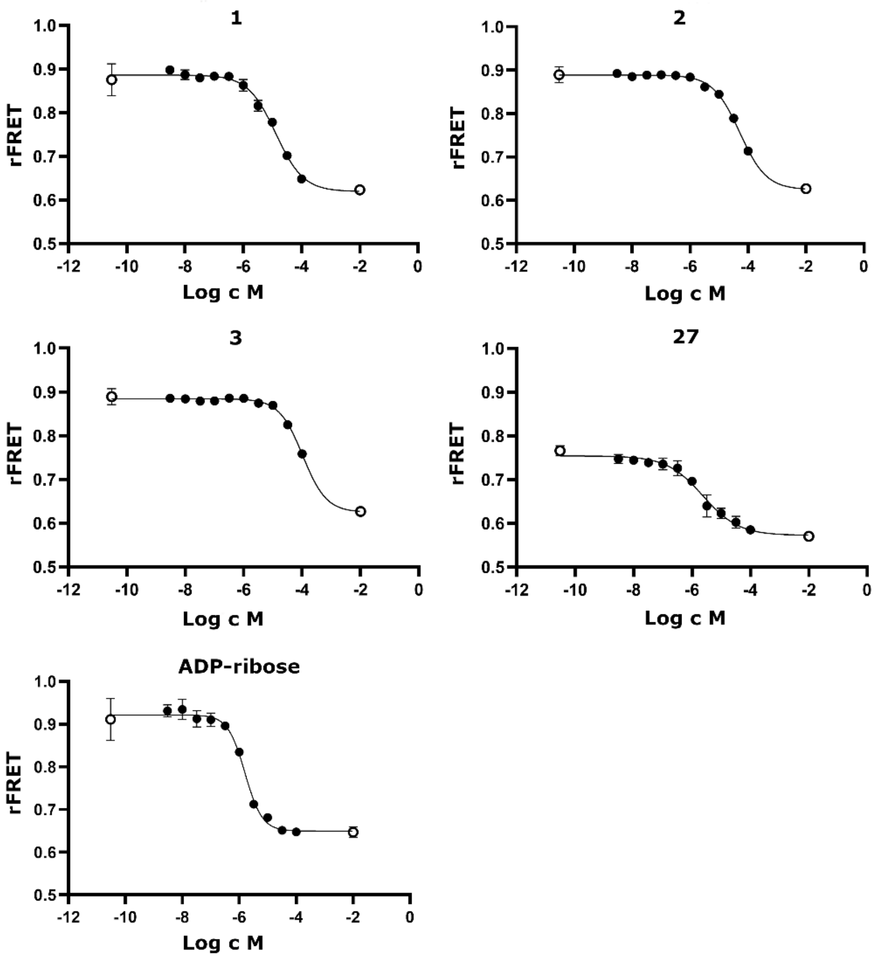


**Figure S1.** Representative $IC_{50}$ measurements of SARS-CoV-2 Mac1 for ADPr and initial hit compounds. Measurements were carried out in half-logarithmic dilution starting from 100 to 0.003 µM for each curve. Negative control (containing no compound) was set one logarithmic unit below the lowest concentration. positive control (containing 200 µM ADPr) was set one logarithmic unit above the highest concentration of the compound. Data is reported as mean ± SD of four replicates.

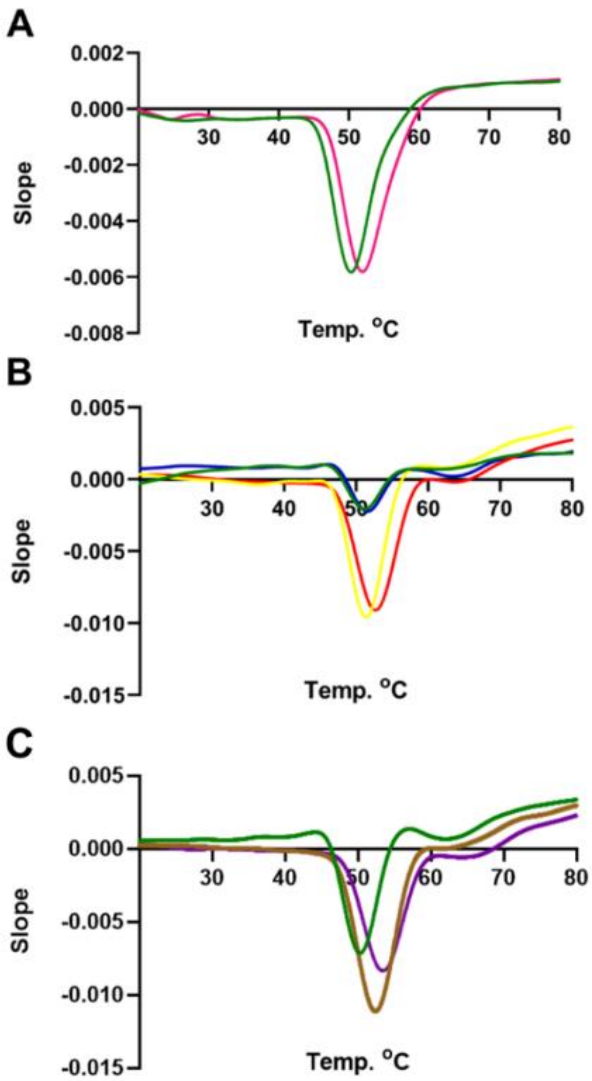


**Figure S2.** Thermal shift assay of SARS-CoV-2 Mac1 for ADPr and initial hit compounds and the most potent compound **27**. (A) buffer control (green). ADPr (pink) (B) DMSO control (green). **1** (Red). **2** (Yellow). **3** (blue) (C) DMSO control (green). 30 µM **27** (brown), 60 µM **27** (purple). Each curve was measured three times, and a representative second derivative curve is shown.

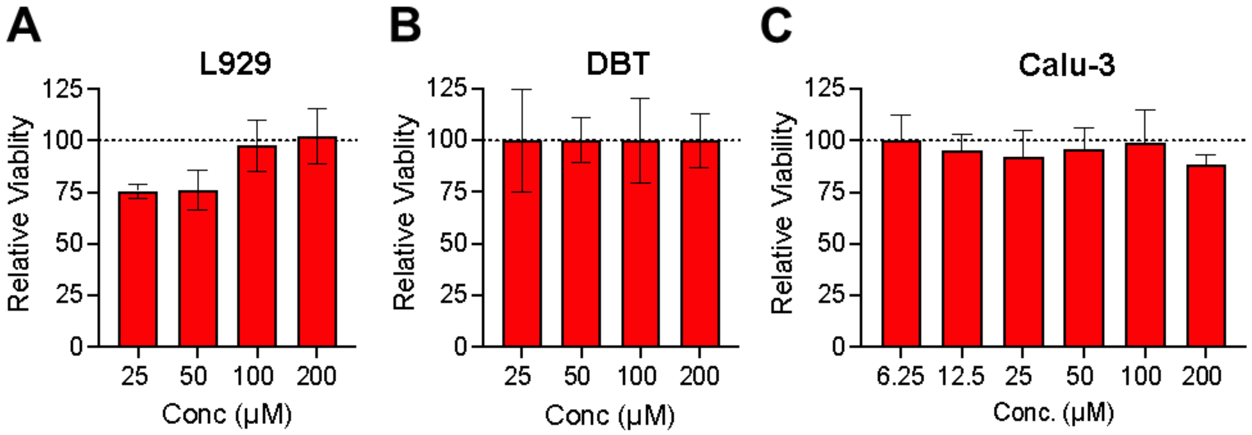


**Figure S3.** Compound **27** does not affect the viability of mouse or human cell lines. (A-B) L929 (A), DBT (B) and Calu-3 (C) cells were treated with compound **27** at the indicated concentrations for 24 hours and then viability was measured by an MTT assay and normalized to the viability level of DMSO treated cells. The results in A and B are representative of 3 independent experiments, while the results in C are the combined results of 3 independent experiments. n=3.

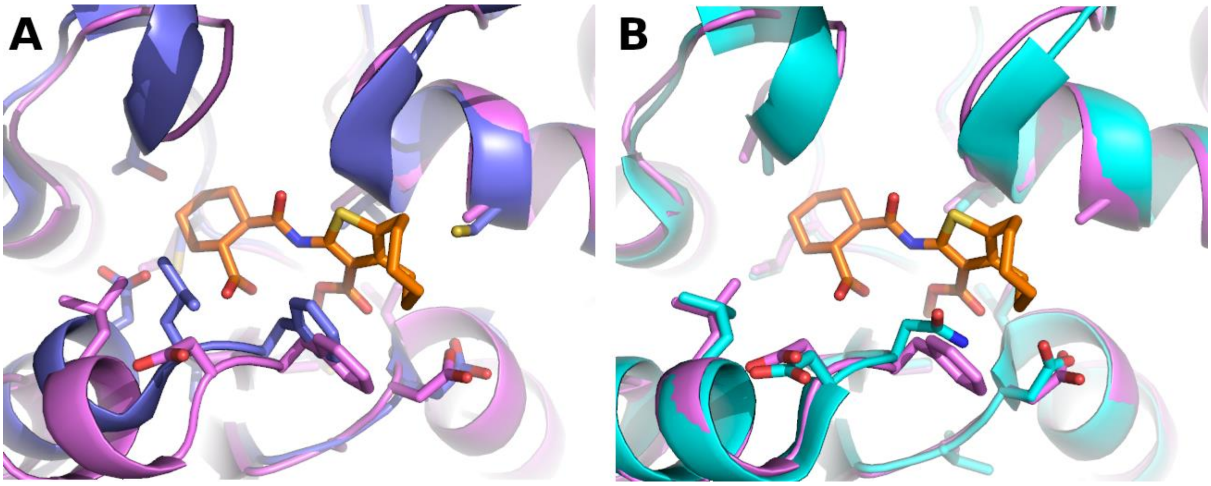


**Figure S4.** Comparison of macrodomain crystal structures. (A) Superimposed SARS-CoV-2 Mac1 in complex with **27** (PDB: 8TV7, orange) and MacroD2 (PDB: 6Y4Y, blue). (B) SARS-CoV-2 Mac1 in complex with **27** and SARS-CoV Mac1 (PDB: 2ACF, cyan). Binding site residues and **27** is shown in sticks (orange).

**Table S1.** Assay validation statistics.

| Statistical parameters | FRET assay |
|---|---|
| S/B | 1.33 ± 0.03 |
| S/N | 22.04 ± 2.48 |
| Z' | 0.81 ± 0.02 |
| Well-to-well CV (max/min. %) | 0.95 ± 0.22 / 0.82 ± 0.19 |
| Plate-to-plate CV (%)* | 0.33 |
| Day-to-day CV (%)* | 2.55 |

*Calculated from Z' values.

**Table S2.** Hit validation and counter screening results with TNKS ARC4 assay.

| ID | Figure | TNKS2 ARC4 | NanoDSF ΔTm °C | Inhibition % primary screening | | $IC_{50}$ ($pIC_{50}$ ± SEM), n=3 |
|---|---|---|---|---|---|---|
| | | | | 410 nm | 430 nm | |
| **1** | 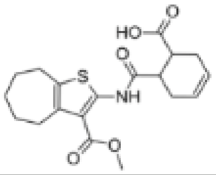 | NI* | 1.0 | 60.5 | 73.3 | 14 µM (-4.85 ± 0.02) |
| **2** | 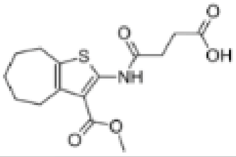 | NI | 0.5 | 23.3 | 40.6 | 48 µM (-4.32 ± 0.02) |
| **3** | 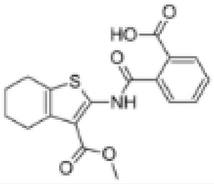 | NI | 1.0 | 29.2 | 31.8 | 110 µM (-3.96 ± 0.02) |
| **4** | 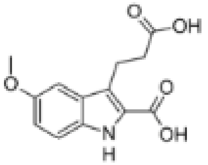 | NI | 0.3 | 37.6 | 39.5 | 89 µM (-4.05 ± 0.01) |
| **5** | 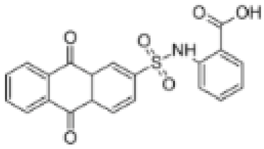 | NI | 0.1 | 32.3 | 30.5 | 45 µM (-4.34 ± 0.03) |
| **6** | 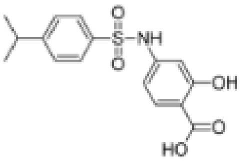 | NI | 1.0 | 36.0 | 48.9 | 57 µM (-4.24 ± 0.03) |
| **7** | 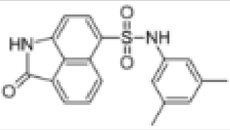 | NI | 0.1 | 64.5 | 59.7 | NI |
| **8** | 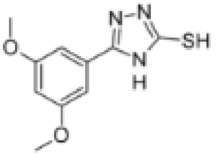 | NI | 0.3 | 75.5 | 77.5 | 57 µM ( -4.26 ± 0.07) |

*NI. no inhibition

**Table S3.** Data collection and refinement statistics for co-crystal structures.

| **Inhibitor (PDB id.)** | **1 (8TV6)** | **27 (8TV7)** |
|---|---|---|
| **Data collection** | | |
| Beamline | ID30A-1 ESRF | ID30A-1 ESRF |
| Wavelength (Å) | 0.9655 | 0.9655 |
| Space group | P1 | $P2_12_12_1$ |
| Unit cell dimensions | | |
| *a*, *b*, *c* (Å) | 30.20, 37.80, 64.10 | 38.0, 55.0, 71.9 |
| α, β, γ (°) | 95.60, 98.10, 90.40 | 90.0, 90.0, 90.0 |
| Resolution range (Å) | 37.58-1.74 (1.80-[illegible] | 43.72-1.50 (1.53-1.50) |
| Total no. of reflections | 28635 (27298) | 132023 (10008) |
| No. of unique reflections | 27365 (1499) | 25009 (1245) |
| Completeness (%) | 95.4 (94.6) | 99.9 (99.9) |
| $I/\sigma(I)$ | 7.3 (2.1) | 9.8 (1.7) |
| CC1/2 (%) | 99.0 (76.0) | 99.7 (64.1) |
| $R_{meas}$ | 10.5 (46.7) | 11.5 (92.5) |
| **Model building and refinement** | | |
| R-factor | 17.4 | 17.0 |
| R-free | 22.4 | 19.0 |
| No. of atoms | | |
| Protein | 2580 | 1363 |
| Ligands* | 46 | 39 |
| Water | 220 | 146 |
| RMSD | | |
| Bonds (Å) | 0.0075 | 0.0124 |
| Angles (°) | 0.1441 | 0.1590 |
| Average *B* factors (Å$^2$) | 23.03 | 17.39 |
| Protein | 15.27 | 12.94 |
| Ligands* | 31.11 | 15.44 |
| Water | 22.73 | 23.80 |
| Ramachandran plot | | |
| Favoured (%) | 100 | 94 |
| Allowed (%) | 0.0 | 6.0 |
| Outliers (%) | 0.0 | 0.0 |

[1]Values within parentheses refers to the highest resolution shell. *Inhibitor, PEG and glycerol molecules.

**Figure S4.** $^{1}H$ NMR spectrum of compound **9a** in $CDCl_3$ (-0.5–15 ppm).

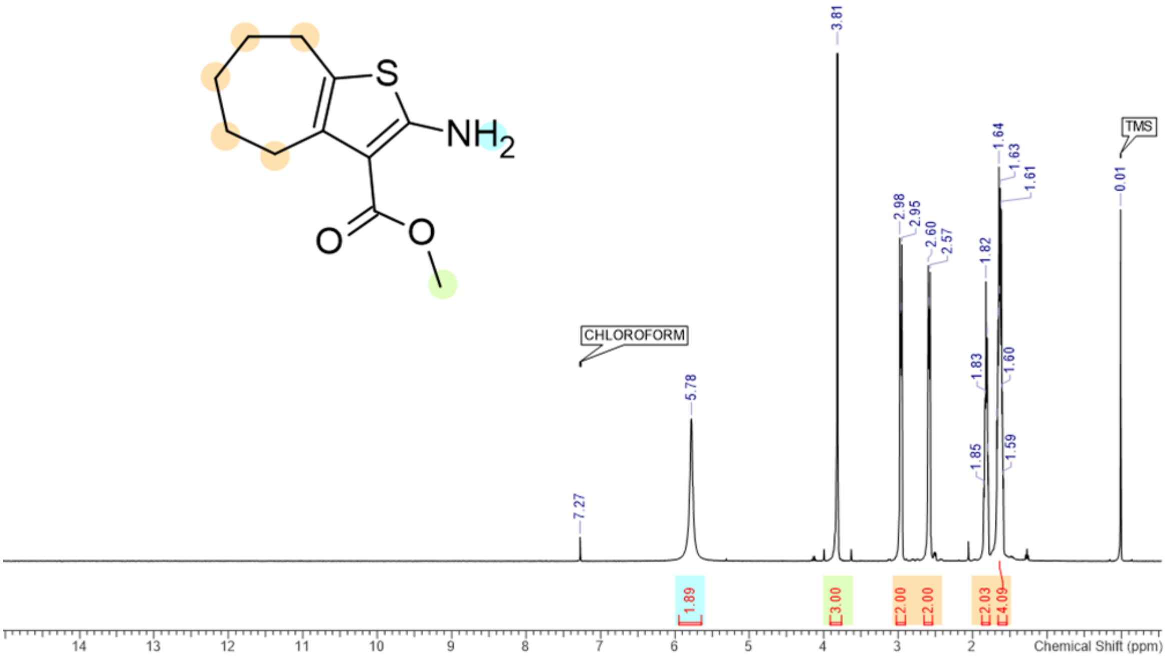


**Figure S5.** $^{1}H$ NMR spectrum of compound **9b** in $CDCl_3$ (-0.5–15 ppm).

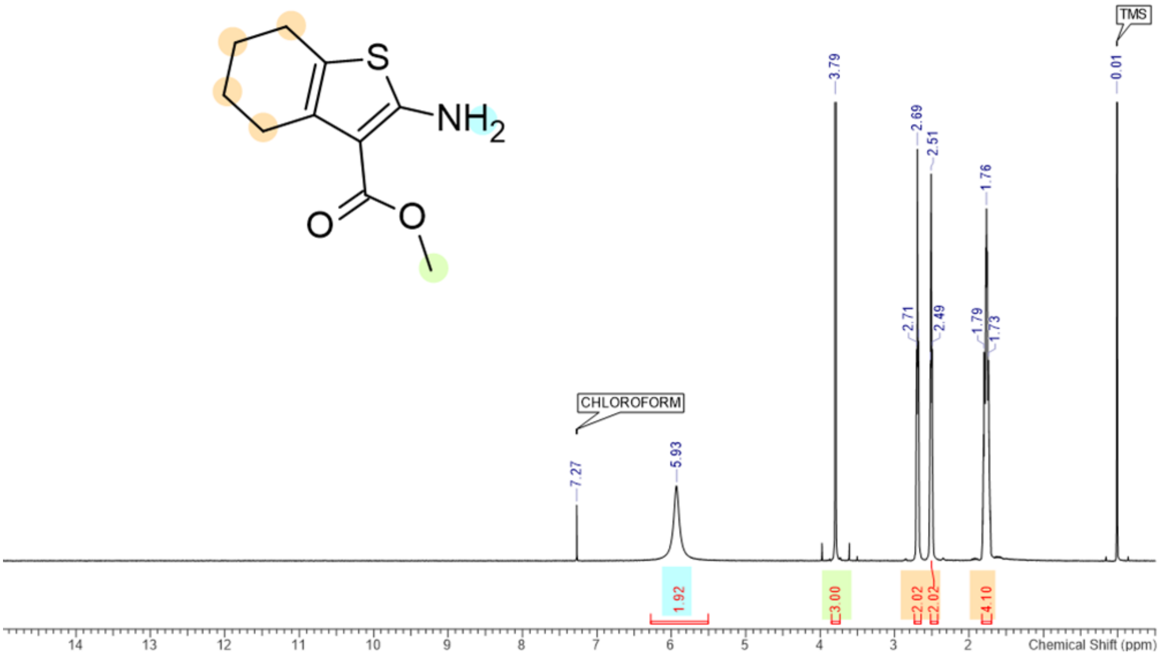

**Figure S6.** [1]H NMR spectrum of compound **9c** in $CDCl_3$ (-0.5–15 ppm).

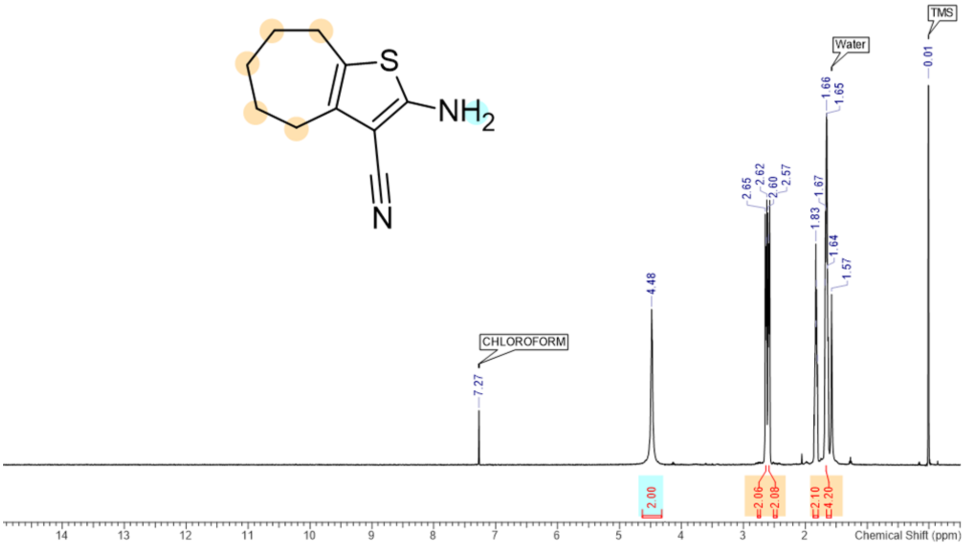


**Figure S7.** [1]H NMR spectrum of compound **9d** in $CDCl_3$ (-0.5–15 ppm).

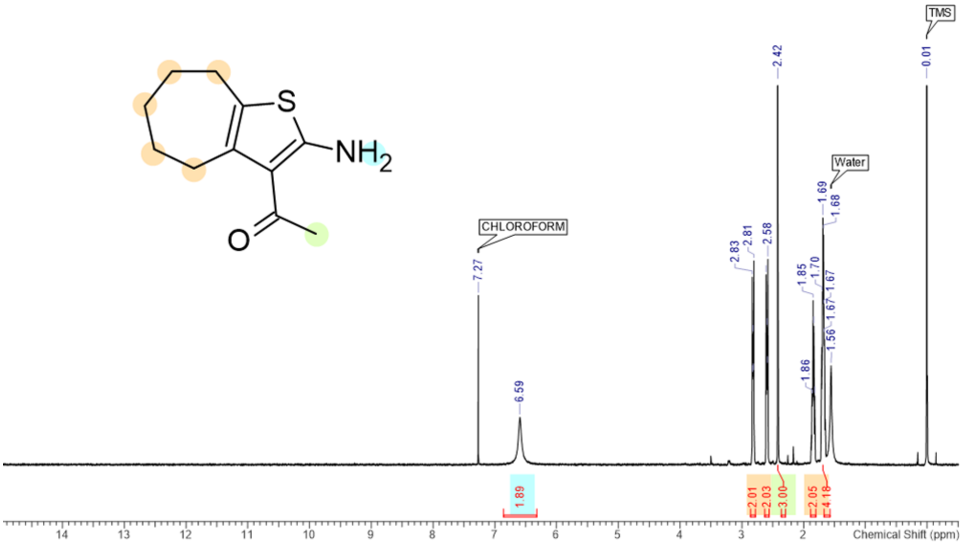

**Figure S8.** $^{1}$H NMR spectrum of compound **9e** in $d_6$-DMSO (-0.5–15 ppm).

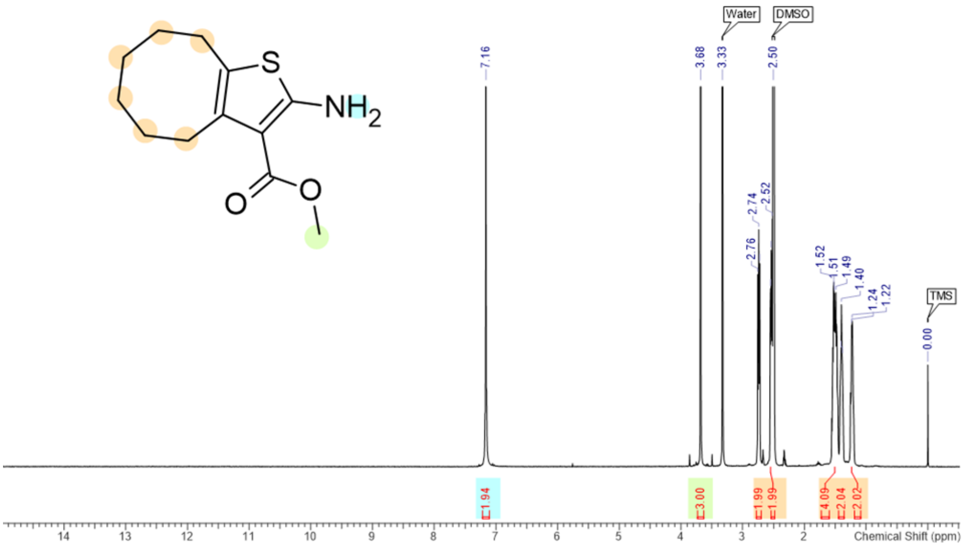


**Figure S9.** $^{1}$H NMR spectrum of compound **1r** in $CDCl_3$ (-0.5–15 ppm).

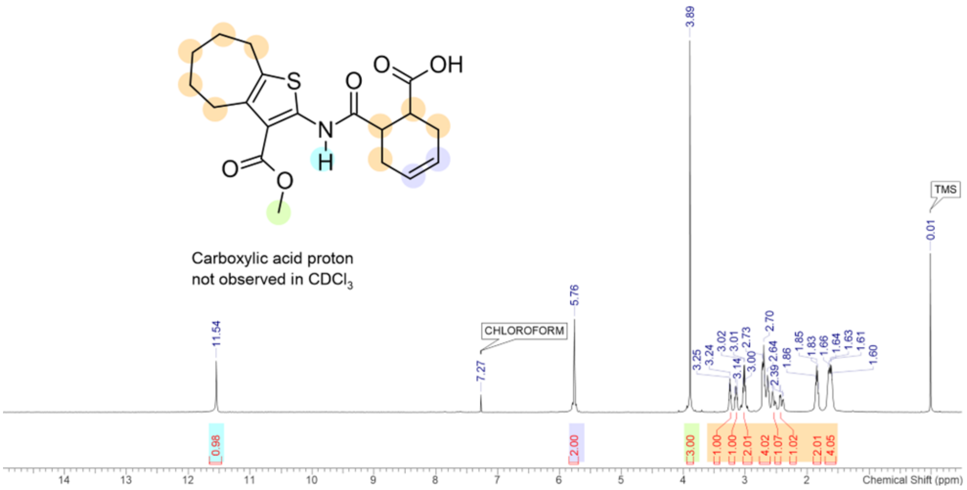

**Figure S10.** $^{1}$H NMR spectrum of compound **1r** in $d_6$-DMSO (-0.5–15 ppm).

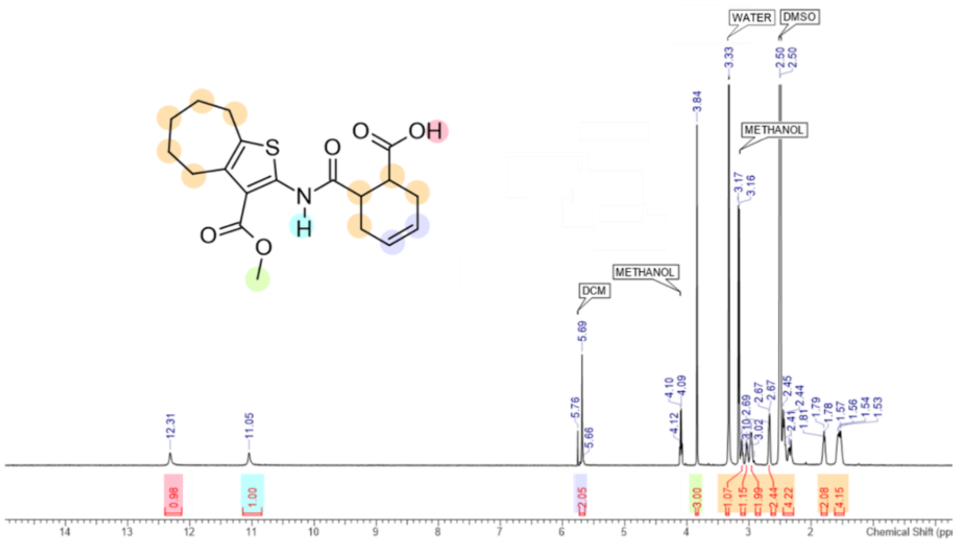


**Figure S11.** $^{13}$C NMR spectrum of compound **1r** in $CDCl_3$ (0–220 ppm)

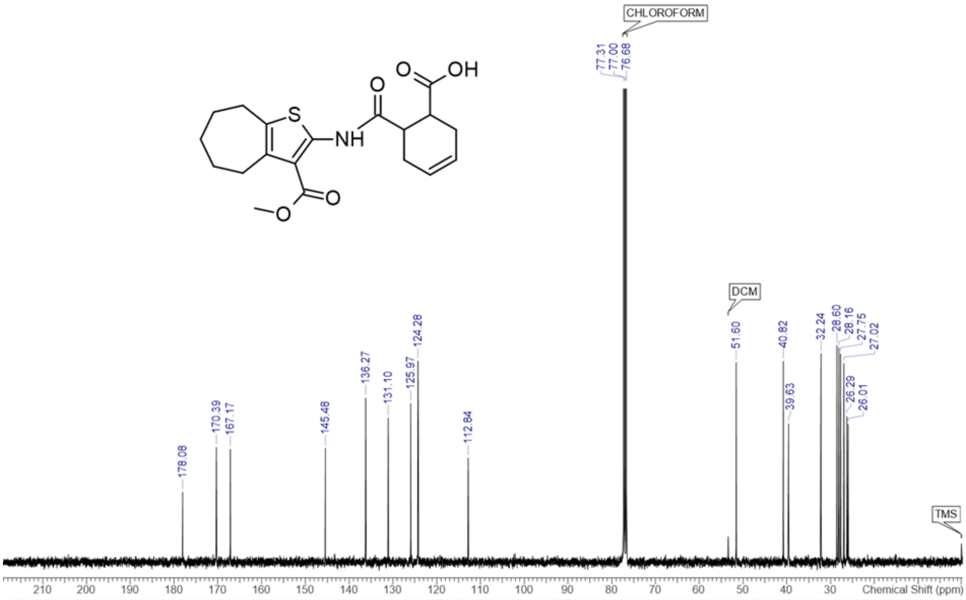


| No. | (ppm) | (Hz) | Height | No. | (ppm) | (Hz) | Height | No. | (ppm) | (Hz) | Height |
|---|---|---|---|---|---|---|---|---|---|---|---|
| 1 | 26.01 | 2617.2 | 0.1200 | 8 | 39.63 | 3987.5 | 0.1206 | 15 | 136.27 | 13710.6 | 0.1429 |
| 2 | 26.29 | 2645.0 | 0.1268 | 9 | 40.82 | 4107.1 | 0.1748 | 16 | 145.48 | 14637.1 | 0.0992 |
| 3 | 27.02 | 2718.4 | 0.1729 | 10 | 51.60 | 5192.1 | 0.1739 | 17 | 167.17 | 16819.6 | 0.0984 |
| 4 | 27.75 | 2791.7 | 0.1813 | 11 | 112.84 | 11352.8 | 0.0909 | 18 | 170.39 | 17143.1 | 0.1000 |
| 5 | 28.16 | 2833.6 | 0.1864 | 12 | 124.28 | 12503.8 | 0.1750 | 19 | 178.08 | 17917.0 | 0.0611 |
| 6 | 28.60 | 2877.6 | 0.1886 | 13 | 125.97 | 12674.0 | 0.1382 | | | | |
| 7 | 32.24 | 3243.6 | 0.1815 | 14 | 131.10 | 13190.5 | 0.1255 | | | | |

**Figure S12.** $^{1}$H NMR spectrum of compound **3r** in $d_6$-DMSO (-0.5–15 ppm).

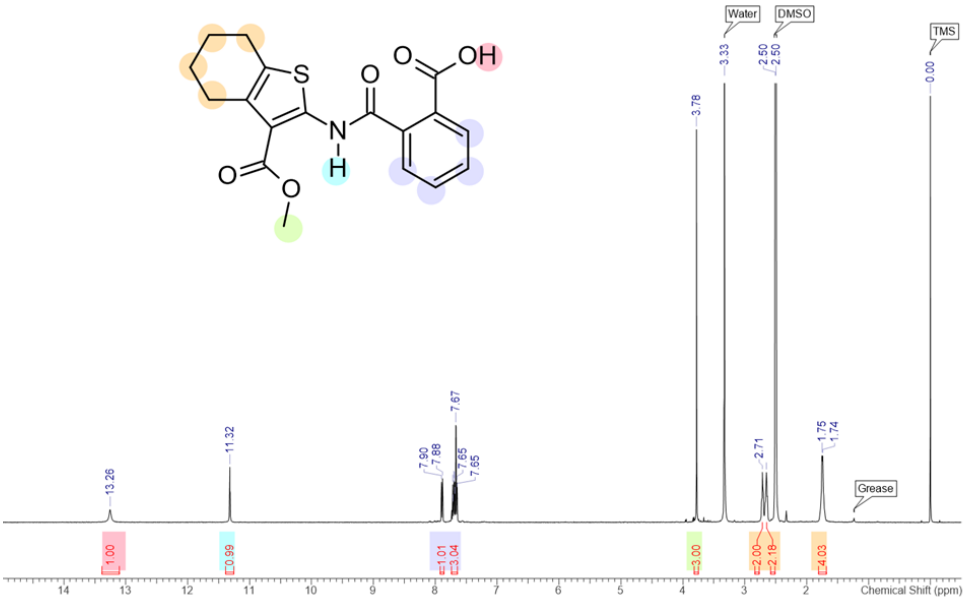


**Figure S13.** $^{13}$C NMR spectrum of compound **3r** in $CDCl_3$ (0–220 ppm).

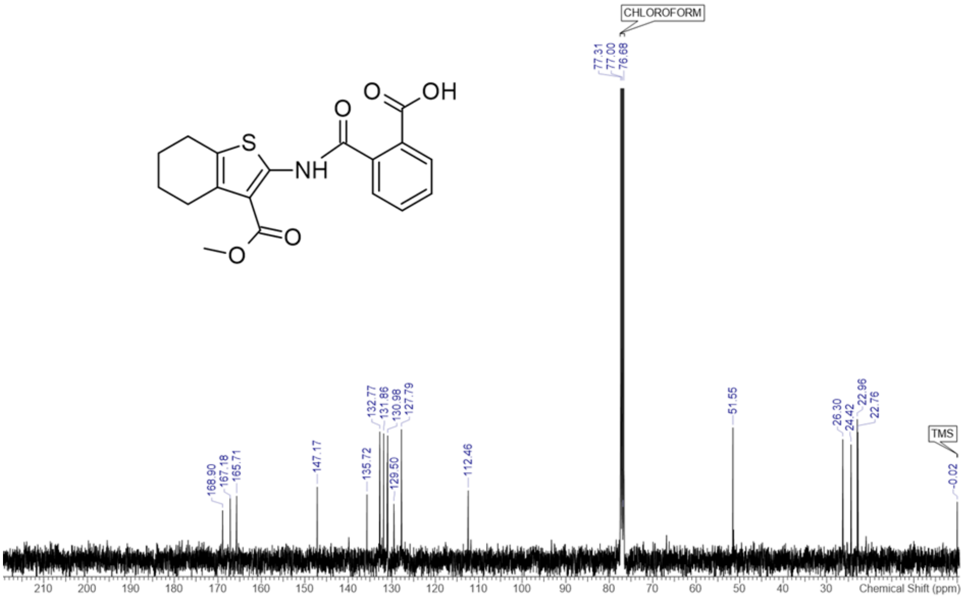


| No. | (ppm) | (Hz) | Height | No. | (ppm) | (Hz) | Height | No. | (ppm) | (Hz) | Height |
|---|---|---|---|---|---|---|---|---|---|---|---|
| 1 | 22.76 | 2290.0 | 0.0375 | 7 | 127.75 | 12853.7 | 0.0236 | 13 | 132.77 | 13358.5 | 0.0377 |
| 2 | 22.96 | 2309.8 | 0.0413 | 8 | 127.79 | 12857.4 | 0.0383 | 14 | 135.72 | 13655.6 | 0.0193 |
| 3 | 24.42 | 2456.5 | 0.0339 | 9 | 129.50 | 13029.1 | 0.0165 | 15 | 147.17 | 14807.3 | 0.0215 |
| 4 | 26.30 | 2645.8 | 0.0354 | 10 | 130.98 | 13178.0 | 0.0365 | 16 | 165.71 | 16672.1 | 0.0189 |
| 5 | 51.55 | 5186.9 | 0.0389 | 11 | 131.06 | 13186.8 | 0.0287 | 17 | 167.18 | 16820.3 | 0.0172 |
| 6 | 112.46 | 11314.7 | 0.0204 | 12 | 131.86 | 13266.8 | 0.0371 | 18 | 168.90 | 16993.4 | 0.0146 |

**Figure S14.** $^{1}$H NMR spectrum of compound **10** in $d_6$-DMSO (-0.5–15 ppm).

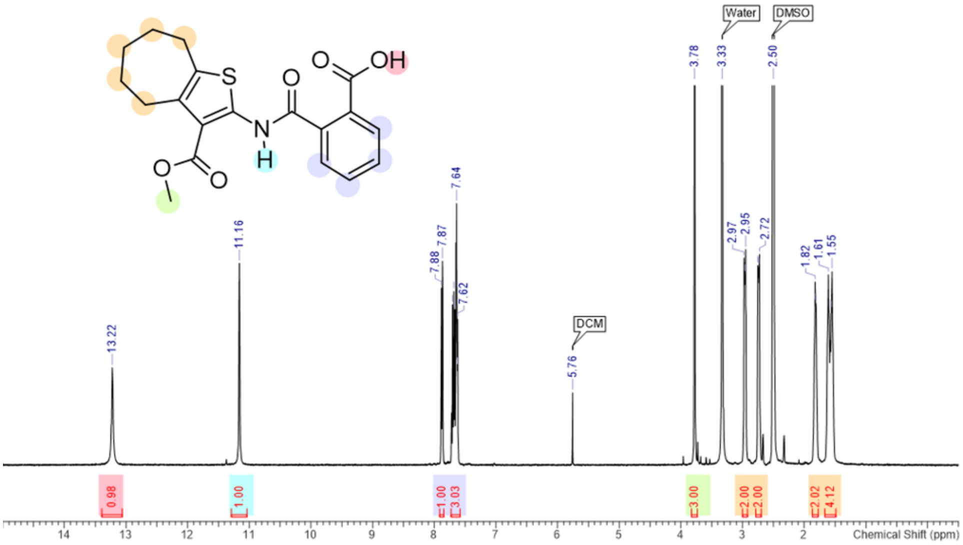


**Figure S15.** $^{13}$C NMR spectrum of compound **10** in $CDCl_3$ (0–220 ppm).

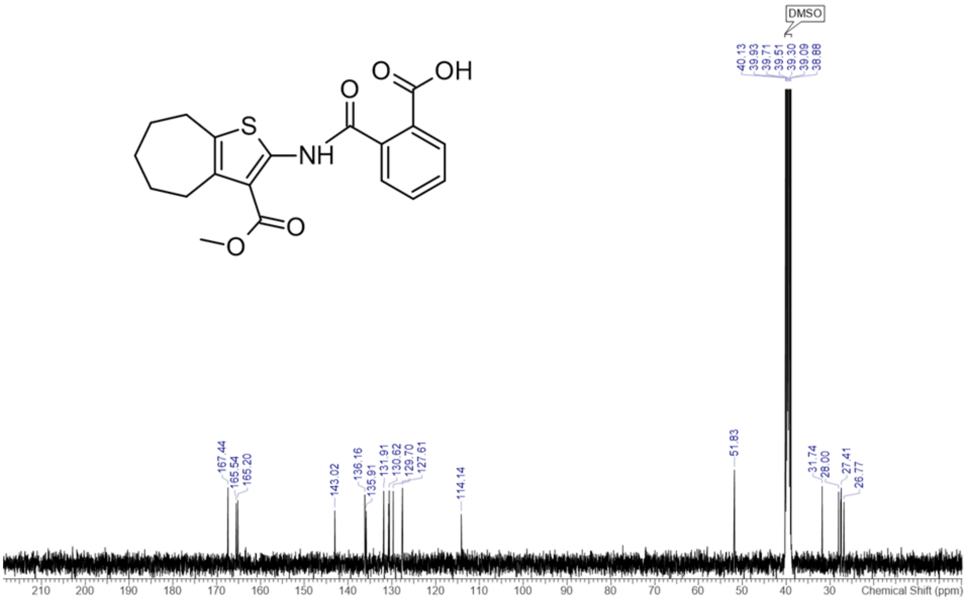


| No. | (ppm) | (Hz) | Height |
|---|---|---|---|
| 1 | 26.77 | 2693.6 | 0.0151 |
| 2 | 27.41 | 2757.4 | 0.0185 |
| 3 | 27.50 | 2767.0 | 0.0150 |
| 4 | 28.00 | 2816.9 | 0.0176 |
| 5 | 31.74 | 3193.2 | 0.0189 |
| 6 | 51.83 | 5214.3 | 0.0229 |
| 7 | 114.14 | 11484.3 | 0.0122 |
| 8 | 127.61 | 12839.2 | 0.0185 |
| 9 | 129.70 | 13049.8 | 0.0177 |
| 10 | 130.62 | 13142.2 | 0.0179 |
| 11 | 130.77 | 13156.9 | 0.0147 |
| 12 | 130.84 | 13164.2 | 0.0092 |
| 13 | 131.91 | 13272.1 | 0.0178 |
| 14 | 135.91 | 13674.1 | 0.0129 |
| 15 | 136.16 | 13699.8 | 0.0169 |
| 16 | 143.02 | 14389.3 | 0.0130 |
| 17 | 165.20 | 16620.9 | 0.0155 |
| 18 | 165.54 | 16655.4 | 0.0148 |
| 19 | 167.44 | 16846.9 | 0.0186 |

**Figure S16.** $^{1}$H NMR spectrum of compound **11** in $CDCl_3$ (-0.5–15 ppm).

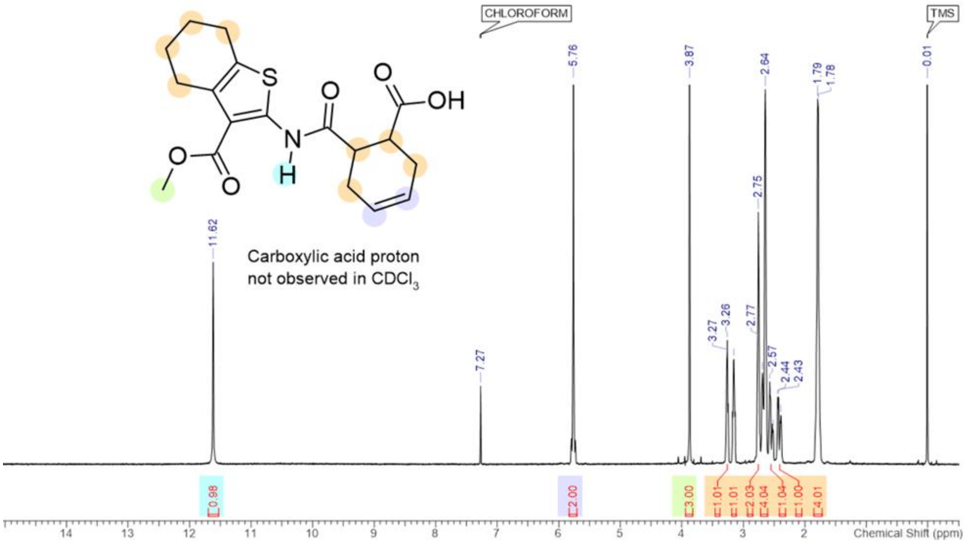


**Figure S17.** $^{13}$C NMR spectrum of compound **11** in $CDCl_3$ (0–220 ppm).

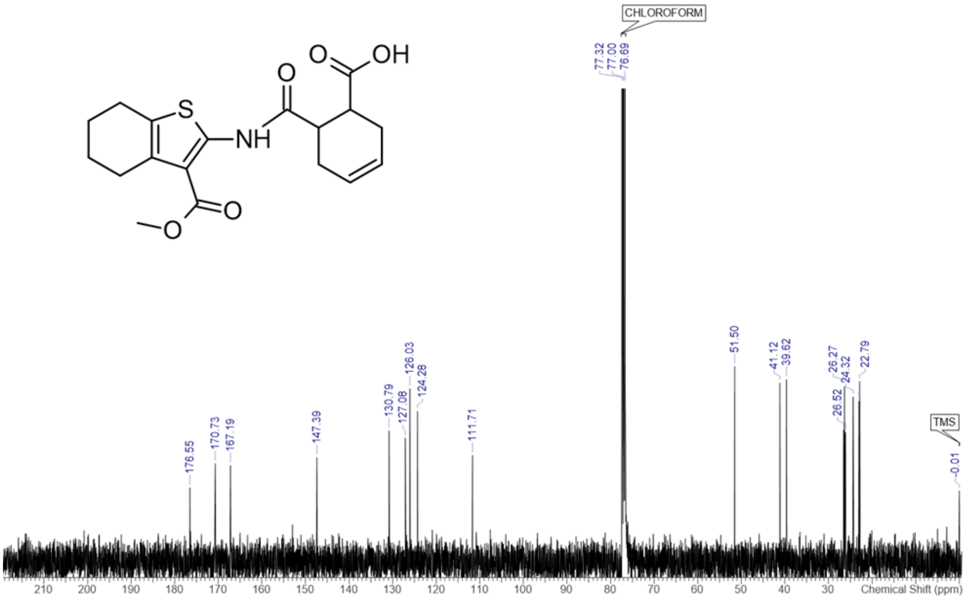


| No. | (ppm) | (Hz) | Height | No. | (ppm) | (Hz) | Height | No. | (ppm) | (Hz) | Height |
|---|---|---|---|---|---|---|---|---|---|---|---|
| 1 | 22.79 | 2292.9 | 0.0366 | 7 | 39.62 | 3986.8 | 0.0369 | 13 | 127.08 | 12786.3 | 0.0251 |
| 2 | 22.96 | 2309.8 | 0.0325 | 8 | 41.12 | 4137.2 | 0.0362 | 14 | 130.79 | 13158.9 | 0.0265 |
| 3 | 24.32 | 2447.0 | 0.0334 | 9 | 51.50 | 5181.8 | 0.0396 | 15 | 147.39 | 14829.3 | 0.0211 |
| 4 | 26.11 | 2626.7 | 0.0263 | 10 | 111.71 | 11239.1 | 0.0216 | 16 | 167.19 | 16821.0 | 0.0195 |
| 5 | 26.27 | 2643.6 | 0.0355 | 11 | 124.28 | 12504.6 | 0.0305 | 17 | 170.73 | 17177.6 | 0.0199 |
| 6 | 26.52 | 2667.8 | 0.0267 | 12 | 126.03 | 12679.9 | 0.0351 | 18 | 176.55 | 17763.0 | 0.0150 |

**Figure S18.** $^{1}$H NMR spectrum of compound **15r** in $d_6$-DMSO (-0.5–15 ppm).

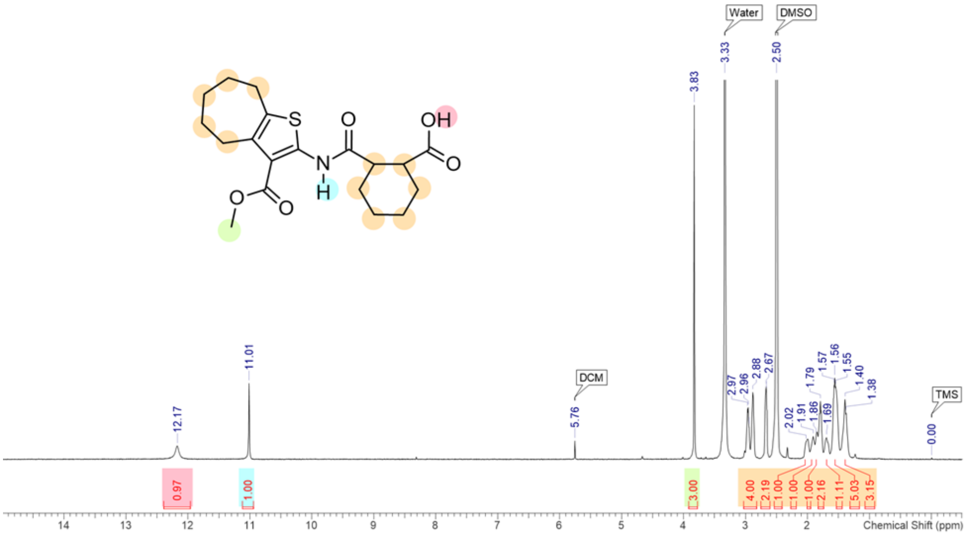


**Figure S19.** $^{13}$C NMR spectrum of compound **15r** in $d_6$-DMSO (0–220 ppm).

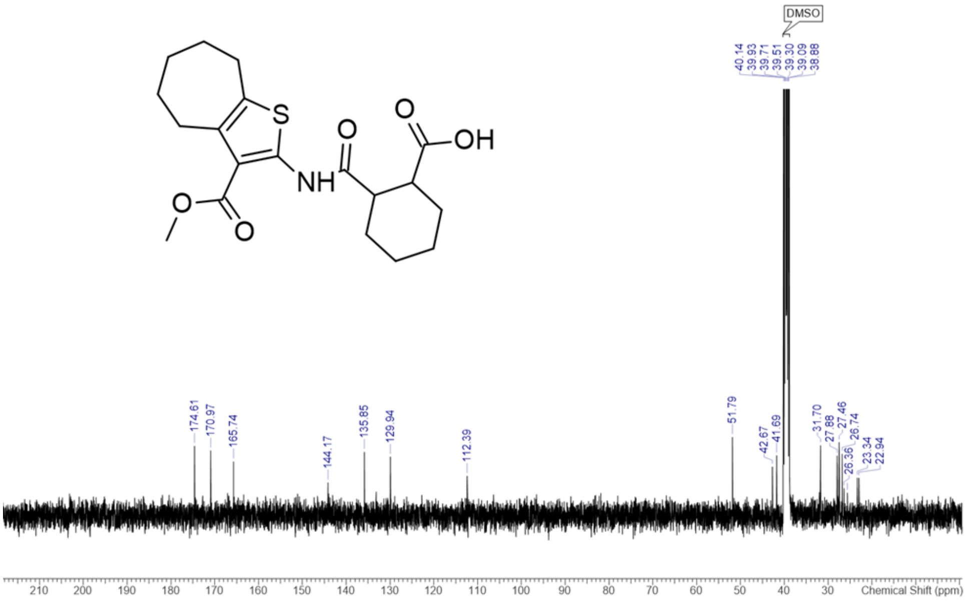


| No. | (ppm) | (Hz) | Height |
|---|---|---|---|
| 1 | 22.94 | 2307.8 | 0.0076 |
| 2 | 23.34 | 2348.1 | 0.0076 |
| 3 | 25.56 | 2571.8 | 0.0046 |
| 4 | 26.36 | 2651.8 | 0.0055 |
| 5 | 26.74 | 2690.0 | 0.0124 |
| 6 | 27.41 | 2758.2 | 0.0139 |
| 7 | 27.46 | 2763.3 | 0.0148 |
| 8 | 27.88 | 2805.1 | 0.0121 |
| 9 | 31.70 | 3189.5 | 0.0141 |
| 10 | 41.69 | 4194.6 | 0.0121 |
| 11 | 42.67 | 4292.9 | 0.0098 |
| 12 | 51.79 | 5210.6 | 0.0158 |
| 13 | 112.39 | 11308.2 | 0.0080 |
| 14 | 129.94 | 13073.3 | 0.0118 |
| 15 | 135.85 | 13668.2 | 0.0128 |
| 16 | 144.17 | 14505.2 | 0.0066 |
| 17 | 165.74 | 16675.2 | 0.0109 |
| 18 | 170.97 | 17201.9 | 0.0131 |
| 19 | 174.61 | 17568.0 | 0.0140 |

**Figure S20.** $^{1}H$ NMR spectrum of compound **21** in $d_6$-DMSO (-0.5–15 ppm).

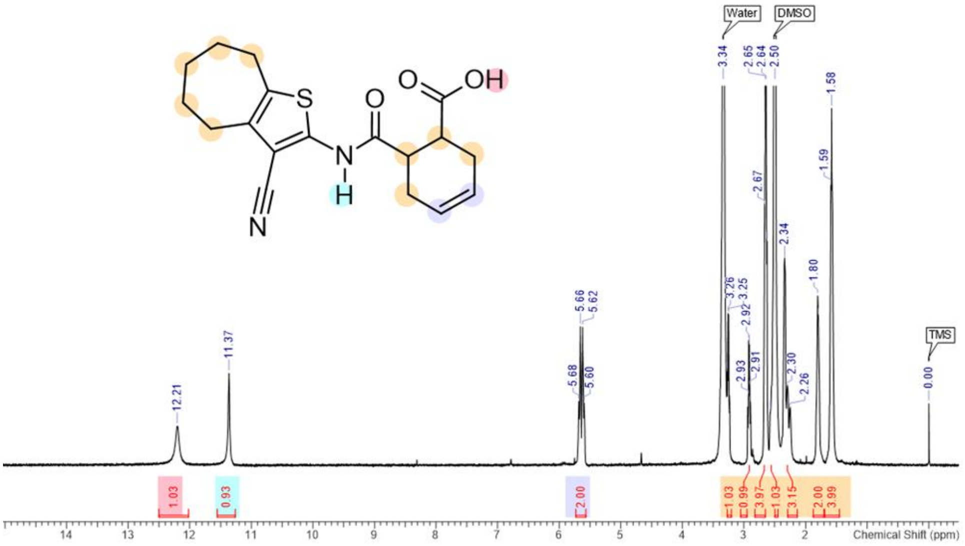


**Figure S21.** $^{13}C$ NMR spectrum of compound **21** in $CDCl_3$ (0–220 ppm).

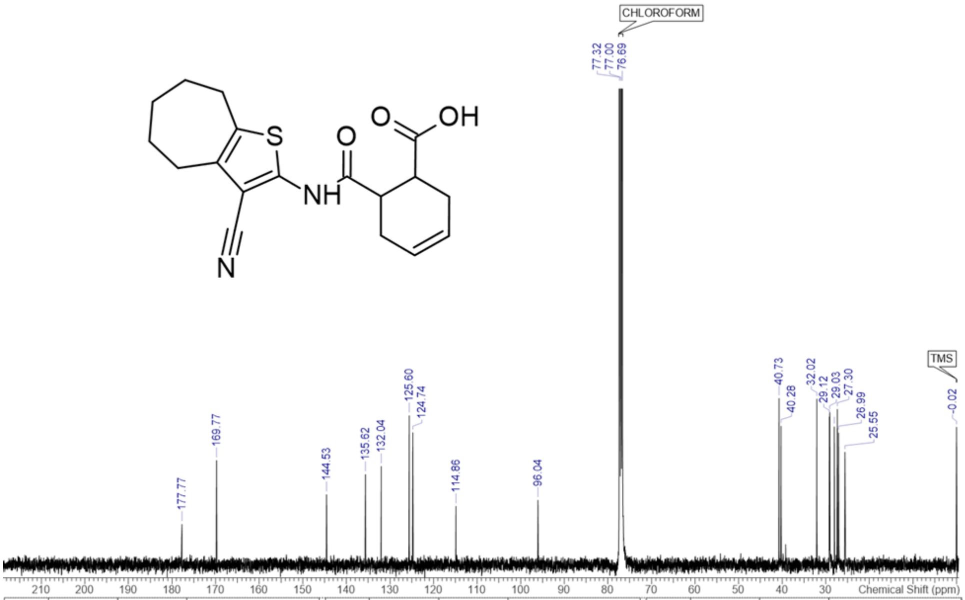


| No. | (ppm) | (Hz) | Height | No. | (ppm) | (Hz) | Height | No. | (ppm) | (Hz) | Height |
|---|---|---|---|---|---|---|---|---|---|---|---|
| 1 | 25.55 | 2570.9 | 0.0131 | 7 | 32.02 | 3221.6 | 0.0193 | 13 | 125.60 | 12636.6 | 0.0174 |
| 2 | 26.99 | 2715.5 | 0.0154 | 8 | 40.28 | 4052.8 | 0.0162 | 14 | 132.04 | 13285.1 | 0.0115 |
| 3 | 27.30 | 2747.0 | 0.0181 | 9 | 40.73 | 4098.3 | 0.0194 | 15 | 135.62 | 13645.3 | 0.0105 |
| 4 | 28.00 | 2817.4 | 0.0161 | 10 | 96.04 | 9662.6 | 0.0075 | 16 | 144.53 | 14541.7 | 0.0082 |
| 5 | 29.03 | 2920.9 | 0.0178 | 11 | 114.86 | 11556.0 | 0.0068 | 17 | 169.77 | 17081.5 | 0.0122 |
| 6 | 29.12 | 2929.7 | 0.0173 | 12 | 124.74 | 12550.0 | 0.0154 | 18 | 177.77 | 17886.2 | 0.0047 |

**Figure S22.** $^{1}$H NMR spectrum of compound **22** in $d_6$-DMSO (-0.5–15 ppm).

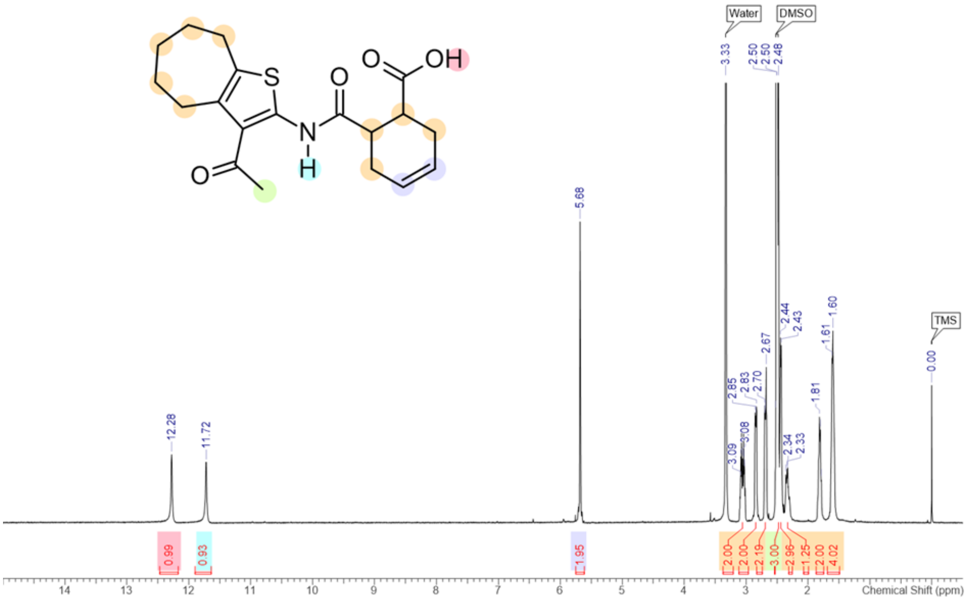


**Figure S23.** $^{13}$C NMR spectrum of compound **22** in $CDCl_3$ (0–220 ppm).

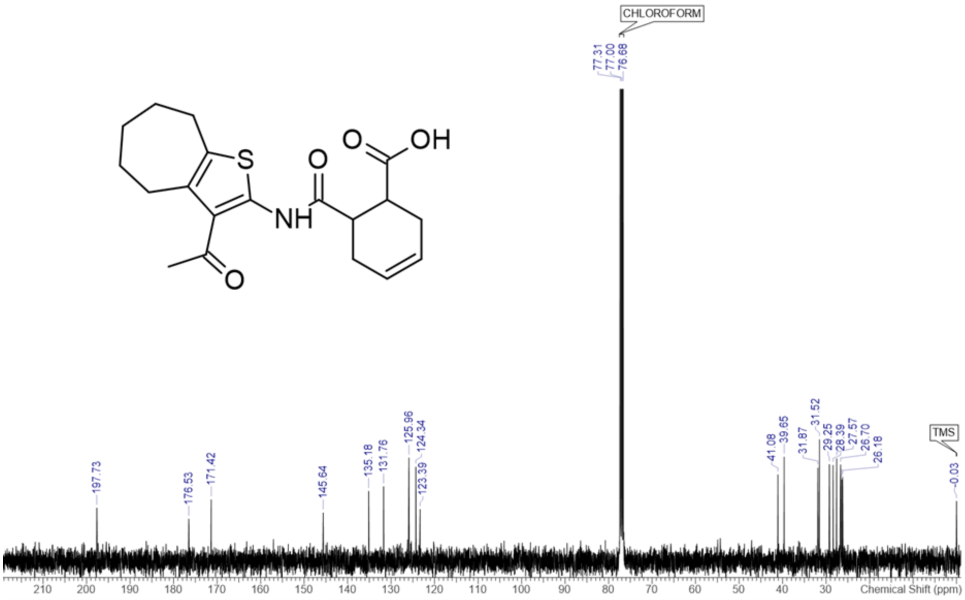


| No. | (ppm) | (Hz) | Height | No. | (ppm) | (Hz) | Height | No. | (ppm) | (Hz) | Height |
|---|---|---|---|---|---|---|---|---|---|---|---|
| 1 | 26.18 | 2634.0 | 0.0296 | 8 | 31.87 | 3206.2 | 0.0328 | 15 | 135.18 | 13600.5 | 0.0246 |
| 2 | 26.35 | 2651.6 | 0.0287 | 9 | 39.65 | 3989.0 | 0.0366 | 16 | 145.64 | 14653.3 | 0.0170 |
| 3 | 26.70 | 2686.1 | 0.0339 | 10 | 41.08 | 4133.5 | 0.0304 | 17 | 171.42 | 17247.2 | 0.0216 |
| 4 | 27.57 | 2774.1 | 0.0360 | 11 | 123.39 | 12415.1 | 0.0182 | 18 | 176.53 | 17761.5 | 0.0149 |
| 5 | 28.39 | 2856.3 | 0.0337 | 12 | 124.34 | 12510.4 | 0.0331 | 19 | 197.73 | 19894.1 | 0.0187 |
| 6 | 29.25 | 2942.9 | 0.0340 | 13 | 125.96 | 12673.3 | 0.0363 | | | | |
| 7 | 31.52 | 3171.0 | 0.0427 | 14 | 131.76 | 13256.5 | 0.0262 | | | | |

**Figure S24.** $^{1}$H NMR spectrum of compound **23** in $d_6$-DMSO (-0.5–15 ppm).

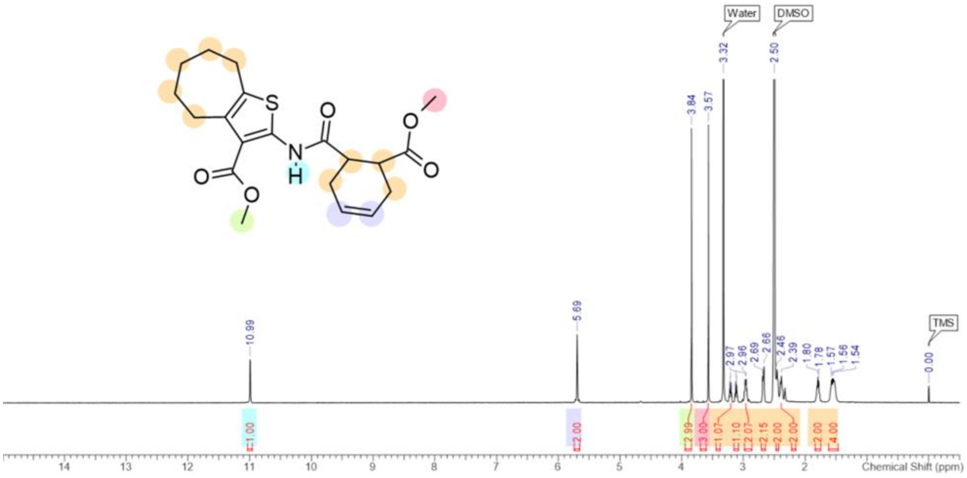


**Figure S25.** $^{13}$C NMR spectrum of compound **23** in $CDCl_3$ (0–220 ppm).

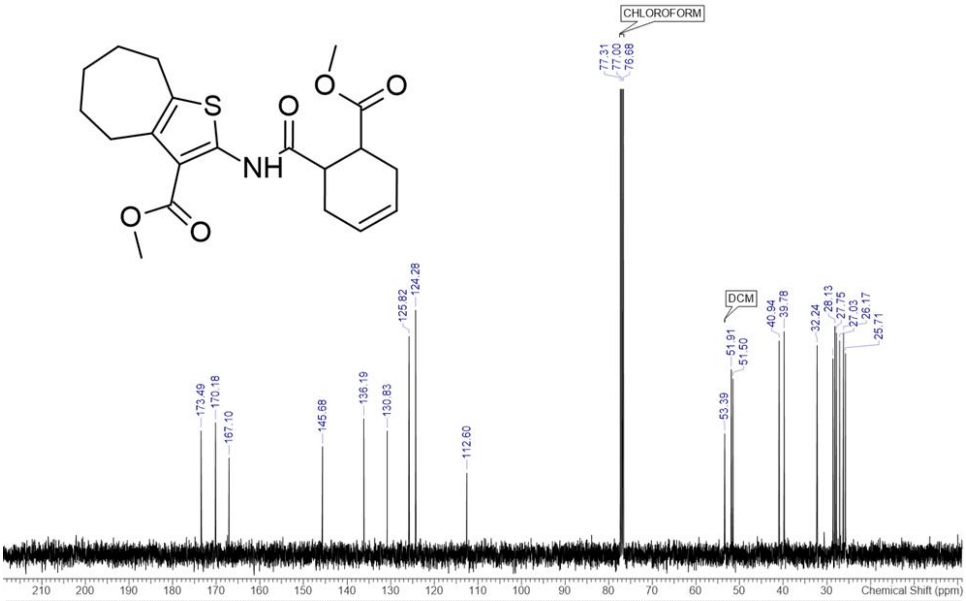


| No. | (ppm) | (Hz) | Height | No. | (ppm) | (Hz) | Height | No. | (ppm) | (Hz) | Height |
|---|---|---|---|---|---|---|---|---|---|---|---|
| 1 | 25.71 | 2587.1 | 0.2566 | 8 | 39.78 | 4002.2 | 0.2846 | 15 | 130.83 | 13163.3 | 0.1580 |
| 2 | 26.17 | 2633.3 | 0.2826 | 9 | 40.94 | 4119.6 | 0.2728 | 16 | 136.19 | 13702.5 | 0.1734 |
| 3 | 27.03 | 2719.1 | 0.2730 | 10 | 51.50 | 5181.8 | 0.2241 | 17 | 145.68 | 14657.7 | 0.1380 |
| 4 | 27.75 | 2792.5 | 0.2829 | 11 | 51.91 | 5222.9 | 0.2361 | 18 | 167.10 | 16812.2 | 0.1233 |
| 5 | 28.13 | 2830.6 | 0.2914 | 12 | 112.60 | 11328.6 | 0.1042 | 19 | 170.18 | 17122.5 | 0.1684 |
| 6 | 28.58 | 2875.4 | 0.2500 | 13 | 124.28 | 12503.8 | 0.3119 | 20 | 173.49 | 17455.6 | 0.1581 |
| 7 | 32.24 | 3243.6 | 0.2669 | 14 | 125.82 | 12659.3 | 0.2782 | | | | |

**Figure S26.** $^{1}$H NMR spectrum of compound **24** in $d_6$-DMSO (-0.5–15 ppm).

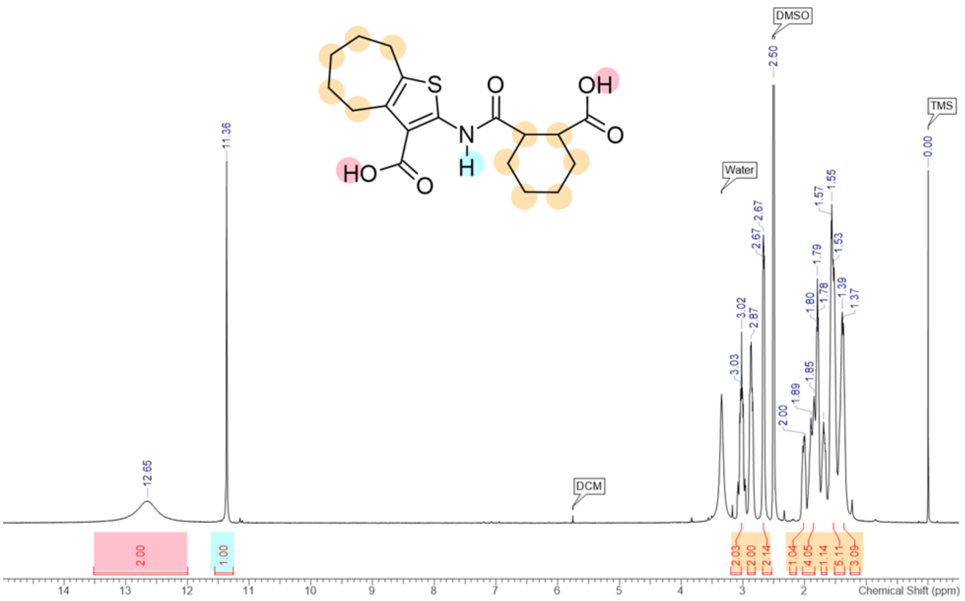


**Figure S27.** $^{13}$C NMR spectrum of compound **24** in $d_6$-DMSO (0–220 ppm).

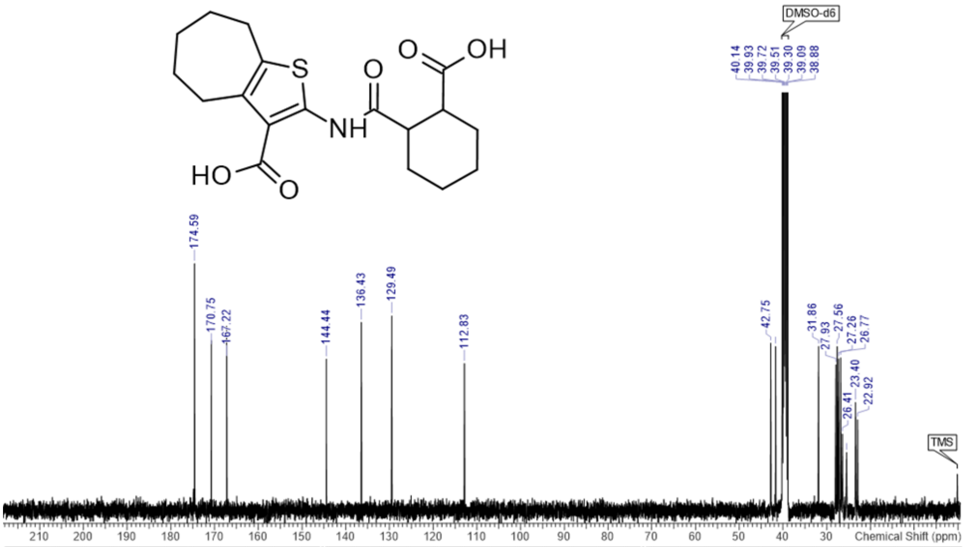


| No. | (ppm) | (Hz) | Height | No. | (ppm) | (Hz) | Height | No. | (ppm) | (Hz) | Height |
|---|---|---|---|---|---|---|---|---|---|---|---|
| 1 | 22.92 | 2305.5 | 0.0250 | 7 | 27.56 | 2772.8 | 0.0452 | 13 | 129.49 | 13028.5 | 0.0536 |
| 2 | 23.40 | 2354.0 | 0.0298 | 8 | 27.93 | 2810.3 | 0.0401 | 14 | 136.43 | 13726.2 | 0.0518 |
| 3 | 25.43 | 2558.6 | 0.0160 | 9 | 31.86 | 3205.7 | 0.0452 | 15 | 144.44 | 14532.4 | 0.0417 |
| 4 | 26.41 | 2657.7 | 0.0211 | 10 | 41.66 | 4191.6 | 0.0452 | 16 | 167.22 | 16824.1 | 0.0418 |
| 5 | 26.77 | 2693.6 | 0.0421 | 11 | 42.75 | 4301.7 | 0.0461 | 17 | 170.75 | 17179.9 | 0.0450 |
| 6 | 27.26 | 2742.8 | 0.0418 | 12 | 112.83 | 11352.2 | 0.0405 | 18 | 174.59 | 17565.8 | 0.0680 |

**Figure S28.** $^{1}$H NMR spectrum of compound **25** in $CDCl_3$ (-0.5–15 ppm).

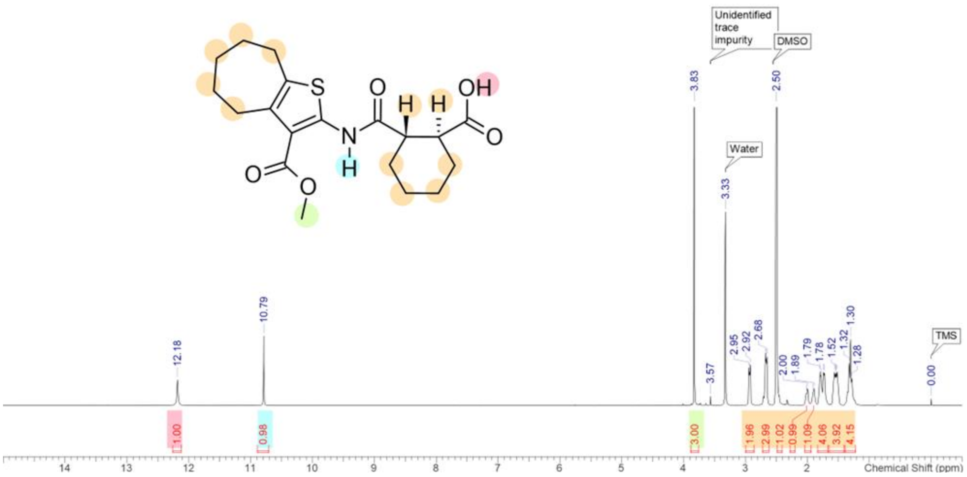


**Figure S29.** $^{13}$C NMR spectrum of compound **25** in $CDCl_3$ (0–220 ppm).

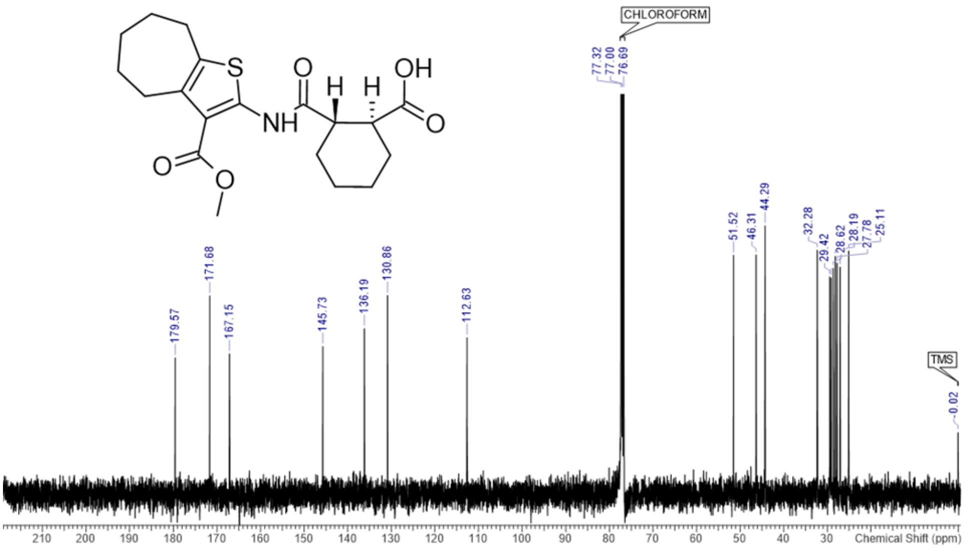


| No. | (ppm) | (Hz) | Height |
|---|---|---|---|
| 1 | 25.07 | 2522.5 | 0.0516 |
| 2 | 25.11 | 2526.2 | 0.0561 |
| 3 | 27.05 | 2722.1 | 0.0524 |
| 4 | 27.78 | 2795.4 | 0.0533 |
| 5 | 28.19 | 2836.5 | 0.0548 |
| 6 | 28.62 | 2879.0 | 0.0521 |
| 7 | 29.12 | 2929.7 | 0.0499 |
| 8 | 29.42 | 2960.5 | 0.0501 |
| 9 | 32.28 | 3247.3 | 0.0562 |
| 10 | 44.29 | 4456.3 | 0.0619 |
| 11 | 46.31 | 4659.5 | 0.0552 |
| 12 | 51.52 | 5184.0 | 0.0551 |
| 13 | 112.63 | 11332.3 | 0.0362 |
| 14 | 130.86 | 13166.3 | 0.0458 |
| 15 | 136.19 | 13702.5 | 0.0382 |
| 16 | 145.73 | 14662.8 | 0.0341 |
| 17 | 167.15 | 16817.4 | 0.0324 |
| 18 | 171.68 | 17273.7 | 0.0458 |
| 19 | 179.57 | 18067.4 | 0.0315 |

**Figure S30.** $^{1}H$ NMR spectrum of compound **26** in $d_6$-DMSO (-0.5–15 ppm).

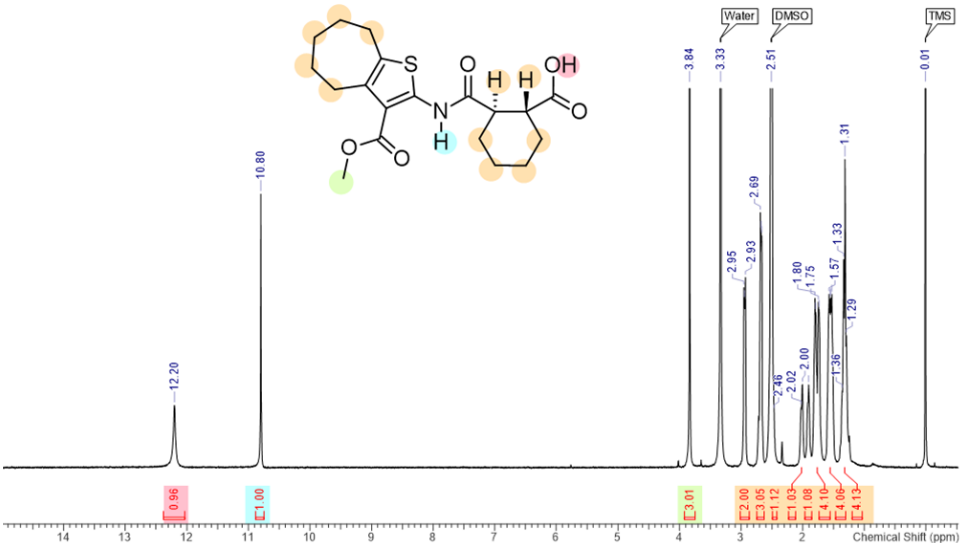


**Figure S31.** $^{13}C$ NMR spectrum of compound **26** in $CDCl_3$ (0–220 ppm).

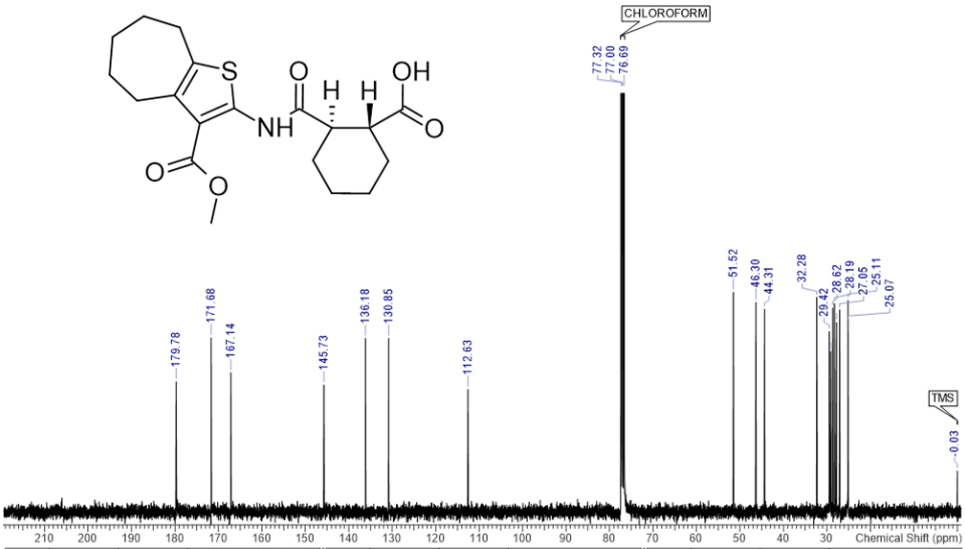


| No. | (ppm) | (Hz) | Height | No. | (ppm) | (Hz) | Height | No. | (ppm) | (Hz) | Height |
|---|---|---|---|---|---|---|---|---|---|---|---|
| 1 | 25.07 | 2522.5 | 0.0751 | 8 | 29.42 | 2960.5 | 0.0713 | 15 | 136.18 | 13701.8 | 0.0686 |
| 2 | 25.11 | 2526.2 | 0.0837 | 9 | 32.28 | 3247.3 | 0.0849 | 16 | 145.73 | 14662.8 | 0.0502 |
| 3 | 27.05 | 2722.1 | 0.0799 | 10 | 44.31 | 4458.5 | 0.0801 | 17 | 167.14 | 16816.6 | 0.0551 |
| 4 | 27.78 | 2795.4 | 0.0749 | 11 | 46.30 | 4658.0 | 0.0829 | 18 | 171.68 | 17273.7 | 0.0688 |
| 5 | 28.19 | 2836.5 | 0.0824 | 12 | 51.52 | 5183.3 | 0.0869 | 19 | 179.78 | 18088.0 | 0.0515 |
| 6 | 28.62 | 2879.0 | 0.0806 | 13 | 112.63 | 11331.5 | 0.0485 | | | | |
| 7 | 29.13 | 2930.4 | 0.0635 | 14 | 130.85 | 13165.5 | 0.0686 | | | | |

**Figure S32.** $^{1}$H NMR spectrum of compound **27** in $d_6$-DMSO (-0.5–15 ppm).

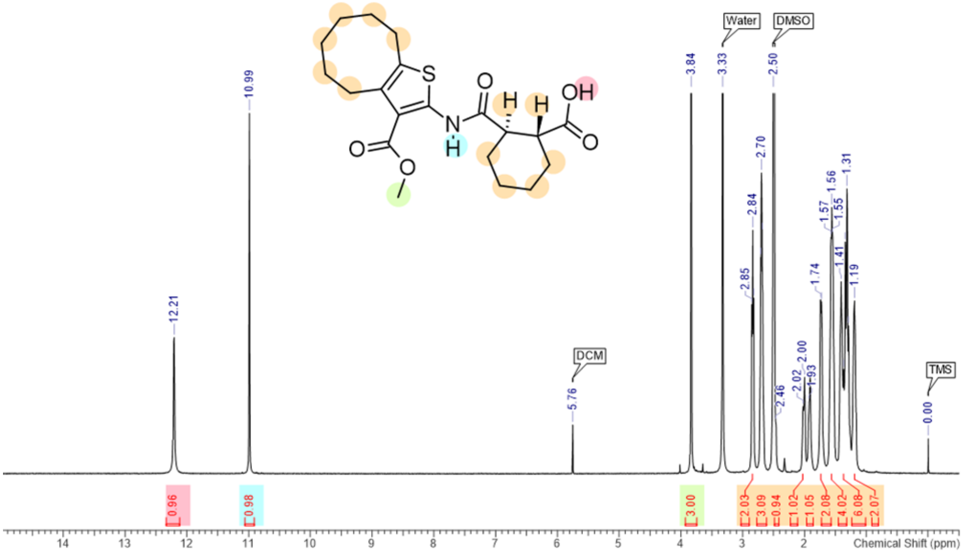


**Figure S33.** $^{13}$C NMR spectrum of compound **27** in $CDCl_3$ (0–220 ppm)

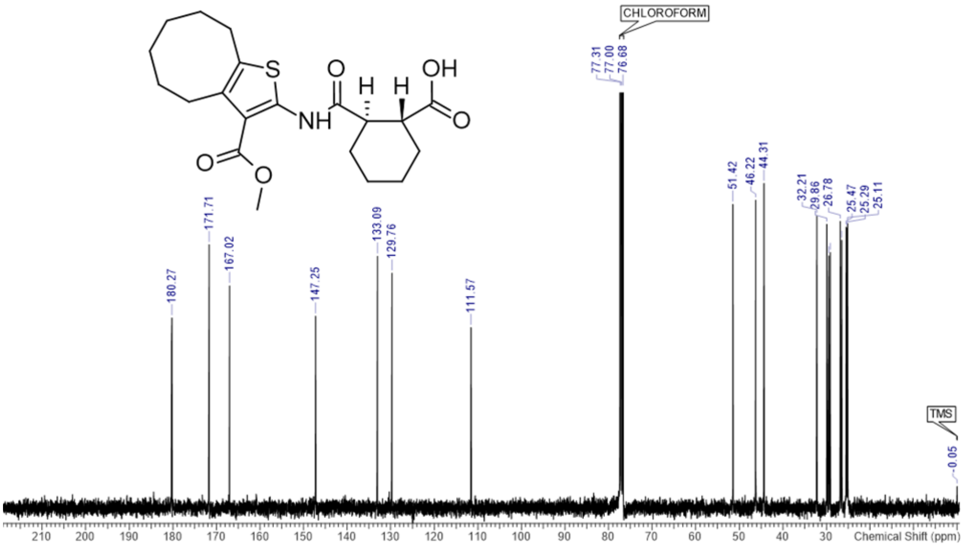


| No. | (ppm) | (Hz) | Height |
|---|---|---|---|
| 1 | 25.06 | 2521.8 | 0.1889 |
| 2 | 25.11 | 2526.2 | 0.1960 |
| 3 | 25.29 | 2544.5 | 0.1928 |
| 4 | 25.47 | 2562.1 | 0.1935 |
| 5 | 26.46 | 2661.9 | 0.1845 |
| 6 | 26.78 | 2694.9 | 0.1975 |
| 7 | 29.12 | 2929.7 | 0.1760 |
| 8 | 29.45 | 2962.7 | 0.1737 |
| 9 | 29.86 | 3004.5 | 0.1955 |
| 10 | 32.21 | 3240.7 | 0.2013 |
| 11 | 44.31 | 4457.7 | 0.2237 |
| 12 | 46.22 | 4649.9 | 0.2121 |
| 13 | 51.42 | 5173.7 | 0.2092 |
| 14 | 111.57 | 11225.2 | 0.1244 |
| 15 | 129.76 | 13055.5 | 0.1619 |
| 16 | 133.09 | 13390.7 | 0.1735 |
| 17 | 147.25 | 14815.4 | 0.1322 |
| 18 | 167.02 | 16804.2 | 0.1531 |
| 19 | 171.71 | 17276.6 | 0.1815 |
| 20 | 180.27 | 18137.8 | 0.1310 |

**Figure S34.** $^{1}$H NMR spectrum of compound **28** in $CDCl_3$ (-0.5–15 ppm).

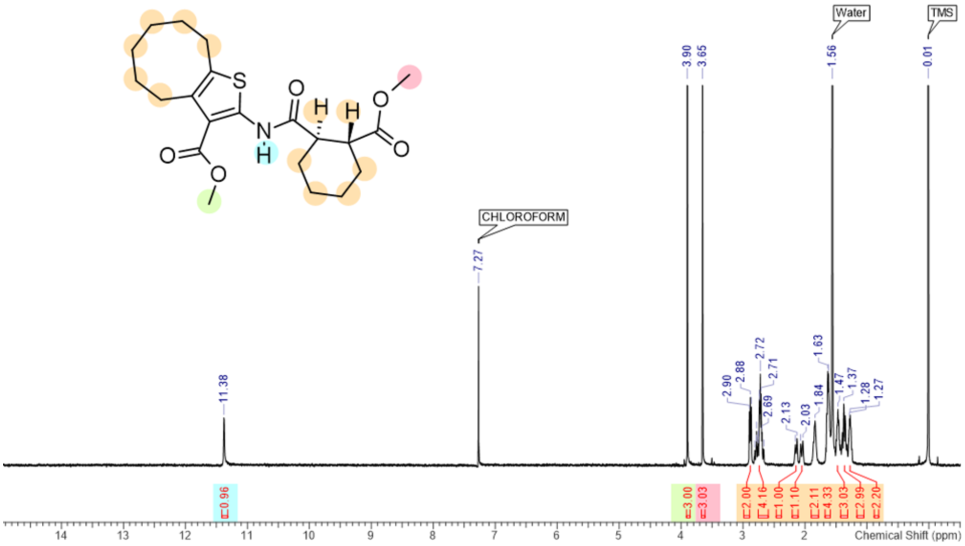


**Figure S35.** $^{13}$C NMR spectrum of compound **28** in $CDCl_3$ (0–220 ppm).

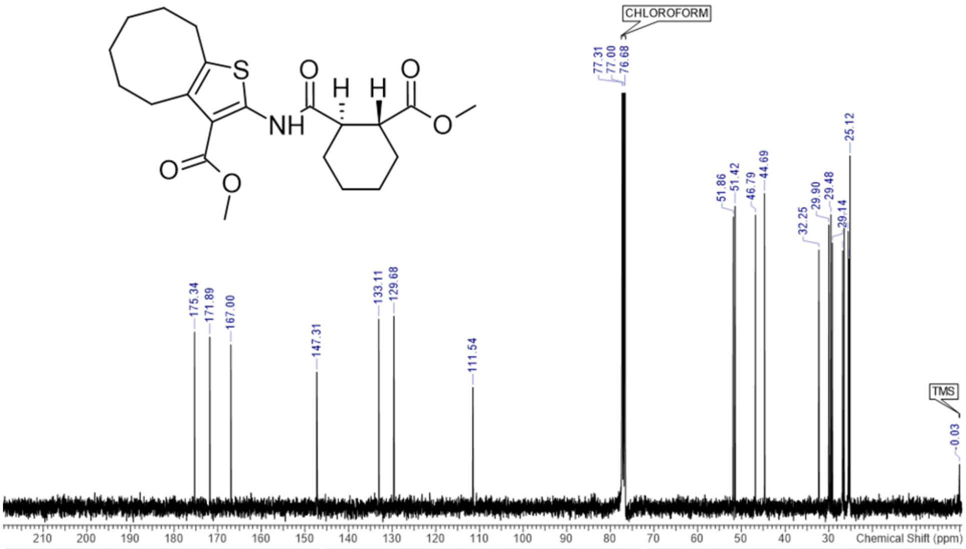


| No. | (ppm) | (Hz) | Height | No. | (ppm) | (Hz) | Height | No. | (ppm) | (Hz) | Height |
|---|---|---|---|---|---|---|---|---|---|---|---|
| 1 | 25.10 | 2525.5 | 0.1164 | 8 | 29.48 | 2965.6 | 0.1158 | 15 | 111.54 | 11222.2 | 0.0475 |
| 2 | 25.12 | 2527.7 | 0.1390 | 9 | 29.90 | 3008.2 | 0.1117 | 16 | 129.68 | 13047.4 | 0.0756 |
| 3 | 25.30 | 2546.0 | 0.0983 | 10 | 32.25 | 3245.1 | 0.1018 | 17 | 133.11 | 13392.9 | 0.0744 |
| 4 | 25.47 | 2562.1 | 0.1092 | 11 | 44.69 | 4495.9 | 0.1241 | 18 | 147.31 | 14821.2 | 0.0535 |
| 5 | 26.46 | 2661.9 | 0.1135 | 12 | 46.79 | 4707.9 | 0.1156 | 19 | 167.00 | 16802.7 | 0.0643 |
| 6 | 26.80 | 2696.4 | 0.1015 | 13 | 51.42 | 5173.7 | 0.1191 | 20 | 171.89 | 17294.2 | 0.0674 |
| 7 | 29.14 | 2931.9 | 0.1045 | 14 | 51.86 | 5217.7 | 0.1148 | 21 | 175.34 | 17641.9 | 0.0694 |

**Figure S36.** HPLC chromatogram of **27** at 220nm detection wavelength.

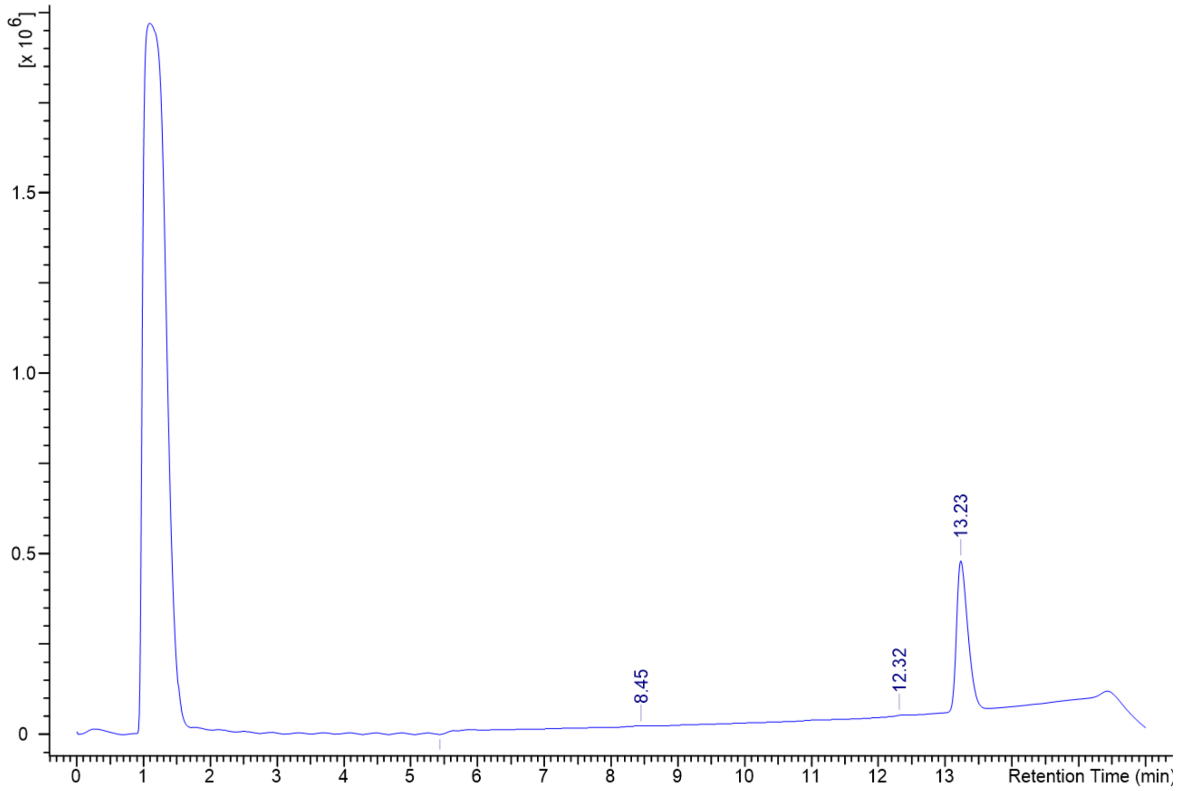


| No. | tR | Peak Area (Y units*ms) | Area Percent | Width |
|---|---|---|---|---|
| 1 | 8.45 | 36641216.000 | 0.739 | 0.650 |
| 2 | 12.32 | 16795072.000 | 0.339 | 0.179 |
| 3 | 13.23 | 4902276096.0( | 98.922 | 0.308 |

**Figure S37.** HPLC chromatogram of **27** at 260nm detection wavelength.

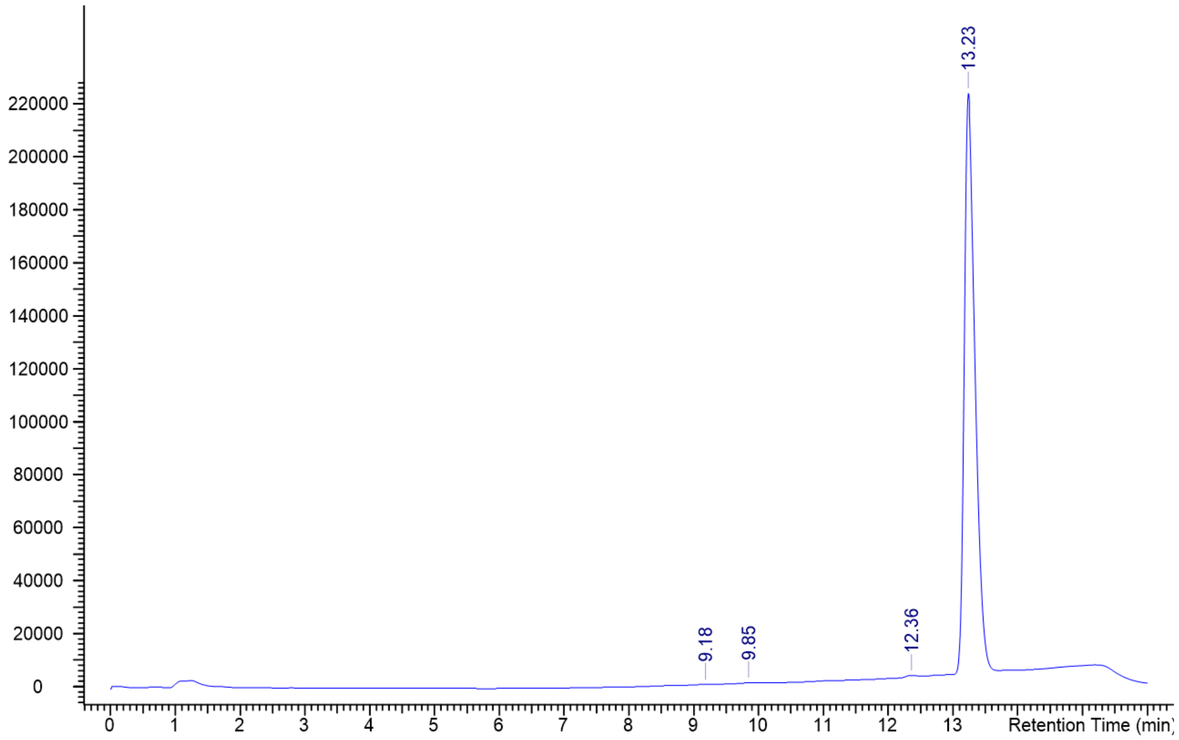


| No. | tR | Peak Area (Y units*ms) | Area Percent | Width |
|---|---|---|---|---|
| 1 | 9.18 | 4049237.000 | 0.156 | 0.456 |
| 2 | 9.85 | 7347586.500 | 0.283 | 0.790 |
| 3 | 12.36 | 6199013.500 | 0.238 | 0.287 |
| 4 | 13.23 | 2582017024.0[illegible] | 99.323 | 0.309 |
